# Innovation of heterochromatin functions drives rapid evolution of essential ZAD-ZNF genes in *Drosophila*

**DOI:** 10.1101/2020.07.08.192740

**Authors:** Bhavatharini Kasinathan, Serafin U. Colmenares, Hannah McConnell, Janet M. Young, Gary H. Karpen, Harmit S. Malik

**Affiliations:** Medical Scientist Training Program, University of Washington School of Medicine, Seattle, WA; Molecular and Cellular Biology Graduate program, University of Washington School of Medicine, Seattle, WA; Division of Basic Sciences, Fred Hutchinson Cancer Research Center, Seattle WA; Howard Hughes Medical Institute, Fred Hutchinson Cancer Research Center, Seattle WA; Biological Systems and Engineering Division, Lawrence Berkeley National Laboratory, Berkeley, CA; Department of Molecular and Cell Biology, University of California at Berkeley, Berkeley, CA; Innovative Genomics Institute, Berkeley, California

**Keywords:** positive selection, multigene family, gene duplication, gene loss, recombination, viability, fertility, heterochromatin

## Abstract

Contrary to prevailing dogma, evolutionarily young and dynamic genes can encode essential functions. Here, we investigate genetic innovation in *ZAD-ZNF* genes, which encode the most abundant class of insect transcription factors. We find that evolutionarily dynamic *ZAD-ZNF* genes are more likely to encode essential functions in *Drosophila melanogaster* than ancient, conserved *ZAD-ZNF* genes. To understand the basis of this unexpected correlation, we focus on the *Nicknack ZAD-ZNF* gene. *Nicknack* is an evolutionarily young, poorly retained in *Drosophila* species, and evolves under strong positive selection, yet we find that it is necessary for larval development in *D. melanogaster.* We show that *Nicknack* encodes a heterochromatin-localizing protein like its closely related paralog *Oddjob*, also an evolutionarily dynamic, essential *ZAD-ZNF* gene. We find that the divergent *D. simulans* Nicknack protein can still localize to *D. melanogaster* heterochromatin and rescue viability of female but not male *Nicknack*-null *D. melanogaster*. Our findings suggest that innovation for rapidly changing heterochromatin functions might provide a general explanation for the essential functions of many evolutionarily dynamic *ZAD-ZNF* genes in insects.

## Introduction

Although organisms display enormous phenotypic diversity, their cellular organization and early development are highly conserved across broad taxonomic ranges [1]. Such widespread conservation has led to a commonly held view that an ancient, conserved genetic architecture encodes fundamental biological functions, which appears to be largely borne out by comparative genomics [1, 2]. However, recent studies demonstrated that 30% of 185 evolutionarily young genes in *D. melanogaster* [3] have acquired roles in development, cell biology, and reproduction that render them essential for viability or fertility [4–6]. Sometimes, evolutionary turnover of genes underlying essential cellular processes can occur, as seen in the evolution of kinetochore proteins [7]. Even when they are retained over long evolutionary periods, genes encoding essential functions can evolve unexpectedly rapidly across plants and animals [8, 9]. Thus, at least a subset of essential functions is encoded by rapidly evolving genes or genes that are subject to high genetic turnover. The functional basis of this correlation, which runs counter to dogma, is unclear.

To study this unexpected class of rapidly evolving, essential genes further, we focused on the highly dynamic *ZAD-ZNF* gene family, which encodes several essential transcription factors. ZAD-ZNF proteins contain a conserved N-terminal ZAD (Zinc-finger associated domain), a linker, and a C-terminal domain that includes tandem C2H2 zinc fingers [10, 11]. The ZAD facilitates protein-protein interactions but does not have DNA-binding ability, whereas the C2H2 zinc fingers often mediate sequence-specific DNA binding [12]. Unlike the ZAD and ZNF domains, which are highly homologous between different ZAD-ZNF proteins, the linker domains are highly variable across ZAD-ZNF proteins in both sequence and length and have no discernible structural motifs. *ZAD-ZNF* genes arose in the ancestor of vertebrates and arthropods, but dramatically expanded within insect lineages [13], becoming the most abundant class of TFs in many genomes, including in *D. melanogaster* [10]. However, *ZAD-ZNF* gene repertoires can vary quite extensively across insect lineages via gene gains and losses [10, 13]; the causes and consequences of this gene dynamism are poorly understood.

Insect ZAD-ZNF proteins might act analogously to mammalian KRAB zinc finger (KZNFs), providing an explanation for their gene dynamism [10, 13]. *KZNF* genes arose in the ancestor of tetrapods and expanded through lineage-specific gene amplifications to become the most abundant class of transcription factors present in mammalian genomes [14, 15]. KZNF proteins contain an N-terminal KRAB (Krüppel-associated box) domain and C-terminal arrays of C2H2 zinc finger domains that define their DNA-binding specificities. Many KZNF genes play a critical role in genome defense through recognition and repression of transposable elements [16, 17]. As a result, KZNF genes involved in genome defense are subject to rapid genetic turnover and positive selection of their DNA-binding domains [18]. Similar functions might have driven the diversification of ZAD-ZNF genes in insects [10, 13].

Despite their abundance, only a few *ZAD-ZNFs* have been functionally characterized, with most studies restricted to *D. melanogaster*. Approximately half of all *ZAD-ZNF* genes are highly expressed in ovaries and early embryos [10], where they might play crucial roles for fertility or development. Biochemical characterization of 21 ZAD-ZNF proteins shows that they bind to unique DNA consensus sequences, with putative targets in the regulatory regions of specific target genes [19]. For example, the ZAD-ZNF protein M1BP (Motif 1 Binding Protein) binds core promoters and promotes the expression of numerous housekeeping genes [20, 21]. Similarly, *ZAD-ZNF* gene *Grauzone* promotes *cortex* expression and is necessary for meiotic progression during oogenesis [22–24]. Finally *ZAD-ZNF* genes *Molting Defective, Ouija Board,* and *Séance* encode proteins that promote the transcription of heterochromatin-embedded genes, *Spookier* and *Neverland,* required for larval progression [25]. In contrast, other ZAD-ZNF proteins repress rather than drive transcription. For example, the ZAD-ZNF protein Oddjob localizes to heterochromatin and can lead to position-effect variegation [26]. Another ZAD-ZNF protein encoded by *CG17801* helps repress *HetA* and *Blood* transposable elements in the ovary [27].

Not all ZAD-ZNF functions are directly related to transcription. For example, the ZAD-ZNF proteins ZIPIC, Zw5, and Pita help organize chromatin architecture [28, 29] whereas the ZAD-ZNF protein Trade Embargo mediates the initiation of meiotic recombination during oogenesis [30]. Finally, some ZAD-ZNFs might not function in the nucleus at all. Even though the Weckle ZAD-ZNF protein possesses zinc finger domains, it localizes to the plasma membrane instead of nuclear chromatin, where it interacts with the Toll-MyD88 complex to help establish the anterior-posterior axis of the developing embryo [31].

The *ZAD-ZNF* gene family in *Drosophila* is ideal for studying the relationship between genetic innovation and essentiality because of their involvement with essential cellular processes despite rapid evolutionary dynamics. In order to study this relationship between rapid evolution and essentiality, we leveraged the extensive opportunities for phylogenomic and population genetics studies in *Drosophila* species. We also took advantage of genome-wide screens for phenotype and tools for cytological and genetic analyses in *D. melanogaster*. Using evolutionary analyses, we identified the *D. melanogaster ZAD-ZNF* genes that had been either subject to genetic turnover or positive selection. Although only a few *ZAD-ZNF* genes have undergone positive selection, we found that these genes are more likely to be required for viability or fertility in *D. melanogaster* than slowly-evolving *ZAD-ZNF* genes. We focused on the characterization of one of these positively-selected *ZAD-ZNF* genes: *Nicknack (CG17802, Nnk).* We show that *Nicknack* is essential for larval development in *D. melanogaster* despite being evolutionarily young and differentially retained among *Drosophila* species. *Nicknack* belongs to a small cluster of *ZAD-ZNF* paralogs, the best characterized of which is *Oddjob (CG7357, Odj*). We show that both *Odj* and *Nnk* encode heterochromatin-localizing proteins. Although Odj broadly localizes to heterochromatin, we found that Nnk predominantly localizes to discrete foci within heterochromatin. Surprisingly, despite a strong signature of positive selection, we found that the protein encoded by the divergent *D. simulans Nicknack* ortholog can still localize to heterochromatin in *D. melanogaster* cells. Furthermore, *D. simulans Nicknack* can significantly rescue the viability of *D. melanogaster Nicknack-null* females but is unable to rescue *Nicknack-null* males. Based on our functional and cytological analyses, we conclude that rapidly changing requirements for heterochromatin function likely drove essential innovation of *ZAD-ZNF* genes such as *Nicknack* and *Oddjob* in *Drosophila*.

## Results

### *ZAD-ZNFs* genes are dynamic and diverse in *Drosophila*

We searched Flybase to identify all genes in *D. melanogaster* that encode a ZAD domain (PF07776; Pfam database, Pfam.org). We found 91 *ZAD-ZNF* genes distributed across Chromosomes 2, 3, and X. 37 *ZAD-ZNF* genes occur in 13 gene clusters, containing two or more tandemly-arrayed *ZAD-ZNF* genes. Of these, many *ZAD-ZNF* genes share intron/exon structures with their neighbors and likely arose via segmental duplication. In contrast, seven *ZAD-ZNF* genes (*CG3032, CG4318, CG9215, CG44002, CG17361, CG7963, CG17359*) lack introns found in their closest relatives; we infer that these genes were likely born via retrotransposition.

Although ZAD-ZNF proteins are defined as having both a ZAD and a ZNF domain, further analysis of ZAD-containing proteins using NCBI’s Conserved Domain Database revealed significant variation in the ZNF domains. We found that ZAD-containing proteins have an average of ~6 C2H2 domains. However, some ZAD-containing proteins have no C2H2 domains at all (*dlip, dbr, CG15435, CG31109, CG31457*), whereas others contain up to 23 C2H2 domains (*CG11902*). In addition to the ZAD in the N-terminus, 13 ZAD-ZNF proteins possess another N-terminal domain, such as a Sodium-Calcium exchanger domain (*CG12391*) or ASF1 histone chaperone domain (*CG10321*). The functional significance of these additional domains is unknown.

Next, we surveyed *ZAD-ZNF* genes in all 12 previously sequenced and annotated *Drosophila* genomes. These species represent a range of evolutionary divergence from *D. melanogaster*, from just a few million years (*e.g., D. simulans*), to more than 40 million years (*e.g., D. virilis*) [32]. Our analysis reveals a surprisingly wide range in number of *ZAD-ZNF* genes across different *Drosophila* species genomes (Figure 1). For example, we found that the *D. persimilis* genome encodes 130 *ZAD-ZNF* genes whereas the *D. willistoni* genome encodes only 75 *ZAD-ZNF* genes. Such analyses are dependent on the state of completion and annotation of individual *Drosophila* species’ genomes. Thus, this number may be an underestimate. To complement these analyses, we assessed the apparent age of each *D. melanogaster ZAD-ZNF* gene by examining the presence of orthologs in the 12 annotated genomes of *Drosophila* species (flybase.org). We found that 73 of 91 *D. melanogaster ZAD-ZNF* genes arose in or prior to the common ancestor of all *Drosophila* species, 40 million years ago. Of these 73 *ZAD-ZNF* genes, 61 have been preserved over *Drosophila* evolution, whereas 12 have been lost in at least one lineage or species. We estimate that 17 *ZAD-ZNF* genes arose via gene duplication after the last common ancestor of *D. melanogaster* and *D. virilis.* At least 3 *ZAD-ZNF* genes found in *D. melanogaster (CG4318, CG17612, neu2)* originated via gene duplication less than 10 million years ago. Our findings complement previous large-scale surveys that identified rapid changes in *ZAD-ZNF* gene repertoires within insect genomes [10, 11, 13].

**Figure 1:**
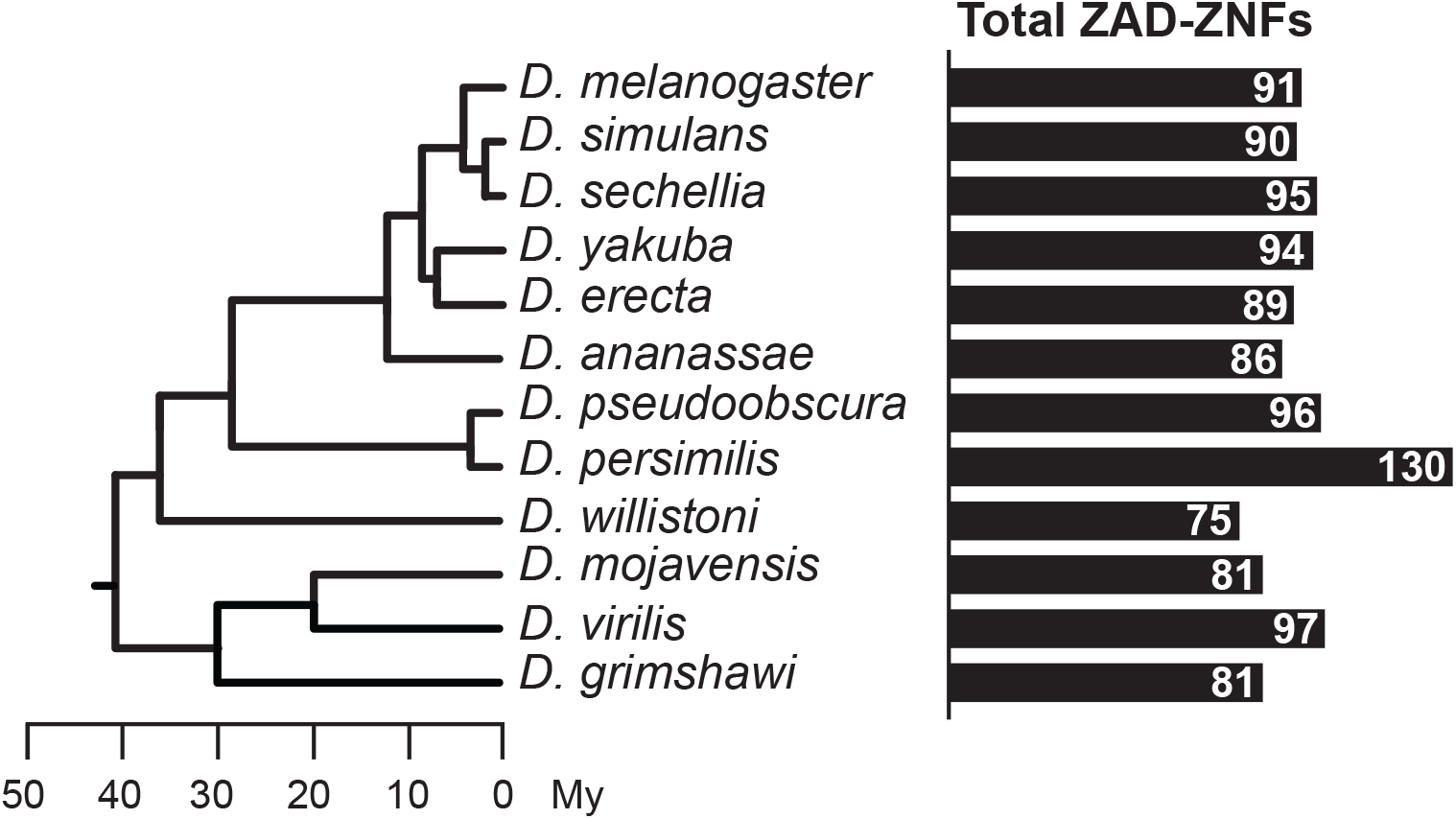
Total number of ZAD-ZNFs across *Drosophila*. Phylogeny of 12 *Drosophila* genomes with a scale bar showing approximate divergence times [32]. The number of ZAD-containing genes in each genome indicated by black bars.

### Rapidly evolving *ZAD-ZNF* genes are frequently essential in *D. melanogaster*

These rapid changes in *ZAD-ZNF* gene repertoires suggested that selection might favor their genetic innovation. We investigated whether evolutionary retention is a predictor of essentiality. We took advantage of the fact that knockdown or knockout phenotypes have been characterized for almost all *D. melanogaster* genes. Indeed, 86 of 91 *D. melanogaster ZAD-ZNF* genes have associated phenotypic outcomes (flybase.org). Of these, knockdown or knockout of 28 *ZAD-ZNF* genes showed complete lethality or sterility in *D. melanogaster*, whereas the other 58 were determined not to be essential.

Of the 62 *ZAD-ZNF* genes with orthologs retained in all 12 annotated *Drosophila* species’ genomes, we found that 17 are essential for either fertility or viability in *D. melanogaster*, whereas 41 genes are not (4 genes have no phenotypic data available). In comparison, we found that 11 of the 29 genes not universally conserved in *Drosophila* species are essential, whereas 17 are not (1 has no phenotypic data available). Thus, somewhat surprisingly, we find that genes not globally retained over *Drosophila* evolution are just as likely to encode a necessary function in *D. melanogaster* as genes that have been strictly maintained over 40 million years of *Drosophila* evolution (11:17 versus 17:41, p>0.4, chi-squared test) (Table 1A). These findings further support the idea that gene families subject to rapid evolutionary turnover may become involved in essential functions.

**Table 1.**
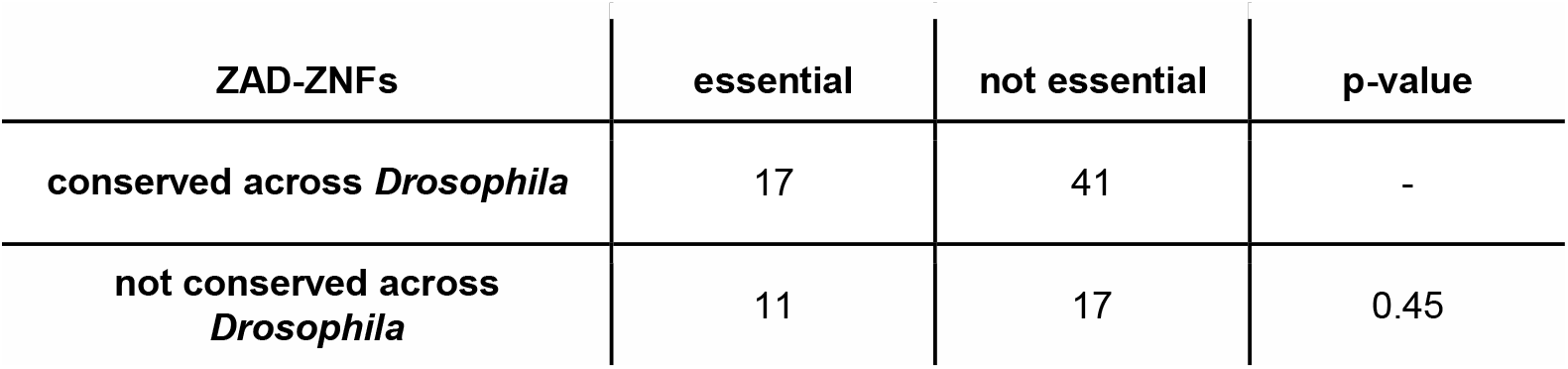
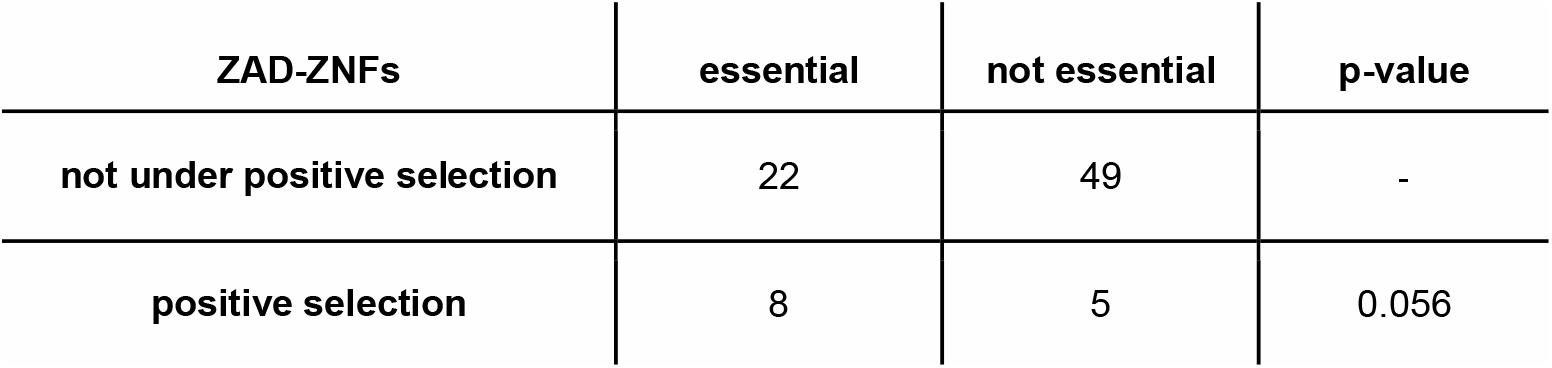
Links between *ZAD-ZNF* gene dynamism and essentiality. **(A)** ZAD-ZNF genes are just as likely to be essential whether or not they are conserved across all 12 *Drosophila* species. Note that 5 genes for which data is not available were not included in these analyses. p-values were calculated by two-tailed Fisher’s exact test. **(B)** Positively-selected ZAD-ZNFs (via McDonald-Kreitman test) are more likely to be essential than ZAD-ZNFs that have not evolved under positive selection. *D. simulans* lacks one of the 91 *D. melanogaster* genes and no data on essentiality is available for 6 genes. p-values were calculated by two-tailed Fisher’s exact test.

Gene duplication and loss is only one form of genetic innovation. We also analyzed the *D. melanogaster ZAD-ZNF* genes for signatures of recent positive selection. We performed McDonald-Kreitman tests of *ZAD-ZNF* orthologs found in both *D. melanogaster* and the closely related species, *D. simulans*. The McDonald-Kreitman test assesses sequence diversity within a species versus between species by comparing the ratio of non-synonymous (amino-acid altering, or replacement) to synonymous substitutions fixed during the divergence of the two species (Dn: Ds), to that of non-synonymous to synonymous polymorphisms within a species (Pn: Ps) [33]. The Pn: Ps ratio is a proxy for functional constraint acting on a gene within species and is expected to be similar to Dn: Ds between species under the null hypothesis. However, a higher than expected number of fixed non-synonymous changes would indicate the action of adaptive evolution during species divergence [33].

We took advantage of previous efforts that sequenced the genomes of hundreds of *D. melanogaster* strains and a few *D. simulans* strains [34–36] to perform the McDonald-Kreitman test using the Popfly server (popfly.uab.cat) [37]. Of the 91 *D. melanogaster ZAD-ZNF* genes, only *CG2202* is absent in *D. simulans*. We found that 13 out of the remaining 90 *ZAD-ZNF*s show evidence for recent adaptive evolution, *i.e.,* have an excess of fixed non-synonymous changes (Dn) (Table 2). Subsequently, we used a domain-restricted analysis to define which of the three protein domains (ZAD, linker, or ZNF) were subject to positive selection. In five cases, we were unable to define the domain subject to positive selection, due to small numbers of intra-species polymorphisms. One gene (*CG7386*) showed signatures of adaptive evolution in its ZAD domain, whereas three genes (*CG2712, CG7386, CG10321*) showed evidence of adaptive evolution in the ZNF domain. However, we found that the linker domain has evolved under positive selection in six *ZAD-ZNF* genes. The biochemical function of these linker domains are largely unexplored as they lack predicted structural motifs [10, 13], are highly variable in length and sequence even between orthologs, and are predicted to encode intrinsically disordered domains [38]. Our findings implicate the poorly characterized linker region in mediating the adaptive potential of several *ZAD-ZNF* genes.

**Table 2:**
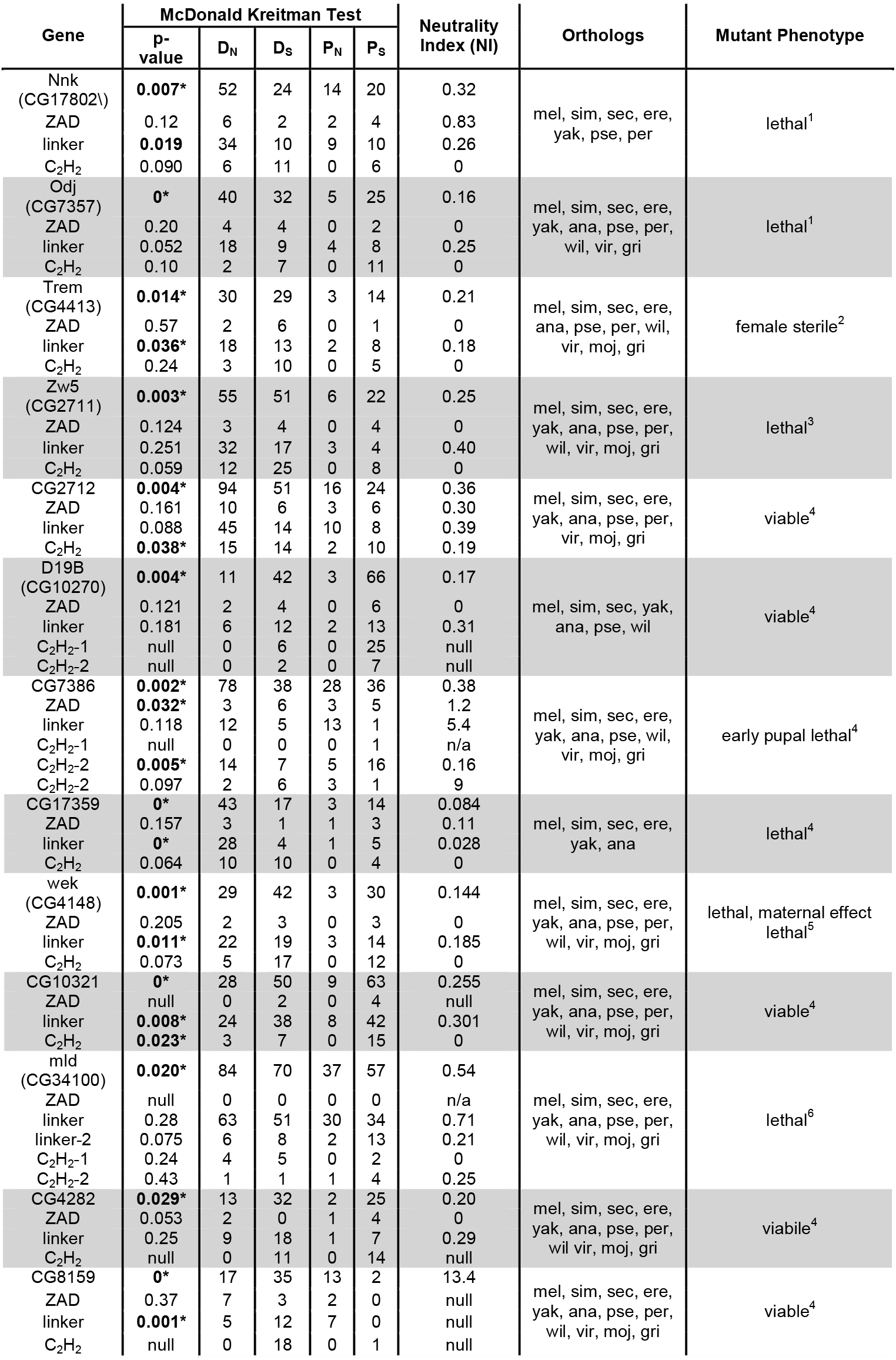
Positively selected *ZAD-ZNF*s. Summary statistics for the McDonald-Kreitman (MK) test with statistically significant values (p<0.05) denoted with an asterisk. The number of polymorphisms within *D. melanogaster* at synonymous (Ps) and non-synonymous (Pn) sites are compared to the number of fixed synonymous (Ds) and non-synonymous (Dn) changes between *D. melanogaster* and *D. simulans*. A neutrality index (N.I.) of <1 suggests an excess of fixed non-synonymous changes between species (i.e. positive selection). Orthologs column indicates species that contain an ortholog as determined by OrthoDB. The phenotype column reports phenotypes as follows: ^1^this study, ^2^ [30], ^3^ [28], ^4^ FlyBase.org, ^5^ [31], ^6^ [45].

Using this signature of positive selection in a subset of *ZAD-ZNF* genes, we evaluated whether positively selected genes encode essential functions. Intriguingly, we found that 8 of 13 *ZAD-ZNF* genes that have evolved under positive selection are essential, whereas only 22 of the remaining 71 genes that have been phenotypically assayed are essential. Thus, contrary to the dogma, we find that *ZAD-ZNF* genes that are subject to positive selection are more likely to be essential for viability or fertility (8:5 versus 22:49, Fisher’s exact test, p=0.056) (Table 1B). Our findings not only imply that several essential *ZAD-ZNF* genes evolve rapidly, but also raise the possibility that rapid evolution of some *ZAD-ZNF* genes might be critical for organismal viability.

### The *Oddjob-Nicknack* cluster of *ZAD-ZNF* genes evolves dynamically in *Drosophila*

To further investigate the biological basis for the correlation between positive selection and gene essentiality, we decided to focus on one cluster of *ZAD-ZNF* genes on chromosome 3 of *D. melanogaster* (Figure 2A). This cluster of five genes contains two of the eight positively-selected, essential genes in *D. melanogaster*: *Oddjob (Odj, CG7357)* and *CG17802*. *Odj* is the best-characterized gene in this cluster. The Oddjob protein interacts with heterochromatin protein HP1a, dynamically localizes to heterochromatin, and is required for heterochromatin-mediated gene silencing, or position-effect variegation [26]. Knockdown or over-expression of *Odj* is lethal in *D. melanogaster*, although the functional basis for this lethality is still unknown [39]. In keeping with the *Oddjob* nomenclature theme of ‘James Bond henchmen,’ and the mutant phenotype (described below), we renamed *CG17802* as *Nicknack,* or *Nnk.* This cluster of *ZAD-ZNF* genes also includes three genes that do not evolve under positive selection *(CG17801, CG17806, and CG17803)* in *D. melanogaster* (Figure 2A). Although previous studies found that the knockdown of *CG17801* in the germline led to de-repression of HetA and Blood transposable elements in the ovary, *CG17801* knockdown did not significantly impair fertility or viability [27]. The other two *ZAD-ZNF* genes in this cluster (*CG17801, CG17803*) have not been previously investigated.

**Figure 2:**
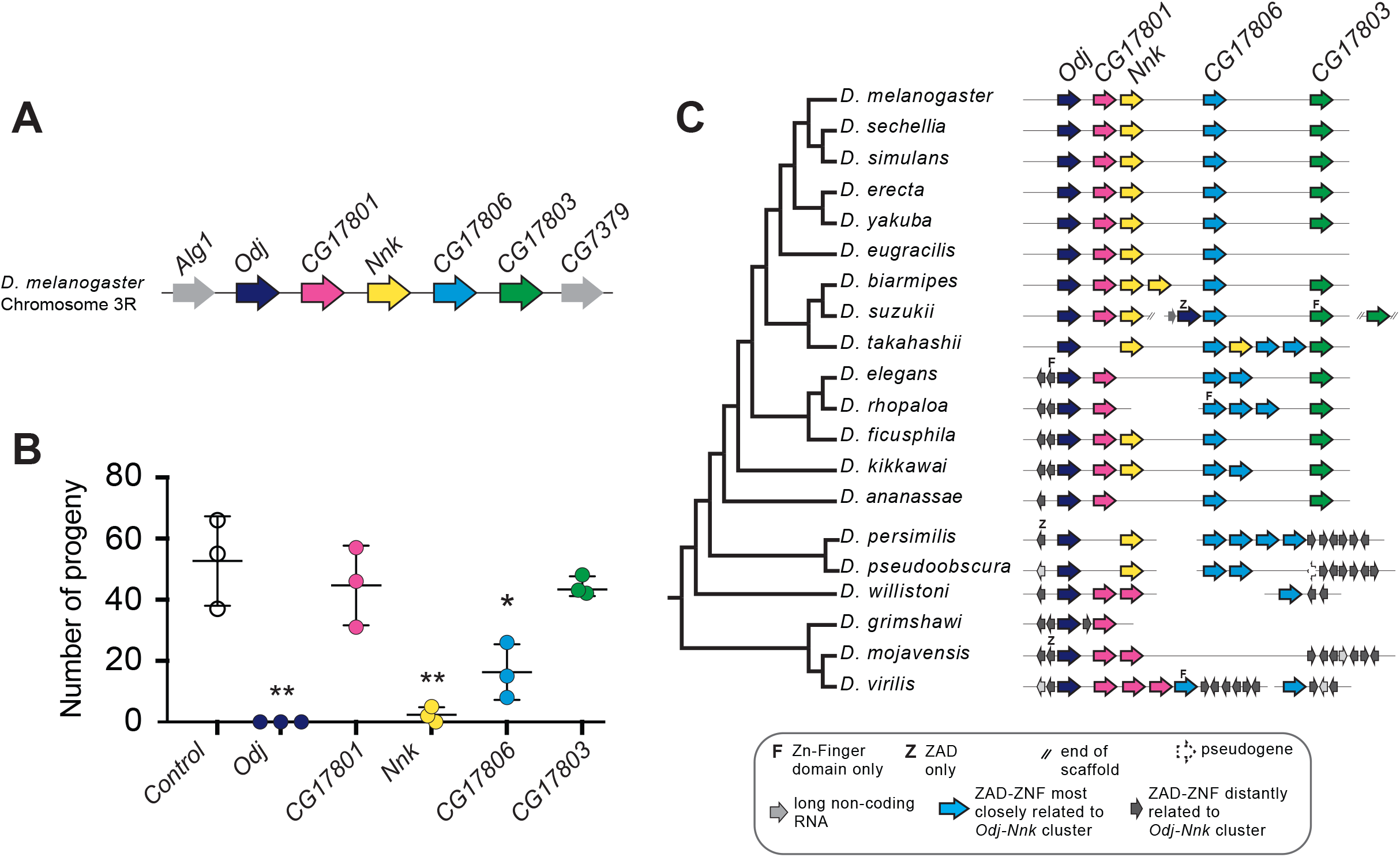
*Nicknack* is essential for viability. **(A)** A schematic of the *Oddjob-Nicknack* cluster of *ZAD-ZNF* genes in *D. melanogaster*. **(B)** Viability of adult flies ubiquitously knocked down for each *Odj-Nnk* cluster member. The vertical axis shows number of knockdown progeny per cross; each cross is represented by a point. Ubiquitous knockdown of either *Odj* or *Nnk* greatly reduces adult viability, whereas knockdown of *CG17806* has a mild but statistically significant effect on viability. Horizontal bars represent the mean and one standard deviation. We compared controls and ZAD-ZNF knockdowns using a two-tailed Student’s t-test; ** denotes p-value < 0.01, * denotes p-value <0.05. **(C)** Phylogenomic analysis of the *Odj-Nnk* cluster shows that *Oddjob*, *CG17801* and *CG17806* all date back to the origin of the *Drosophila* genus. However, *Oddjob* is the only gene within the cluster retained in all queried *Drosophila* species; *CG17801* has been lost within the *obscura* clade and in *D. takahashii,* whereas *CG17806* has undergone numerous gains and losses. *CG17803* arose in the ancestor of *D. melanogaster* and *D. ananassae* but was lost in *D. eugracilis. Nicknack* arose in the ancestor of *D. melanogaster* and *D. pseudoobscura,* duplicated in *D. biarmipes* and *D. takahashii,* and was lost in *D. ananassae* and the *D. elegans/D. rhopaloa* lineage.

We evaluated the functional roles of all five *ZAD-ZNF* genes in the *Odj-Nnk* cluster by generating knockdowns using an Actin5C-Gal4 driver and RNAi constructs specific to each gene (Figure 2B) to generate ubiquitous knockdown of individual genes in the resulting zygotes. Consistent with previous results [39], we found that the ubiquitous knockdown of *Odj* or *Nnk* is lethal. In contrast, we found that knockdowns of *CG17801,* and *CG17803* are viable, while knockdown of *CG17806* had significantly reduced viability (Figure 2B). Based on these results, we conclude that the two positively-selected members of this *ZAD-ZNF* cluster – *Odj* and *Nnk* – are both essential, confirming the unexpected correlation we previously observed between positive selection and gene essentiality in the *ZAD-ZNF* gene family (Table 1B).

To gain deeper insight into the evolutionary dynamics of the *Odj-Nnk* cluster, we identified orthologs of these genes using reciprocal TBLASTN searches with each of the five *D. melanogaster* genes as queries. We searched both the originally-sequenced, well-annotated 12 *Drosophila* genomes [32], as well as eight additional genomes that were subsequently sequenced to sample the melanogaster group within the *Sophophora* subgenus more densely [40]. In several cases, we were not able to confidently assign the TBLASTN hits to orthologous groups because they matched closely to more than one *D. melanogaster* gene, or there were several putative hits within a single genome, or because the hit contained only the ZAD or ZNF domain.

Using a multiple alignment of the ZNF domain (Supplementary Data S1), we conducted phylogenetic analyses (Supplementary Figure S1). Our phylogenomic analyses reveal that *Odj-Nnk* cluster evolution was highly dynamic during the evolution of the *Drosophila* genus (summarized in Figure 2C). Although *Oddjob* orthologs are present throughout 40 million years of *Drosophila* evolution, no other member of the *D. melanogaster Odj-Nnk* cluster is universally present in *Drosophila* species. In addition to *Odj*, *CG17801* and *CG17806* also date back prior to the origin of *Drosophila* but have since been lost in some species (Figure 2C). However, *CG17801* has been lost in the *obscura* group whereas *CG17806* underwent multiple independent duplication and loss events. *CG17803* arose in the ancestor of *D. melanogaster* and *D. ananassae* but underwent two independent losses. Finally*, Nnk* appears to have arisen in the ancestor of *D. melanogaster* and *D. pseudoobscura* (~35 mya, timetree.org), but later experienced multiple independent duplications and losses (Figure 2C). We note that our estimates for the age of *Nnk* are higher than those reported previously [3], likely because the prior estimate was based on fewer sequenced species. Based on these analyses, we conclude that, despite its essential function in *D. melanogaster, Nnk* is not universally conserved in *Drosophila* species.

### *Nicknack* is an essential *ZAD-ZNF* gene in *D. melanogaster*

*Nnk* has a dramatic evolutionary history: young evolutionary age, differential retention, positive selection. Yet, it serves an essential function in *D. melanogaster* based on our and other previous analyses. However, this claim of essentiality has been challenged by two previous findings. First, the screen that originally identified *Nnk* as an essential gene used the KK RNA-interference (RNAi) collection from the Vienna *Drosophila* Stock Center (VDRC) (Figure 3A) [3]. Many lines in this ‘KK’ collection were later found to harbor a second-site mutation that caused lethality as a result of ectopic *tiptop* expression when crossed to *GAL4-*driver lines [41, 42]. As a result of its dependence on the KK lines, the claim of *Nnk* essentiality remained ambiguous. A second *Nnk* mutant allele, created by CRISPR-Cas9-mediated mutagenesis (hereafter referred to as *CRISPR-null*), had a four base pair deletion within the coding sequence of the gene that created a frameshift and a premature stop codon within the linker domain of *Nnk* [43] (Figure 3A). Although this deletion was homozygous lethal, it was unexpectedly viable when paired with a deficiency covering this *ZAD-ZNF* cluster [43], again challenging the result that *Nnk* is an essential gene.

**Figure 3:**
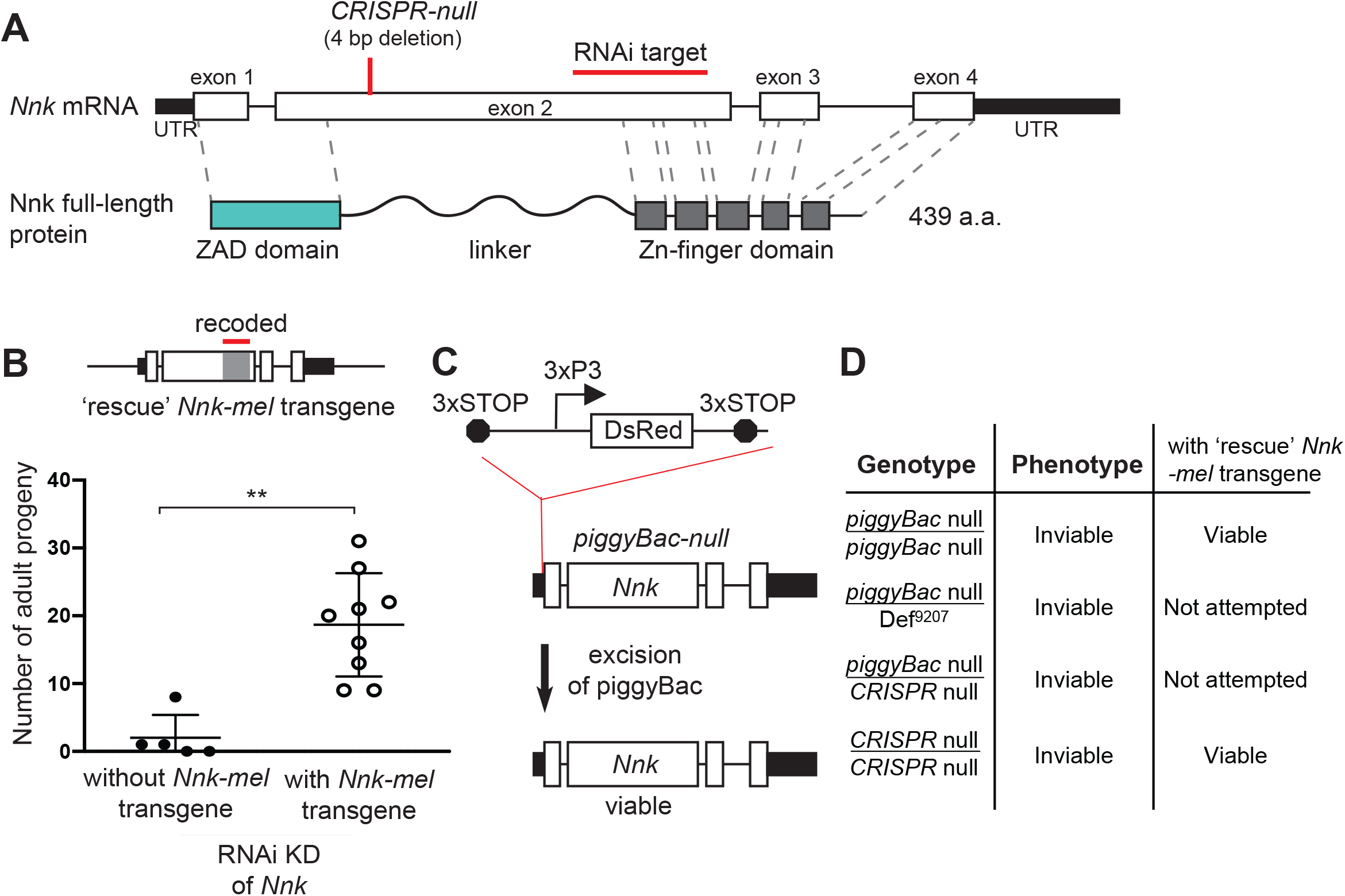
*Nicknack* RNAi and mutant alleles. **(A)** Schematic of *Nnk* gene. The *Nnk* gene contains four coding exons and encodes a full-length protein containing a ZAD, linker, and ZNF domains. The CRISPR-null is a 4 bp deletion within exon 2 [43]: this deletion creates a frameshift and a premature stop codon within the linker region. The VDRC RNAi KK line creates double-stranded RNA corresponding to a region of exon 2 (schematized). **(B)** Schematic of *Nnk-rescue* transgene design, which contains the genomic region 1 kb upstream of the *Nnk* start codon and 1 kb downstream of the stop codon. The region targeted by the RNAi construct (highlighted) was recoded by synonymous mutations to make it resistant to RNAi. Upon ubiquitous knockdown of *Nnk* using Act5c-GAL4 driven RNAi, adult viability is very low, but is significantly restored by the RNAi-resistant *Nnk*-rescue transgene. Horizontal bars represent the mean and error bars represent one standard deviation of replicates of single-pair matings for each cross. Asterisks indicate p<0.01, two-tailed Student’s t-test. **(C)** *piggyBac-*null is a *piggyBac* insertion in the 5’ UTR of *Nnk* that contains a fluorescent reporter (DsRed) driven by an eye specific promoter (3xP3) and flanked by stop codons in all three reading frames which terminate translation through *Nnk* [44]. Excision of this *piggyBac* insertion restores viability. (**D**) Allelic combinations of CRISPR/*piggyBac* nulls/genetic deficiencies were tested, in some cases in the presence of the *Nnk*-rescue transgene, to investigate and confirm *Nnk* essentiality (Supplementary Figure 3).

Given these ambiguous results and the importance of *Nnk* for our claims of *ZAD-ZNF* essentiality despite evolutionary innovation, we re-investigated whether *Nnk* is essential for viability in *D. melanogaster.* We found several lines of evidence to support the conclusion that it is indeed essential (Figure 3B-D). First, we began by validating the RNAi line used in the original study [3], which first identified *Nnk* as a young, essential gene. We found that the *Nnk* VDRC RNAi line does not have an insertion upstream of the *tiptop* gene, which is associated with lethality in other KK lines [41, 42]. Second, we were able to rescue this lethality via complementation with an intact *Nnk* transgene that has been recoded via mutations in synonymous sites to make it resistant to RNAi knockdown (Figure 3B) (Supplementary Data S2). Endogenous levels of *Nnk* expression are too low for us to validate expression of the transgene. However, our ability to rescue *Nnk* knockdown-mediated inviability strongly imply that the recoded transgene is appropriately expressed (Figure 3B).

To rule out any indirect effects arising from RNAi knockdown, we also examined two previously generated mutant alleles of *Nnk*. The first is a *piggyBac* insertion two bp upstream of the start codon in the 5’ UTR of *Nnk* [44](Figure 3C). This *piggyBac* insertion, which is marked by a DsRed reporter driven by an eye-specific promoter, is flanked by stop codons in all three reading frames, which prevents translation downstream of the insertion. We refer to this insertion as a *piggyBac-*null allele of *Nnk*. We found that this insertion allele is homozygous lethal but its viability can be fully rescued upon mobilization of the *piggyBac* element, which repairs the intact 5’ UTR in a ‘scarless’ fashion to restore *Nnk* function (schematized in Figure 3C). Furthermore, *piggyBac-*null flies can also be rescued by the *Nnk*-rescue transgene (Figure 3D) (Supplementary Figure S2). Second, we re-examined the previously-generated *CRISPR-null Nnk* allele [43] (Figure 3A). Contrary to previous results, we found that this allele is lethal when paired with a *Nnk*-spanning deficiency (BL9207, Figure 3D) and also fails to complement the *piggyBac*-null allele (Figure 3D) (Supplementary Figure S2). Moreover, the *Nnk-rescue* transgene can restore the viability of the *CRISPR-*null *Nnk* allele (Figure 3D) (Supplementary Figure S2). Based on all these results, we conclude that *Nnk* is unambiguously an essential gene in *D. melanogaster*.

### *Nicknack* is required for larval development in *D. melanogaster*

Next, we investigated at what developmental stage *Nnk-null* progeny die. For this, we crossed *Nnk-null* heterozygotes to each other (Figure 4A). These flies contain the *piggyBac* insertion upstream of the *Nnk* gene (Figure 3C) on one chromosome, along with a balancer chromosome (Tm3G) marked with *GFP* and carrying a wildtype *Nnk* allele. Progeny homozygous for the balancer chromosome Tm3G die as early embryos. As a result, all larvae lacking *GFP* expression are *Nnk-null* homozygotes whereas those that are heterozygous express the *GFP* encoded on the balancer. We conducted egg-lay experiments from crosses between heterozygote *Nnk-null* flies and tracked the developmental progression of *GFP*-negative, *Nnk-null* progeny relative to their *GFP*-expressing heterozygote siblings. We found that *Nnk-null* progeny progress through embryogenesis at the same rate as their heterozygote siblings and are morphologically indistinguishable from heterozygotes until the L1 larval stage. The *Nnk-null* larvae are able to move towards and consume yeast paste much like their heterozygote siblings. However, when heterozygote siblings molt into the L2 stage 48 hours after egg laying (AEL), *Nnk-null* larvae do not molt (Figure 4B). Instead, 48 hours AEL, *Nnk-null* larvae progressively become unable to move or eat, eventually dying by 60 hours AEL.

**Figure 4:**
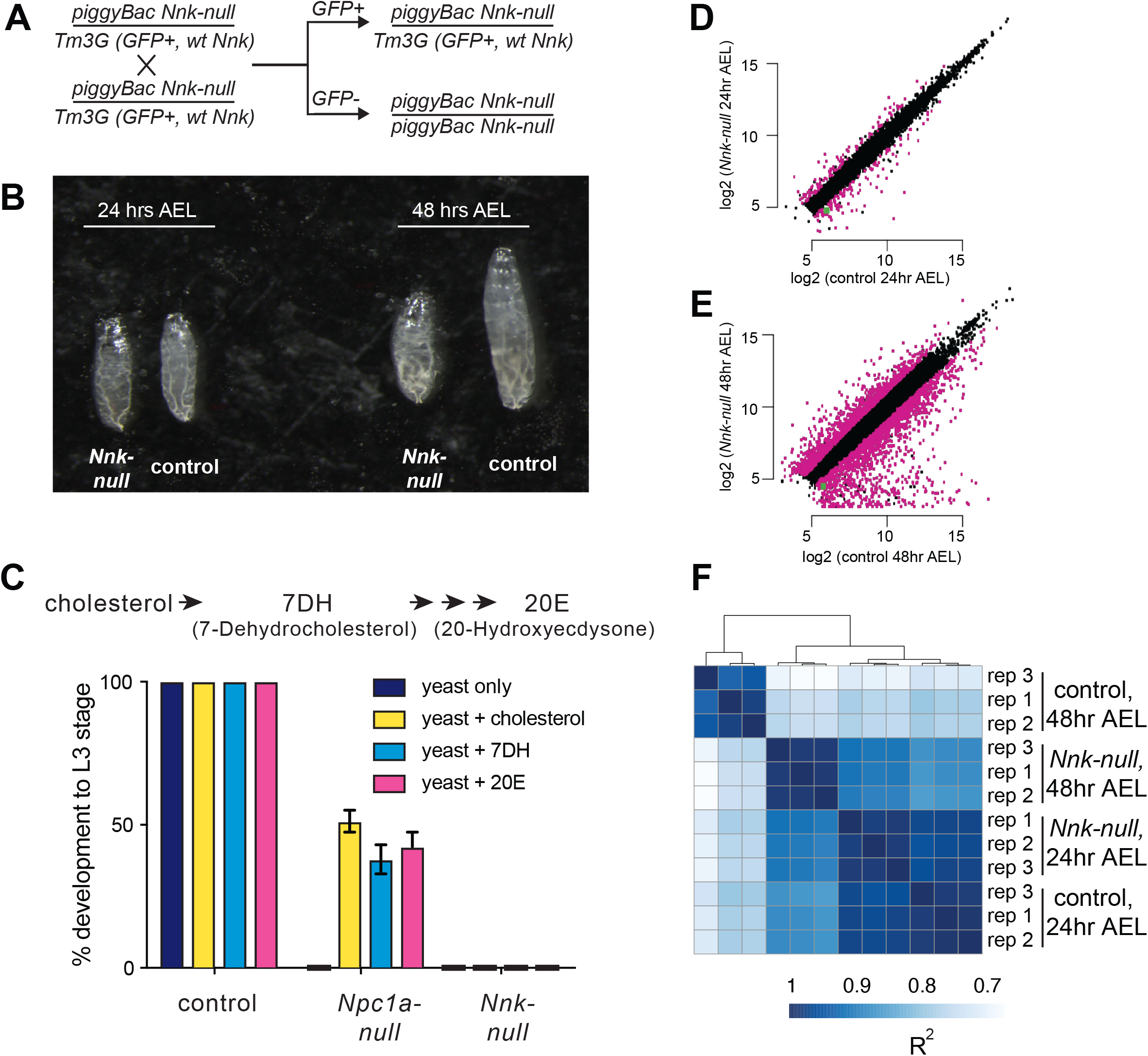
*Nicknack* null larvae arrest in early larval development. (**A**) Schematic of cross to generate homozygous *piggyBac*-null larvae and age-matched control (heterozygote) siblings. *Tm3G* homozygous progeny are not shown, as they die as early embryos. (**B**) 24 hours after egg laying (AEL), *Nnk-null* larvae are morphologically indistinguishable from control larvae. 48 hours AEL, *Nnk-null* larvae are significantly smaller than the age-matched controls and have failed to undergo the first larval molt. (**C**) Adding sterols (cholesterol, 7DH or 20E) to the food of control (w1118) larvae did not alter their ability to develop to L3 larvae within 60 hours AEL. *Npc1a*-null larvae (*Npc1a57/ Npc1a57)* used as a control do not develop on food supplemented with yeast alone but can be partially rescued (>40% molting into L3 larvae) with the addition of cholesterol, 7DH, or 20E. In contrast, *Nnk-null* mutants fail to progress through development even with the addition of dietary sterols. Vertical axis shows the percentage of L1 larvae that progressed to L3 larvae. Graphs show mean of three replicates with error bars showing standard deviation; n>100 for all genotypes and treatments. (**D, E**) Scatter plots showing RNA-seq results for all expressed genes at 24 hrs (D) and 48 hrs (E) AEL. X-axis indicates the expression (normalized abundance) of genes in control larvae whereas the Y-axis shows the expression of genes in *Nnk-null* larvae at the same time point. Magenta dots represent genes significantly over- and under-expressed in *Nnk-null* larvae (green dot represents expression of *Nnk* itself). (**F**) Overall correlations between transcriptome profiles of three replicates each for control and *Nnk-null* larvae reveals that *Nnk-null* transcriptomes 48 hours AEL are more similar to transcriptomes of control and *Nnk-null* larvae 24 hours AEL than they are to age-matched controls, reflecting their developmental delay. Sample-to-sample Spearman R^2^ distance matrix with hierarchical clustering using raw read counts of all 11,428 expressed genes.

The *Nnk-null* larval arrest phenotype is reminiscent of previous findings with the *ZAD-ZNF* genes *Séance, Ouija Board* and *Molting Defective*, which encode proteins necessary for expression of the *Spookier* and *Neverland* genes required for ecdysone biosynthesis [25, 45, 46]. Defects in any of the *Séance, Ouija Board* and *Molting Defective ZAD-ZNF* genes leads to arrest of larval development and death [25, 45, 46]. However, this lethality can be rescued by supplementing the diet with ecdysone or by overexpression of ecdysone biosynthetic enzymes, bypassing the requirement for these three *ZAD-ZNF* genes. We, therefore, tested whether dietary supplementation with an ecdysone precursor, cholesterol (7DH, or 7-Dehydrocholesterol) or ecdysone (20E, or 20-Hydroxyecdysone) could bypass or significantly delay the death of *Nnk-null* larvae (Figure 4C). We found that it could not. In contrast, the same dietary supplementation is able to significantly restore the viability of *Npc1a* larvae, which lack an essential transporter of cholesterol (Figure 4C). Based on these results, we conclude that *Nnk* plays an essential role in larval progression in *D. melanogaster* that is distinct from known steps of the ecdysone biosynthesis pathway.

### Transcriptional consequences of *Nicknack* knockout in *D. melanogaster* larvae

To support our developmental findings, we also performed RNA-seq analyses, comparing the transcriptomes of *Nnk-null* larvae to their age-matched heterozygote siblings. We compared the two genotypes at each of two important time-points. First, we compared transcriptomes 24 hours AEL, when the two genotypes are morphologically indistinguishable in L1 larvae. At this timepoint, we find that 249 genes (2.4% of all expressed genes) are differentially expressed between *Nnk-null* larvae and control larvae, with 116 genes at least two-fold upregulated and 133 genes two-fold downregulated in *Nnk-null* larvae (Figure 4D). Among genes downregulated in *Nnk-null* L1 larvae, we find a significant over-representation of functional categories related to proteolysis and sterol transport (both important functions during larval development), as well as dopamine monooxygenase activity (Supplementary Table S1). In contrast, we find that genes related to lysosome, cytochrome P450s (also important for larval molts), and eye-related functions are upregulated upon *Nnk* loss.

Second, we performed comparisons 48 AEL when the *Nnk-null* larvae are significantly smaller and appear developmentally arrested compared to their age-matched controls. At this timepoint, we find that 3,027 (28.1% of all expressed genes) genes are differentially expressed, with 1,301 genes at least two-fold upregulated and 1,726 genes two-fold downregulated in the *Nnk-null* mutants compared to the age matched controls (Figure 4E). Thus, there are significantly more genes affected by *Nnk* loss by 48 hrs AEL. Intriguingly, clustering samples by the transcriptional profile of all genes shows that *Nnk-null* larvae at 48 hours AEL are transcriptionally more similar to control larvae at 24 hours AEL of either genotype than they are to age-matched control larvae (Figure 4F). The transcriptional status of *Nnk-null* larvae therefore mirrors the phenotypes we observe, displaying a severe developmentally arrested phenotype and transcriptional profile at 48 hours AEL (Figure 4B).

### *Nicknack* encodes a heterochromatin-localizing protein in *D. melanogaster*

Most of the *Odj-Nnk* cluster of *ZAD-ZNF* genes are functionally uncharacterized. Since Odj encodes a heterochromatin-localizing protein [26], we speculated that its close paralog, *Nnk,* might also encode a protein with heterochromatic localization in *D. melanogaster* cells. To test this possibility, we used transient transfections to introduce epitope-tagged *Odj* and *Nnk* genes into *D. melanogaster* Schneider 2 (S2) cells and induced their expression with a heat-shock promoter (Figure 5A). Upon induction, we find that Oddjob has a broad localization pattern within heterochromatin (marked by histone H3 lysine 9 methylation, H3K9me3). We found that the *Nnk-*encoded protein also localizes to heterochromatin, but its localization is restricted to discrete foci, unlike Oddjob (Figure 5A). Since heat-shock induction can alter chromatin properties in cells, we also employed a complementary transient transfection strategy in which we expressed mCherry epitope-tagged *Odj* and *Nnk* genes under the control of a constitutive pCopia promoter in S2 cells. These analyses also revealed a broad heterochromatic localization of Odj and discrete foci within heterochromatin for Nnk protein (Figure 5B). These foci do not overlap with centromeres (identified by the centromeric histone Cid) or dual-strand piRNA clusters (marked by the piRNA-binding HP1 protein Rhino) (Supplementary Figure S3).

**Figure 5:**
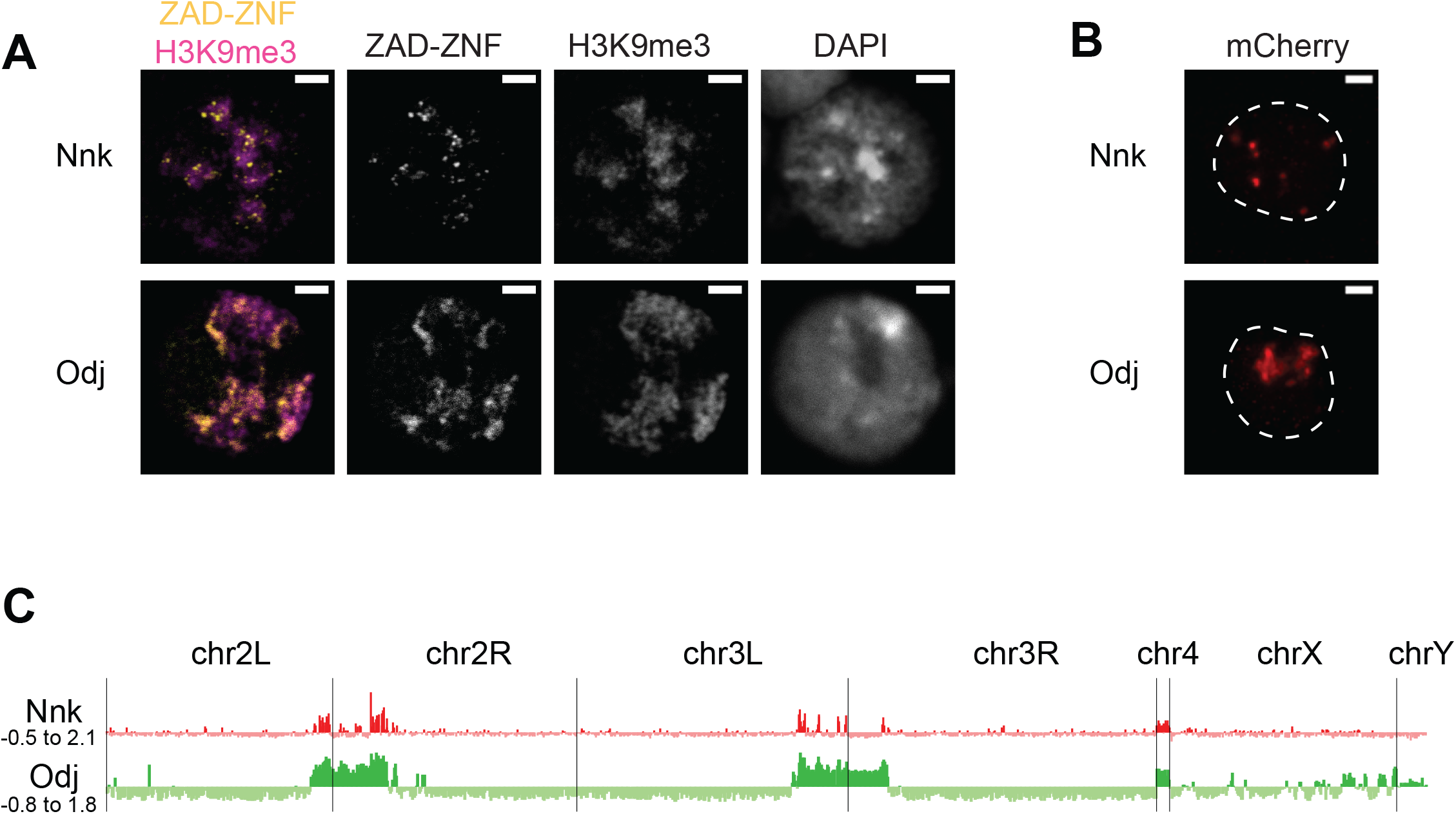
Heterochromatic localization of Odj and Nnk proteins. **(A)** Odj and Nnk proteins localize within heterochromatin (marked by H3K9me3, magenta) of *D. melanogaster* S2 cells. Whereas Oddjob localizes broadly to heterochromatin, Nicknack localizes to discrete foci within heterochromatin. All images are of a representative S2 cell nucleus transfected with a Venus-tagged ZAD-ZNF (yellow) under the control of an inducible heatshock-promoter. DAPI marks DNA in each nucleus. Scale bar = 2 μm. **(B)** Constitutive pCopia-driven expression confirms the broad Odj and discrete Nnk localization within heterochromatin of *D. melanogaster* S2 cells. White dashed line outlines nucleus. Scale bar = 1 μm. **(C)** ChIP-seq analyses confirms that Odj binding is highly enriched throughout heterochromatin, whereas Nnk localizes only to some regions within heterochromatin, and to the 4^th^ chromosome; S2 cells do not have an intact Y chromosome [105]. The y-axis represents normalized ChIP-versus-input log2 ratios, using IGV to visualize and smooth data. Numbers on the y-axis give the range of ratios displayed for each factor

Odj has a broad heterochromatic localization in *D. melanogaster* cells [26], which could result from direct interaction with the heterochromatin protein HP1a. Indeed, Odj has two potential PxVxL motifs, which are putative interaction sites for HP1a [47], in the linker and ZNF domains (Supplementary Figure S4A). Investigating Odj orthologs in other *Drosophila* species revealed that the PxVxL motif in the linker domain is well-conserved, but the one in the ZNF domain is not. We found that mutation of the putative HP1a-interaction site in the linker domain (V164A) converted Odj localization from a broad pattern to a discrete pattern that at least partially overlapped with Nnk (Supplementary Figure S4B). In contrast, mutation of the second putative PxVxL site (V321A) did not significantly affect Odj localization. Our results suggest that the putative PxVxL motif in the Odj linker region is a major contributor to Oddjob’s broad localization to heterochromatin potentially by mediating a direct interaction with HP1a. In the absence of this interaction, Oddjob and Nnk (which lacks a canonical PxVxL motif) localize similarly to discrete foci within heterochromatin (Figure 5B). Our findings suggest that altered protein-protein interactions or DNA-binding specificity via the linker domain may provide a means for functional diversification between closely related ZAD-ZNF paralogs.

To gain deeper insight into the heterochromatic localization of Nnk and Odj, we performed chromatin immunoprecipitation and sequencing (ChIP-seq) using transient transfection of S2 cells with the epitope-tagged pCopia constructs for *Odj* and *Nnk*. Consistent with its cytological localization (Figure 5A and B), we found that Odj was highly enriched throughout the large chromosome regions previously defined as heterochromatin [48] whereas Nnk-bound regions are more narrowly distributed within heterochromatin (Figure 5C). Finer-scale analyses suggested that Nnk and Odj signals are enriched close to transcription start sites classified by modENCODE as residing in TSS-proximal chromatin in S2 cells, both in heterochromatin and euchromatin (Supplementary Figure S5)[49]. However, many of these regions overlap with false positive ‘phantom peaks’ previously identified in ChIP-seq experiments in S2 cells [50] (Supplementary Figure S5). While some heterochromatic TSSs could still represent true binding, we are more confident that the broader occupancy we observe outside TSSs represents Odj and Nnk’s true localization. Moreover, we did not observe a significant effect of *Nnk* loss on heterochromatin-embedded gene expression at the larval L1 stage via RNA-seq analyses (Figure 4D, Supplementary Figure S6). Instead, we found that some heterochromatin-embedded repetitive elements are derepressed upon *Nnk* loss (Supplementary Figure S6). Based on these results, we hypothesize that rather than acting as a transcription regulator for specific heterochromatin-embedded genes like its paralogs *Séance, Ouija Board* and *Molting Defective* [25], *Nnk* may instead repress heterochromatic repeats.

### *D. simulans Nnk* can rescue female, but not male viability of *Nnk-null D. melanogaster*

Our analyses revealed *Nnk* is an evolutionarily young, differentially-retained gene that is essential for larval development in *D. melanogaster*. *Nnk* has also evolved under dramatic positive selection in just the 2.5-million-year divergence between *D. melanogaster* and *D. simulans*, having undergone 52 fixed non-synonymous differences in the 439 aa protein-coding region (Table 2). Given the intriguing correlation between essentiality and positive selection that we had observed in the *ZAD-ZNF* genes, we next investigated whether positive selection of *Nnk* has affected its function.

We first assayed the subcellular localization of epitope-tagged *Nnk* orthologs from *D. melanogaster* and *D. simulans* in *D. melanogaster* S2 cells. We used transient transfections to introduce these genes into S2 cells and used heat shock to drive their expression. We found that both proteins similarly localize to foci within heterochromatin (Figure 6A,B). Thus, the rapid evolution of *Nnk* during *D. melanogaster-D. simulans* divergence has not dramatically affected its gross subcellular localization.

**Figure 6:**
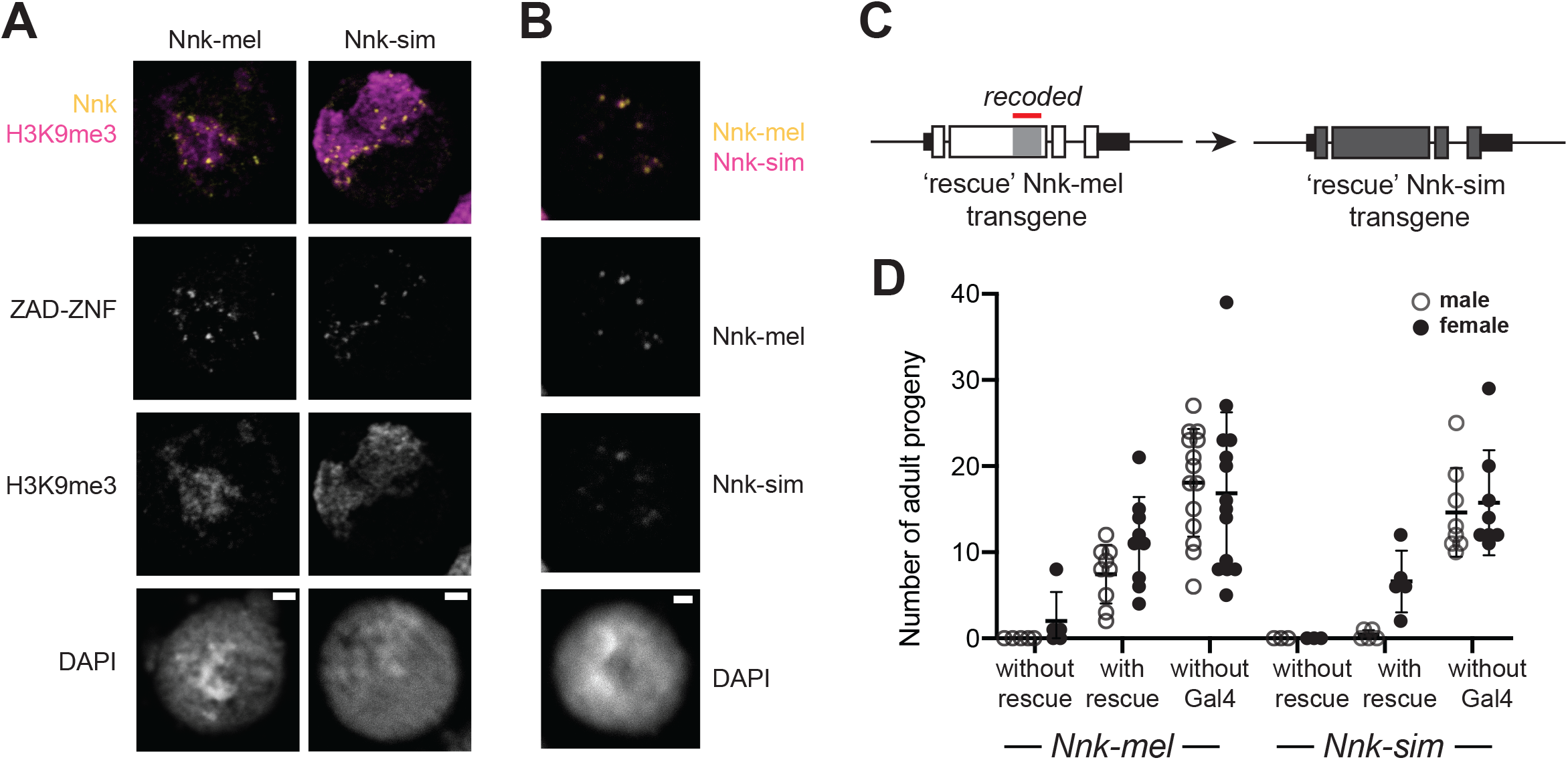
*D. simulans* Nicknack localizes to *D. melanogaster* heterochromatin but can only partially rescue essential function. (**A)** Venus-tagged *D. melanogaster* Nnk (Nnk-mel) or Venus-tagged *D. simulans* Nnk (Nnk-sim) (yellow) localize to foci within heterochromatin (marked by H3K9me3, magenta) in *D. melanogaster* S2 cells. Each image is of a representative S2 cell nucleus with DAPI marking DNA. Scale bar = 2 μm. **(B)** Transient co-transfection of Venus-tagged Nnk-sim (magenta) and FLAG-tagged Nnk-mel (yellow) shows that they overlap in their localization to discrete heterochromatic foci in *D. melanogaster* S2 cells. Scale bar = 2 μm. (**C)** Schematic of rescue *Nnk-sim* transgene, which contains the genomic region 1 kb upstream of the *Nnk* start codon and 1 kb downstream of the stop codon, but with the *D. simulans* coding region (dark gray) from start to stop codon, including introns. The region of *Nnk-mel* targeted by the RNAi hairpin is highlighted by the red line; because of its divergence and codon-optimization, the *Nnk-sim* rescue transgene is resistant to RNAi. **(D)** Ability of *Nnk-mel* and *Nnk-sim* transgenes to rescue ubiquitous knockdown of *Nnk* using Act5C-GAL4 drivers. Progeny that do not inherit the Gal4-driver have no *Nnk* knockdown and are viable. Progeny that inherit the Gal4-driver but not a *Nnk-rescue* transgene are mostly inviable. Closed circles represent number of male adult progeny and open circles represent number of female adult progeny recovered from single pair matings. Horizontal bars show the mean and standard deviation.

Next, we examined the consequences of *Nnk* positive selection on viability in *D. melanogaster*. We created a *D. simulans* Nnk ‘rescue’ transgene (Figure 6C). This rescue transgene is similar to the *D. melanogaster Nnk* rescue construct (Figure 3B), except that it contains the *D. simulans Nnk* coding sequence (codon optimized to *D. melanogaster*) with 1 kb *Nnk*-flanking sequences from *D. melanogaster* (Supplementary Data S3). We introduced this *D. simulans Nnk-*rescue transgene into the same *attP* site on *D. melanogaster* X chromosome as the *D. melanogaster Nnk*-rescue transgene via PhiC31-mediated transgenesis (Figure 6D). The *attP-*insertion rescue design allowed us to put the transgene in the same genetic location in the *Nnk-mel* and *Nnk-sim* rescue crosses, normalizing for variability in expression of transgenes. This allowed a near-isogenic comparison of the *D. simulans* and *D. melanogaster Nnk* transgenes’ ability to rescue inviability of *Nnk-null D. melanogaster* flies, despite their high level of sequence divergence. Unfortunately, low levels of endogenous *Nnk* expression did not allow us to assess whether the expression levels of both *Nnk* transgenes were equivalent.

We crossed heterozygous females either containing one or no copy of the (RNAi-resistant) *Nnk-rescue* transgene and the *Nnk-RNAi* allele to *Act5C-GAL4/CyO-GFP* males. We expect that absence of a rescue transgene would result in lethality of all progeny. In contrast, in the resulting progeny from rescue transgene-bearing females, we expect half of the progeny to inherit the *Nnk-rescue* transgene (see Methods). We found that the *Nnk-mel* transgene significantly rescued *Nnk* knockdown compared to no-transgene controls; however, males were recovered at slightly lower levels than females (67 males: 101 females compared to expectation of 1:1 ratio; p=0.08, Fisher’s exact test). We similarly found that the *D. simulans Nnk* transgene can significantly rescue the lethality caused by knockdown of endogenous *D. melanogaster Nnk* in females (Figure 6C), although at a slightly lower level than *Nnk-mel* rescue. In contrast, rescue of male viability by *D. simulans Nnk* is extremely poor (2 males: 33 females compared to expectation of 1:1 ratio; p<0.0001, Fisher’s exact test) resulting in a severe sex-bias. Thus, *Nnk-sim* is much worse than *Nnk-mel* in rescuing male viability (67:101 versus 2:33, p<0.0001, Fisher’s exact test). Based on these findings, we infer that the hypomorphic rescue of the *D. simulans Nnk-rescue* transgene is further exacerbated in the presence of the heterochromatin-rich Y chromosome, resulting in poor male rescue. Our findings suggest that not only is *Nnk* a positively-selected, essential *ZAD-ZNF* gene, but also that its positive selection is require for optimal function in the *D. melanogaster* genome.

## Discussion

In this study, we explored the relationship between genetic innovation and essentiality in the *ZAD-ZNF* gene family, which encodes the most abundant class of transcription factors in insects. Due to their lineage-specific amplification, protein structure and expression patterns, *ZAD-ZNF* genes were previously hypothesized to be analogous to the KZNF (KRAB-Zinc Finger) transcription factor-encoding gene family found in vertebrates [10, 13], many of which target transposable element sequences (TEs) inserted in the genome. Just like *KZNF* genes in mammals, we find strong evidence for evolutionary dynamism in *Drosophila ZAD-ZNF* genes. For example, we find that only 62 of 91 *ZAD-ZNFs* found in *D. melanogaster* are universally retained in most *Drosophila* species. Furthermore, 13 *ZAD-ZNFs* have evolved under positive selection during *D. melanogaster*-*D. simulans* divergence.

Despite these similarities, however, there are considerable differences between these gene families. First, a direct connection to TEs has only been revealed for one *ZAD-ZNF* gene in *Drosophila*; *CG17801* has a role in regulating *HetA* and *Blood* transposable elements in the female germline [27]. Second, we find that *ZAD-ZNF* genes that evolve under positive selection are more likely to encode essential functions in embryonic axial patterning, larval development, and meiosis [22–25, 30, 31] (Table 2). In contrast, most *KZNF* genes that have been shown to be essential for sterility or viability in mammals are slowly evolving [51, 52]. Finally, unlike *KZNFs* [18], positive selection in *ZAD-ZNFs* is not primarily focused on their C2H2 domains but rather on the poorly-characterized linker domains that connect the ZAD and C2H2 domains (Table 2). We speculate that the linker, which is often comprised of intrinsically disordered domains, may play an important role in chromatin localization of ZAD-ZNF proteins either via direct DNA-binding or via protein-protein interactions [53, 54]. For example, the Oddjob linker domain encodes a PxVxL motif that is important for its broad heterochromatic localization, potentially by mediating direct interaction with HP1a (Figure 5, Supplementary Figure S4).

Based on these dissimilarities, we do not favor the possibility that ZAD-ZNF innovation is driven by arms-races with TEs. Instead, we instead favor the hypothesis that the recurrent adaptation of a subset of *ZAD-ZNF* stems from their roles in heterochromatin organization or regulation of gene expression. Indeed, many genes that encode crucial heterochromatin functions are often critical for viability or fertility, yet are quite variable even among closely-related species [6, 55–58]. Heterochromatin is a gene-poor component of most eukaryotic genomes. Yet, its establishment and maintenance is nevertheless essential for many cellular processes including chromosome condensation and segregation, repression of TEs, and genome stability [59–73]. Thus, the rapid evolution of genes encoding heterochromatin functions might reflect lineage-specific mechanisms to package heterochromatic DNA, or silence TEs.

Several characterized *ZAD-ZNFs* have been found to play key roles at heterochromatin. For example, the Oddjob protein directly binds to HP1a [26] and broadly localizes to heterochromatin (Figure 5). Similarly, ZIPIC, Pita and Zw5 proteins are found at heterochromatin and euchromatin [29], while Séance and Ouija Board control the expression of heterochromatin-embedded genes necessary for larval development [25]. Based on these observations, we propose that *ZAD-ZNF* diversification (marked by gene turnover and positive selection) is driven by the high turnover of sequences embedded within heterochromatin. Although the bulk of heterochromatin is made up of highly repetitive elements such as satellite DNAs and TEs, heterochromatin also harbors many genes that are deeply embedded within heterochromatin [74–79]. These genes, many of which encode essential functions [80], require a heterochromatic environment to ensure their correct expression and regulation [81]. We posit that the constant turnover of flanking and embedded sequence elements such as TEs and satellite DNAs may require constant adaptation of transcription factors required for the proper expression of genes embedded in heterochromatin. *ZAD-ZNF* adaptation might also be necessary to protect against the inappropriate expression of heterochromatin-embedded elements especially at crucial developmental stages.

By testing the causal link between positive selection and essential function, we also find further support for Nnk’s essential function being related to heterochromatin biology. We find that *D. simulans Nnk* can significantly rescue *Nnk-null* inviability in females, but not in males. We speculate that this inability to rescue male viability could be due to the heterochromatin-rich Y chromosome in males. For example, if the *D. melanogaster* Nnk protein, but not the *D. simulans* Nnk protein, could appropriately repress the *D. melanogaster* Y chromosome, this might explain the sex-bias seen in the *D. simulans Nnk* rescue progeny. The Y chromosome is itself not essential for viability or sex determination in *Drosophila*. However, de-repression of the heterochromatin-rich Y chromosome could nevertheless lead to detrimental consequences on larval development. This effect could be direct, leading to inappropriate expression of Y-chromosome-embedded genetic elements that block larval development. Alternatively, this effect could be indirect; de-repression of the Y chromosome could indirectly impact several other chromatin processes genome-wide *e.g.,* due to inappropriate recruitment of transcription factors, which could exacerbate an already hypomorphic function of the *D. simulans Nnk* allele [82–86]. In either scenario, we hypothesize that rapid co-evolution with heterochromatic sequences might have driven the rapid evolution of Nnk and possibly other heterochromatin-interacting ZAD-ZNF proteins.

Based on our findings, we hypothesize that constant adaptation in *ZAD-ZNF* genes is driven by rapid alterations in heterochromatin across *Drosophila* and other insect species. This co-evolutionary arms-race may provide the explanation for the unexpected correlation we find between gene essentiality and innovation in the largest family of transcription factors in insect genomes.

## Methods

### Phylogenomic analysis of *ZAD-ZNF* genes in *Drosophila*

We used the Flybase database (http://flybase.org) to identify all ZAD-containing proteins (Pfam motif PF07776) in 12 sequenced and annotated *Drosophila* species [32]. Using NCBI’s Conserved Domains search, we identified other domains found in the ZAD-containing proteins [87]. To estimate the evolutionary age of these ZAD-containing genes, we used OrthoDB to identify orthologs of the 91 ZAD-containing peptides across *Drosophila* [88].

### Defining gene essentiality

We used FlyBase gene summaries (FlyBase.org) and published studies when available to define gene essentiality. Our criteria for essentiality were broad: if there was a lethal allele reported, we counted the gene as essential for viability in *D. melanogaster.*

### Analyses of positive selection

For the McDonald-Kreitman test, we extracted the gene of interest from *D. melanogaster* population genetic datasets available through Popfly (www.popfly.com) and removed low frequency (<0.05) variants from the dataset to minimize the effects of false positives and low-frequency variants that may not have been subject to selection [37, 89]. We used a manually trimmed alignment of the *D. melanogaster* filtered dataset and the reference *D. simulans* sequence for the McDonald-Kreitman test (http://mkt.uab.es/mkt/) [33, 90].

### Viability studies

We used an *Act-GAL4/CyO-GFP* driver for ubiquitous knockdown. The RNAi lines used to specifically target *Oddjob* cluster genes are the VDRC KK or GD lines: *Oddjob* (27971), *CG17803* (38869), *CG17801* (29501), *CG17806* (40106) and *Nicknack* (102311). RNAi controls used for the experiment were *Cid* (43856) and *HP1B* (26097) for the ubiquitous knockdown. Ubiquitous knockdown of *Cid* produced no viable progeny. We crossed 5 virgin females carrying *Act-GAL4/CyO-GFP* to 3 males of each RNAi line. We allowed the females to lay eggs for 3 days and flipped the flies into fresh vials three times. Each cross was performed in triplicate. Progeny were counted 10-15 days after each cross was set up. A minimum of 20 *CyO-GFP* males and 20 *CyO-GFP* females (*i.e.,* control genotype) were required for us to quantify the crosses. Plotting and statistical analyses were conducted using Graphpad Prism 8 software.

### Defining orthologs

Since *Oddjob* cluster genes experienced numerous independent segmental duplications, it was not possible to determine orthologs by synteny. Instead, we used TBLASTN [91] to identify candidate orthologs of *Oddjob* cluster genes, using the genes in *D. melanogaster* as queries. We used a reciprocal blast search strategy to identify potential orthologs and further investigated these candidates by making a maximum likelihood phylogenetic tree (LG substitution model in PhyML with 100 bootstrap replicates) of a manually trimmed protein alignment (constructed using Clustal Omega program [92] in the Geneious package, www.geneious.com) (Supplementary Data 1, Supplementary Figure 1). We assigned orthologs based on genes that formed a monophyletic clade with the each of the *D. melanogaster Oddjob* cluster genes. We mapped these *Oddjob* cluster orthologs back to each genome assembly to determine the composition of the *Oddjob* locus across *Drosophila* species. In cases where there were other ZAD-ZNFs present, we blasted them against the *D. melanogaster* genome to examine if there were any orthologs present in *D. melanogaster.* If the top hit was not a member of the Oddjob locus, we did not include it in the tree. In the case where there are partial ZAD-ZNFs (containing just the ZAD domain or the zinc finger domains), we performed a blast search against the *D. melanogaster* genome. If the top hit was a member of the *Oddjob* cluster, we assigned orthology by making a phylogenetic tree with all other *Oddjob* cluster orthologs.

### *Nnk*-rescue transgene design, construction, and crosses

We designed a *D. melanogaster* recoded transgene comprising a 3.3 kb fragment containing the genomic region of *Nicknack* plus 1 kb upstream and downstream of the start and stop codons, based on the *D. melanogaster* reference assembly. We recoded the sequence targeted by the VDRC RNAi KK line (103211) by making synonymous changes at each codon. The resulting sequence was synthesized by GENEWIZ Co. Ltd. (Suzhou, China) and cloned into a plasmid we generated that contains 3xP3-DsRed attP, which produces fluorescent red eyes in the adult to mark the presence of the transgene. To generate the *D. simulans* transgenic allele, we codon-optimized the *D. simulans Nnk* coding sequence for the *D. melanogaster* genome using IDT’s codon optimization tool. The resulting sequence was synthesized by GENEWIZ Co. Ltd. (Suzhou, China) and swapped for the *D. melanogaster Nnk* coding sequence in the plasmid described above, using the NEBuilder kit (New England Biolabs). We submitted transgenic constructs to The BestGene Inc. (Chino Hills, CA) for injection into the X-chromosome *attP18* line (BL 32107) using *PhiC31* site specific integration [93].

For the transgene rescue cross, we crossed 5 virgin female flies bearing one copy of *Nnk-rescue* transgene (on the X chromosome) and the RNAi allele (on the second chromosome) to 3 *Act5C-GAL4/CyO-GFP* males. We allowed the females to lay eggs for 3 days and flipped the cross three times. We set up each cross in triplicate and progeny were counted after 10-15 days. At least 20 progeny were counted per replicate. Each cross had at least three replicates. All flies were raised at 25°C.

### Characterization of *Nnk-null* mutants

We placed 50-75 flies heterozygous for the *Nnk-null* allele (*Nnk pBac null/TM3G)* or for the *Nnk* CRISPR allele (*Nnk CRISPR /TM3G)* into a small embryo collection cage containing a grape-juice plate with a thin strip of yeast paste and collected embryos for 3 hours at 25°C. We transferred the larvae to fresh grape-juice plates containing yeast paste daily and scored developmental stage by mouth hook morphology. We used fluorescence to distinguish between heterozygotes (GFP-positive larvae) and homozygotes (GFP-negative larvae). For the trans-heterozygote evaluation, we crossed 30-40 virgin female *Nnk CRISPR/TM3G* to 10 *Nnk pBac null/TM3G* males. Crosses were done in triplicate and at least 100 progeny were counted per cross.

### Larval collection for RNA sequencing

We placed 50-75 flies heterozygous for the *Nnk-null* allele (*Nnk pBac/TM3G)* into a small embryo collection cage containing a grape-juice plate with a thin strip of yeast paste and collected embryos for 3 hours at 25°C. The first time point was collected 24 hours after egg laying (AEL) and the second 48 AEL. We transferred the larvae to fresh grape-juice plates containing yeast paste daily.

### RNA extractions and library preparation

Whole larvae (~30 animals at 24 hrs AEL and ~20 animals at 48 hrs AEL for each sample; RNA from each time point and genotype was prepared in triplicate) were ground in a 1.5 mL Eppendorf tube containing 50 μL of TRIzol reagent using a DNase, RNase and DNA free 1.5 mL pestle. 450 μL of TRIzol reagent was added after grinding. Immediately, we added 500 μL of chloroform and the tube was inverted gently 2-3 times. We removed the aqueous phase into a fresh tube containing 1 mL of 200 proof EtOH and mixed by inversion. The mixture was then bound to a Zymo-spin column according to the manufacturer’s instructions (Zymo Research). We followed the DNase extraction and purification protocol outlined in the RNA Clean and Concentrator kit (Zymo Research). We eluted the RNA in 15 μL of DNase/RNase-free water and immediately placed the samples at −80°C. We checked the quality of the samples with a 2200 Tapestation (Agilent Technologies) and selected samples that had an RNA integrity number > 9.0 for library preparation. Library construction and Illumina 150-bp paired-end RNA-sequencing were conducted at Novogene Bioinformatics Technology Co., Ltd (Beijing, China).

### Transcriptome data analysis

We used Kallisto [94] to quantify abundances of the *D. melanogaster* reference transcriptome (refMrna.fa for dm6, obtained from UCSC Genome Browser Oct 16^th^, 2018, which contains 34,114 transcripts). For each transcript, we acquired the gene name using R (org.Dm.eg.db). Kallisto counts were read into R using the tximport package [95] and using the summarizeToGene function, we summarized alternative splice-form counts into a single count per gene. We used DESeq2 to identify differentially expressed genes with adjusted p-value <=0.05 and absolute log_2_(fold change) >=1 [96], comparing *Nnk-null* larvae to controls for each timepoint separately. Before performing each comparison, we excluded genes with low expression (<100 counts total across all samples): this filtering yields 10,574 ‘expressed’ genes in 24 hour AEL larvae, and 10,758 ‘expressed’ genes in 48 hour AEL larvae.

We performed Gene Ontology (GO) enrichment analysis on each of four gene lists: 116 over-expressed genes and 133 under-expressed genes in 24 hour AEL larvae, and 1,301 over-expressed genes and 1,726 under-expressed genes in 48 hour AEL larvae. We used the Bioconductor GOstats package to perform conditional hypergeometric enrichment tests for each of three ontologies (biological process, molecular function and cellular component) [97, 98]. For the ‘universe’ of all genes examined (*i.e.,* background) we used the corresponding list of ‘expressed’ genes at each developmental timepoint. We report only over-represented categories with p<0.001, and use annotations found in the org.Dm.eg.db Bioconductor package.

To estimate expression levels aggregated across all instances of each repeat type, we took an approach similar to that of Day *et al*. [99]. Here, rather than mapping reads to the typical reference genome assmbly, we constructed a ‘repeat assembly’, where we used RepeatMasker annotations for dm6 (obtained from UCSC) to extract and concatenate all instances of each repeat type, adding 75bp flanking sequences (half the length of each read) and inserting 150 N bases between each instance. The repeat assembly therefore consists of a single ‘chromosome’ for each repeat type. We then used BWA-mem to map sequences as single reads (not paired-ends) to the repeat assembly [100], and samtools to count reads mapping to each repeat type (‘chromosome’). We combined counts for pairs of simple repeats that represent the reverse-complement of one another. We used DESeq2 to perform statistical analyses on raw counts, using the total number of sequenced fragments as size factors [101]. We normalized counts by dividing each by the number of reads sequenced for that sample in millions.

### Dietary sterol supplementation

To evaluate the ability of dietary sterols to rescue *Nnk-null* phenotype, we carried out sterol supplementation as previously outlined [25]. Briefly, we mixed together 20 mg of dry yeast in 38 uL of water. To this yeast paste, we added either 2 uL of EtOH (negative control) or 2 uL of EtOH plus cholesterol (Sigma), 7-dehydrocholestrol (Sigma) or 20-hydroxyecdysone. Stocks used for these experiments were balanced over GFP-balancers (*Npc1a57/CyO-GFP* and *Nnk-null/TM3G*). The control used for these experiments was the *w1118* stock. Eggs were laid on yeasted grape-juice plates for 3 hours at 25°C. 24 hours AEL, GFP-negative larvae were transferred onto grape plates containing fresh yeast paste at 25°C. For 72 hours AEL, larvae were transferred to fresh grape-juice plates containing yeast paste daily and scored for their developmental stage based on the morphology of their mouth hooks. *Npc1a57/CyO-GFP* stocks were a kind gift from Leo Pallanck [102].

### Tissue culture and transfection

*Oddjob* cluster genes were amplified from genomic DNA from *D. melanogaster* and *D. simulans* and directionally cloned into pENTER/D-TOPO (ThermoFisher) according to the manufacturer’s instructions. We verified that clones had the appropriate insertions by sequencing. We used LR clonase II (ThermoFisher) to get each *Oddjob* cluster gene into the *Drosophila* Gateway Vector destination vector to express the gene of interest with an N-terminal Venus tag under the control of the *D. melanogaster Hsp70* promoter (pHVW). For Rhino localization in S2 cells, the *Rhino* gene was cloned into the *Drosophila* Gateway Vector destination vector to enable expression of Rhino with an N-terminal 3xFlag tag under the control of the *D. melanogaster Hsp70* promoter (pHFW). *Nnk* and *Odj* were also cloned into pCopia vectors containing N-terminal and C-terminal mCherry epitope tags, respectively, and HP1a was previously tagged with mCerulean at the N-terminus.

Schneider 2 cells were obtained from the *Drosophila* Genomics Resource Center (Bloomington, IN, USA) and grown at 25°C in M3+BPYE+10%FCS. For the transfections, one million cells were seeded, and one day later 2 micrograms of plasmid DNA was transfected into cells using Xtremegene HP transfection reagent (Roche) according to the manufacturer’s specifications. For the Rhino localization experiment, cells were co-transfected with 1 ug of GFP-tagged ZAD-ZNF vector and 1 ug of Flag-tagged Rhino vector. Cells were allowed to recover for 24 hours post transfection, heat shocked for 1 hour at 37°C and recovered for 3 hours at 25°C prior to fixation. For the pCopia vectors, transient transfections were conducted on S2 cells using the TransIT-2020 reagent (Mirus), and live imaging was performed 72 hours later, using an Applied Precision Deltavision microscope and analyzed using SoftworX software.

For the heatshock vector transfections, cells were transferred to coverslips for 30-45 minutes prior to starting the immunohistochemistry protocol. 0.5% sodium citrate hypotonic solution was added to the coverslip for 10 minutes to swell cells, which was then spun at 1900 rpm for 1 minute in a Cytospin III to remove the cytoplasm from the cells. The sodium citrate was immediately removed and cells were subsequently fixed. For fixation, 4% PFA + PBST (PBS + 0.1% Triton) was added to the cells for 10 minutes. Coverslips were then washed in PBST and blocked for 30 minutes in PBST + 3% BSA. Cells were incubated with primary antibody overnight at 4°C in a humid chamber. For immunolocalization, the following dilutions of primary antibodies were used: GFP (Abcam AB13970) 1:1000, H3K9me3 (Abcam AB8898) 1:500 M2 FLAG (Sigma-Aldrich F4042) 1:1000. After washing with PBST three times for 10 minutes per wash, the following fluorescent secondary antibodies were used at 1:1000 dilution: goat anti-chicken (Invitrogen Alexa Fluor 488, A-11039), goat anti-rabbit (Invitrogen Alexa Fluor 568, A-11011) and goat anti-mouse (Invitrogen Alexa Fluor 568, A-11031). The cells were incubated with 1x DAPI in the final wash and mounted in SlowFade Gold Mounting Medium (ThermoFisher). We imaged cells on a Leica TCS SP5 II confocal microscope with LASAF software and images were processed using ImageJ and were representative of the cell population.

### ChIP-seq analyses

S2 cells transfected with either mCherry-Nnk or Odj-mCherry for 72 hours were fixed with 1% paraformaldehyde for 10 minutes and sheared with Bioruptor sonicator (Diagenode) to obtain chromatin. Chromatin fragments were confirmed to contain DNA in the 200-500 bp size range using a Bioanalyzer (Agilent). Each immunoprecipitation was performed on chromatin from 2×10^7^ S2 cells by overnight incubation with Protein-G Dynabeads and 5 ug of anti-mCherry antibody (Novus). Library construction from immunoprecipitated DNA was conducted using TruSeq sample preparation kits (Illumina). 150bp paired-end sequences were generated by the Vincent J. Coates Genomics Sequencing Laboratory at UC Berkeley.

We used BWA-mem to map paired reads to version 6 of the *D. melanogaster* genome assembly (dm6) from which we had removed unplaced scaffolds [100]. We used deepTools’ bamCompare (with the ‘--binsize 1 --extendReads’ options) to obtain log2 ratios of fragment coverage for matched ChIP and input samples, and visualized those ratios in IGV [103]. We further visualized ChIP-seq signal around TSSs using deepTools’ computeMatrix and plotHeatmap tools, using TSS annotations obtained from the TxDb.Dmelanogaster.UCSC.dm6.ensGene BioConductor package, taking the most upstream TSS for genes with alternative start sites. We split TSS annotations according to whether they were within ‘TSS-proximal’ (active) chromatin according to modENCODE’s 9 state annotation for S2 cells, obtained from http://intermine.modencode.org and converted from dm3 to dm6 coordinates using UCSC’s liftOver tool [104]. We further split the ‘active’ TSSs according to whether they are within cytogenetic heterochromatin, using coordinates from Table S2 of Hoskins et al [48] as well as the whole of chromosomes 4 and Y.

## Acknowledgements

We thank past and present members of the Malik lab for their comments and discussion. In addition, we are especially grateful to Celeste Berg and Barbara Wakimoto for their helpful comments during this project, and to Leo Pallanck for helpful discussions and generous gift of fly strains. We also thank Eric Lai for sharing the CRISPR-null *Nnk* fly line. We thank Ching-Ho Chang, Michelle Hays, Pravrutha Raman and Aida de la Cruz for their comments on the manuscript. This work was supported by the following grants: NIH training grant T32 CA009657, NIH F30 CA225077 fellowship, and Julie Tall Achievement Rewards for College Scientists (ARCS) endowment from the Seattle Chapter of the ARCS Foundation (to B.K); NIH R01 GM117420 (to G.H.K.), NIH R01 GM074108 and HHMI Investigator grant (to H.S.M.). The funders had no role in study design, data collection and analysis, decision to publish, or preparation of the manuscript. The authors declare that they have no conflicts of interest.

## Supplementary Materials

**Supplementary Data S1.** Multiple alignment of the ZNF domain encoded by all genes found within the *Odj-Nnk ZAD-ZNF* cluster across 20 *Drosophila* species.

**Supplementary Data S2.** Nucleotide sequence of the *Nnk-mel* rescue transgene that has been recoded by synonymous substitutions to make it impervious to RNAi-mediated knockdown.

**Supplementary Data S3.** Nucleotide sequence of the *Nnk-sim* rescue transgene that has been codon-optimized to match *D. melanogaster* codon preferences.

**Supplementary Table S1.** Gene ontology analysis of transcriptional dysregulation in *Nnk-null* L1 larvae in *D. melanogaster* reveals categories of genes that are significantly over- or under-expressed upon loss of *Nnk*.

**Supplementary Figure S1.** Phylogenetic analysis of all genes found within the *Odj-Nnk ZAD-ZNF* cluster across 20 *Drosophila* species based on a multiple alignment of their ZNF domains (Supplementary data S1). Bootstrap numbers in support of each orthology group (as a percentage of 100 bootstrap trials) are reported. Numbers next to the species names refer to lineage-specific gene duplications within orthology groups (Figure 2C).

**Supplementary Figure S2.** Various Nnk disruption alleles can be all rescued significantly with a *Nnk-mel* rescue transgene. Numbers in parentheses refer to expected numbers in case of full rescue. The *Nnk-rescue* allele can rescue otherwise inviable *piggyBac-null/piggyBac-null, CRISPR-null/CRISPR-null*, and *Nnk^RNAi^/Act5C-Gal4* progeny.

**Supplementary Figure S3**. Nnk localization in heterochromatin in *D. melanogaster* S2 cells. Venus-tagged Nnk or Odj proteins were induced via heat shock, and nuclei were imaged with either **(A)** Cid (to mark centromeres) or **(B)** Rhino (to mark dual-strand piRNA clusters). As shown in Figure 5, Odj has a broad heterochromatin localization whereas Nnk localizes to discrete foci, which do not overlap with centromeres or dual-strand piRNA clusters. Inset refers to specific foci to highlight the absence of significant overlap. Scale bars= 2μm.

**Supplementary Figure S4.** Mutation of the Odj PxVxL motif in the linker disrupts its broad localization within heterochromatin. **(A)** Alignment of the two PxVxL motifs in Oddjob across *Drosophila* species show that the first PxVxL motif in linker region is highly conserved while the second PxVxL motif, within the ZNF domain, is not. **(B)** Mutation of the PxVxL motif within the linker region changes Odj’s (magenta) broad localization to discrete foci within heterochromatin like Nnk (yellow). Mutation of the PxVxL motif within the ZNF has no significant change in localization of Odj within heterochromatin (marked by HP1a, cyan). Scale bars= 1μm.

**Supplementary Figure S5.** Additional analysis of ChIP-seq signals. **(A)** Higher resolution view of a selected 168kb region in the heterochromatic region of chromosome 2L, using IGV for visualization. We show log2 (ChIP / input) ratios for the first replicate each for Nicknack and Oddjob, as well as for two negative control samples (Rsf1mut and Acf1mut) that highlight ‘phantom peak’ regions where false positive signals often occur [50]. **(B)** ChIP-seq signals (log2 ChIP / input) in regions of 10kb surrounding all TSSs in the genome, visualized using deepTools. We split TSSs into groups according to whether they are in active chromatin in S2 cells, according to modENCODE’s “TSS-proximal” state, and further split the active TSSs according to whether they are in cytogenetic heterochromatin or euchromatin. In the heatmap region of the plot (variably shaded boxes), very narrow rows represent 10kb regions surrounding each TSS, and the color scale shows the ChIP/input ratio, with columns for each ChIP/input comparison. Graphs above heatmaps show mean signal profiles for each region type, with inactive TSSs in yellow, euchromatic active TSSs in light blue, and heterochromatic TSSs in dark blue. For both Nicknack and Oddjob, the first replicate showed the strongest signal and the third replicate showed the weakest signal, likely due to use of different batches of antibody. While inactive TSSs show very little signal in any sample, almost all active TSSs have ChIP signal in the ‘phantom’ samples as well as the Oddjob and Nicknack samples, indicating that this signal could represent a ChIP-seq artifact [50]. However, for both Nicknack and Oddjob (but not the ‘phantom’ samples), ChIP signal is stronger and more broadly distributed at active TSSs within heterochromatin than at active TSSs within euchromatin, suggesting that some true signal remains.

**Supplementary Figure S6. (A)** Scatter plots showing RNA-seq results for all expressed genes at 24 hrs AEL. X-axis indicates the expression (normalized abundance) of genes in control larvae whereas the Y-axis shows the expression of genes in *Nnk-null* larvae at the same time point. Grey dots represent euchromatic genes, whereas black dots represent heterochromatin-embedded genes, with magenta dots represent heterochromatic genes that are significantly over- and under-expressed in *Nnk-null* larvae. **(B)** Some repeat elements are misregulated in *Nnk-null* larvae. We used an approach similar to that of Day *et al.* [99] to obtain aggregate counts of reads mapping to all instances of each repeat type. Most repeat types are expressed at similar levels in control larvae 24 hours AEL and *Nnk-null* larvae 24 hours AEL, but a few (shown in magenta) are misregulated. Six repeat types are de-repressed in *Nnk-null* larvae, and two types are expressed at lower levels (absolute log2-fold-change >= 1, adjusted p-value <= 0.05). **(C)** Detailed view of misregulated repeats. Bars show triplicate means of normalized expression levels in reads per million sequenced (RPM), and dots show individual triplicate values.

