## Supplementary Table S1 for "Innovation of heterochromatin functions drives rapid evolution of essential ZAD-ZNF genes in *Drosophila*"

| Gene list | Number of genes tested | Analysis type | Functional category description | Category ID | P-value | Fold enrichment | Number of affected genes | Number of genes in category | Number of genes expected by chance | Percent of category | Percent of genes tested | Gene names | Gene descriptions |
|---|---|---|---|---|---|---|---|---|---|---|---|---|---|
| Over-expressed 24 AEL | 116 | Biological process | desensitization of G protein-coupled receptor signaling pathway by arrestin | GO:0002032 | 8.00E-05 | Inf | 2 | 2 | 0.02 | 100.0% | 1.7% | Arr1,Arr2 | Arrestin 1,Arrestin 2 |
| Over-expressed 24 AEL | 116 | Biological process | negative adaptation of signaling pathway | GO:0022401 | 8.00E-05 | Inf | 2 | 2 | 0.02 | 100.0% | 1.7% | Arr1,Arr2 | Arrestin 1,Arrestin 2 |
| Over-expressed 24 AEL | 116 | Biological process | negative regulation of G protein-coupled receptor signaling pathway | GO:0045744 | 8.00E-05 | Inf | 2 | 2 | 0.02 | 100.0% | 1.7% | Arr1,Arr2 | Arrestin 1,Arrestin 2 |
| Over-expressed 24 AEL | 116 | Biological process | cellular response to light stimulus | GO:0071482 | 2.00E-04 | 16.01 | 4 | 33 | 0.3 | 12.1% | 3.4% | Arr1,Arr2,Rh5,Rh6 | Arrestin 1,Arrestin 2,Rhodopsin 5,Rhodopsin 6 |
| Over-expressed 24 AEL | 116 | Biological process | cellular response to abiotic stimulus | GO:0071214 | 0.00021 | 10.48 | 5 | 61 | 0.55 | 8.2% | 4.3% | Arr1,Arr2,Ku80,Rh5,Rh6 | Arrestin 1,Arrestin 2,Ku80,Rhodopsin 5,Rhodopsin 6 |
| Over-expressed 24 AEL | 116 | Biological process | homeostatic process | GO:0042592 | 0.00034 | 3.66 | 12 | 421 | 3.78 | 2.9% | 10.3% | Arr1,Arr2,ATPCL,CG12896,CG6908,Gba1a,Ku80,mag,mre11,NLaz,ver,Vha100-4 | Arrestin 1,Arrestin 2,ATP citrate lyase,Glucocerebrosidase 1a,Ku80,magro,meiotic recombination 11,Neural Lazarillo,uncharacterized protein,Vacuolar H[+] ATPase 100kD subunit 4,verrocchio |
| Over-expressed 24 AEL | 116 | Biological process | visual perception | GO:0007601 | 0.00043 | 24.58 | 3 | 17 | 0.15 | 17.6% | 2.6% | Crys,Rh5,Rh6 | Crystallin,Rhodopsin 5,Rhodopsin 6 |
| Over-expressed 24 AEL | 116 | Biological process | phototransduction | GO:0007602 | 0.00074 | 11.04 | 4 | 46 | 0.41 | 8.7% | 3.4% | Arr1,Arr2,Rh5,Rh6 | Arrestin 1,Arrestin 2,Rhodopsin 5,Rhodopsin 6 |
| Over-expressed 24 AEL | 116 | Biological process | metarhodopsin inactivation | GO:0016060 | 0.00078 | 75.53 | 2 | 5 | 0.04 | 40.0% | 1.7% | Arr1,Arr2 | Arrestin 1,Arrestin 2 |
| Over-expressed 24 AEL | 116 | Molecular function | opsin binding | GO:0002046 | 8.90E-05 | Inf | 2 | 2 | 0.02 | 100.0% | 1.7% | Arr1,Arr2 | Arrestin 1,Arrestin 2 |
| Over-expressed 24 AEL | 116 | Molecular function | G protein-coupled photoreceptor activity | GO:0008020 | 0.00027 | 214.21 | 2 | 3 | 0.03 | 66.7% | 1.7% | Rh5,Rh6 | Rhodopsin 5,Rhodopsin 6 |
| Over-expressed 24 AEL | 116 | Cellular component | rhabdomere | GO:0016028 | 6.60E-05 | 21.93 | 4 | 30 | 0.22 | 13.3% | 3.4% | Arr1,Arr2,Rh5,Rh6 | Arrestin 1,Arrestin 2,Rhodopsin 5,Rhodopsin 6 |
| Under-expressed 24 AEL | 133 | Biological process | hemolymph coagulation | GO:0042381 | 2.30E-06 | 68.92 | 4 | 9 | 0.11 | 44.4% | 3.0% | Hml,Npc2g,Npc2h,PPO1 | Hemolectin,Niemann-Pick type C-2g,Niemann-Pick type C-2h,Prophenoloxidase 1 |
| Under-expressed 24 AEL | 133 | Biological process | regulation of body fluid levels | GO:0050878 | 8.00E-05 | 21.51 | 4 | 20 | 0.24 | 20.0% | 3.0% | Hml,Npc2g,Npc2h,PPO1 | Hemolectin,Niemann-Pick type C-2g,Niemann-Pick type C-2h,Prophenoloxidase 1 |
| Under-expressed 24 AEL | 133 | Biological process | intracellular cholesterol transport | GO:0032367 | 0.00013 | 42.63 | 3 | 9 | 0.11 | 33.3% | 2.3% | Npc2d,Npc2g,Npc2h | Niemann-Pick type C-2d,Niemann-Pick type C-2g,Niemann-Pick type C-2h |
| Under-expressed 24 AEL | 133 | Biological process | intracellular lipid transport | GO:0032365 | 0.00025 | 31.97 | 3 | 11 | 0.13 | 27.3% | 2.3% | Npc2d,Npc2g,Npc2h | Niemann-Pick type C-2d,Niemann-Pick type C-2g,Niemann-Pick type C-2h |
| Under-expressed 24 AEL | 133 | Biological process | peptidyl-proline hydroxylation to 4-hydroxy-L-proline | GO:0018401 | 0.00034 | 28.41 | 3 | 12 | 0.14 | 25.0% | 2.3% | CG31371,CG9698,PH4alphaMP | prolyl-4-hydroxylase-alpha MP,uncharacterized protein |
| Under-expressed 24 AEL | 133 | Biological process | proteolysis | GO:0006508 | 4.00E-04 | 2.68 | 19 | 684 | 8.16 | 2.8% | 14.3% | CG11842,CG12951,CG16710,CG18477,CG31780,CG34437,CG4017,CG4408,CG4914,Jon65Ai,Jon65Aii,Jon66Ci,Jon66Cii,Jon74E,klhl10,RnrS,SPH93,thetaTry,Uch | Jonah 65Ai,Jonah 65Aii,Jonah 66Ci,Jonah 66Cii,Jonah 74E,kelch like family member 10,Ribonucleoside diphosphate reductase small subunit,Serine protease homolog 93,thetaTrypsin,Ubiquitin carboxy-terminal hydrolase,uncharacterized protein |

| | | | | | | | | | | | | | |
|---|---|---|---|---|---|---|---|---|---|---|---|---|---|
| Under-expressed 24 AEL | 133 | Biological process | protein hydroxylation | GO:0018126 | 0.00083 | 19.66 | 3 | 16 | 0.19 | 18.8% | 2.3% | CG31371,CG9698,PH4alphaMP | prolyl-4-hydroxylase-alpha MP,uncharacterized protein |
| Under-expressed 24 AEL | 133 | Molecular function | serine-type endopeptidase activity | GO:0004252 | 9.20E-09 | 9.63 | 13 | 146 | 1.67 | 8.9% | 9.8% | CG11842,CG12951,CG16710,CG31780,CG34437,CG4914,Jon65Ai,Jon65Aii,Jon66Ci,Jon66Cii,Jon74E,SPH93,thetaTry | Jonah 65Ai,Jonah 65Aii,Jonah 66Ci,Jonah 66Cii,Jonah 74E,Serine protease homolog 93,thetaTrypsin,uncharacterized protein |
| Under-expressed 24 AEL | 133 | Molecular function | serine hydrolase activity | GO:0017171 | 4.70E-08 | 8.3 | 13 | 167 | 1.91 | 7.8% | 9.8% | CG11842,CG12951,CG16710,CG31780,CG34437,CG4914,Jon65Ai,Jon65Aii,Jon66Ci,Jon66Cii,Jon74E,SPH93,thetaTry | Jonah 65Ai,Jonah 65Aii,Jonah 66Ci,Jonah 66Cii,Jonah 74E,Serine protease homolog 93,thetaTrypsin,uncharacterized protein |
| Under-expressed 24 AEL | 133 | Molecular function | peptidase activity, acting on L-amino acid peptides | GO:0070011 | 7.60E-06 | 4.25 | 16 | 390 | 4.47 | 4.1% | 12.0% | CG11842,CG12951,CG16710,CG31780,CG34437,CG4017,CG4408,CG4914,Jon65Ai,Jon65Aii,Jon66Ci,Jon66Cii,Jon74E,SPH93,thetaTry,Uch | Jonah 65Ai,Jonah 65Aii,Jonah 66Ci,Jonah 66Cii,Jonah 74E,Serine protease homolog 93,thetaTrypsin,Ubiquitin carboxy-terminal hydrolase,uncharacterized protein |
| Under-expressed 24 AEL | 133 | Molecular function | oxidoreductase activity, acting on diphenols and related substances as donors, oxygen as acceptor | GO:0016682 | 0.00013 | Inf | 2 | 2 | 0.02 | 100.0% | 1.5% | PPO1,PPO2 | Prophenoloxidase 1,Prophenoloxidase 2 |
| Under-expressed 24 AEL | 133 | Molecular function | L-DOPA monooxygenase activity | GO:0036263 | 0.00013 | Inf | 2 | 2 | 0.02 | 100.0% | 1.5% | PPO1,PPO2 | Prophenoloxidase 1,Prophenoloxidase 2 |
| Under-expressed 24 AEL | 133 | Molecular function | dopamine monooxygenase activity | GO:0036264 | 0.00013 | Inf | 2 | 2 | 0.02 | 100.0% | 1.5% | PPO1,PPO2 | Prophenoloxidase 1,Prophenoloxidase 2 |
| Under-expressed 24 AEL | 133 | Molecular function | oxidoreductase activity, acting on single donors with incorporation of molecular oxygen | GO:0016701 | 0.00014 | 17.97 | 4 | 24 | 0.28 | 16.7% | 3.0% | CG31371,CG6910,CG9698,PH4alphaMP | prolyl-4-hydroxylase-alpha MP,uncharacterized protein |
| Under-expressed 24 AEL | 133 | Molecular function | procollagen-proline 4-dioxygenase activity | GO:0004656 | 0.00023 | 33.38 | 3 | 11 | 0.13 | 27.3% | 2.3% | CG31371,CG9698,PH4alphaMP | prolyl-4-hydroxylase-alpha MP,uncharacterized protein |
| Under-expressed 24 AEL | 133 | Molecular function | peptidyl-proline dioxygenase activity | GO:0031543 | 6.00E-04 | 22.24 | 3 | 15 | 0.17 | 20.0% | 2.3% | CG31371,CG9698,PH4alphaMP | prolyl-4-hydroxylase-alpha MP,uncharacterized protein |
| Under-expressed 24 AEL | 133 | Molecular function | sterol binding | GO:0032934 | 0.00073 | 20.53 | 3 | 16 | 0.18 | 18.8% | 2.3% | Npc2d,Npc2g,Npc2h | Niemann-Pick type C-2d,Niemann-Pick type C-2g,Niemann-Pick type C-2h |

| | | | | | | | | | | | | | |
|---|---|---|---|---|---|---|---|---|---|---|---|---|---|
| Over-expressed 48 AEL | 1301 | Biological process | nervous system process | GO:0050877 | 2.40E-09 | 2.23 | 86 | 410 | 45.67 | 21.0% | 6.6% | 2mit,5-HT1A,Ac78C,Alk,alpha-Est1,amn,Arr1,Arr2,Ca-alpha1T,cac,CG2698,CG31300,CG34371,CG4025,CG7778,CG9447,CG9492,CheB42c,chico,Desi,Dhc93AB,dia,dlp,Dop1R1,Dop2R,dpr11,dpr12,dpr16,dpr18,dpr2,dpr20,dpr4,dpr5,dpr6,drl,dysc,eag,Ets96B,FoxP,futsch,GluClalpha,Gr33a,Gr39a,Gr94a,Grd,hiw,iav,inaC,Ir21a,Ir25a,Ir76b,meru,mGluR, | 2mit,5-hydroxytryptamine (serotonin) receptor 1A,Adenylyl cyclase 78C,alpha-Esterase-1,amnesiac,Anaplastic lymphoma kinase,Arrestin 1,Arrestin 2,Ca[2+]-channel protein alpha[[1]] subunit T,cacophony,Chemosensory protein B 42c,chico,dally-like,defective proboscis extension response 11,defective proboscis extension response 12,defective proboscis extension response 16,defective proboscis extension response 18,defective proboscis extension response 2,defective proboscis extension response |
| Over-expressed 48 AEL | 1301 | Biological process | signaling | GO:0023052 | 9.60E-08 | 1.55 | 237 | 1578 | 175.78 | 15.0% | 18.2% | amn,Arr1,Atg9,bun,Ca-alpha1T,cac,CCKLR-17D1,CCKLR-17D3,CG10362,CG10702,CG13995,CG1657,CG16721,CG32547,CG32683,CG34393,CG42366,CG42450,CG42541,CG5160,CG5937,CG5953,CG9098,Cngl,Coop,Dh44-R1,dimm,DJ-1alpha,Dop2R,Drl-2,eag,erm,ewg,Fs,GABA-B-R3,Gadd45,gcl,Gpb5,Gyc32E,hiw,IMPPP,Ir21a,Ir25a,lb | amnesiac,Arrestin 1,Autophagy-related 9,bunched,Ca[2+]-channel protein alpha[[1]] subunit T,cacophony,Cholecystokinin-like receptor at 17D1,Cholecystokinin-like receptor at 17D3,Corepressor of Pangolin,Cyclic nucleotide-gated ion channel-like,Derailed 2,dimmed,Diuretic hormone 44 receptor 1,DJ-1alpha,Dopamine 2-like receptor,earmuff,erect wing,ether a go-go,Follistatin,germ cell-less,Glycoprotein hormone beta 5,Growth arrest and DNA damage-inducible 45,Guanyl cyclase at 32E,highwire,Immune |
| Over-expressed 48 AEL | 1301 | Biological process | sensory perception of chemical stimulus | GO:0007606 | 1.80E-07 | 3.64 | 29 | 94 | 10.47 | 30.9% | 2.2% | Ac78C,Arr1,Arr2,CheB42c,Desi,dpr11,dpr12,dpr16,dpr18,dpr2,dpr20,dpr4,dpr5,dpr6,eag,Gr33a,Gr39a,Gr94a,iav,Ir21a,Ir25a,Ir76b,Obp18a,Obp49a,Obp56d,Obp56h,Or82a,rdgB,Sh | Adenylyl cyclase 78C,Arrestin 1,Arrestin 2,Chemosensory protein B 42c,defective proboscis extension response 11,defective proboscis extension response 12,defective proboscis extension response 16,defective proboscis extension response 18,defective proboscis extension response 2,defective proboscis extension response 20,defective proboscis extension response 4,defective proboscis extension response 5,defective proboscis extension response 6,Desiccate,ether a go-go,Gustatory receptor 33a,Gustatory receptor 39a,Gustatory receptor |

| | | | | | | | | | | | | |
|---|---|---|---|---|---|---|---|---|---|---|---|---|
| Over-expressed 48 AEL | 1301 | Biological process | cell communication | GO:0007154 | 1.10E-06 | 1.5 | 223 | 1531 | 168.42 | 14.6% | 17.1% | amn,Arr1,Atg8a,bun,Ca-alpha1T,cac,CCKLR-17D1,CCKLR-17D3,CG10362,CG10702,CG13995,CG1657,CG16721,CG32547,CG32683,CG34393,CG42366,CG42450,CG42541,CG5160,CG5937,CG5953,CG9098,Cngl,Coop,Dh44-R1,dimm,DJ-1alpha,Dop2R,Drl-2,eag,erm,ewg,Fs,GABA-B-R3,Gadd45,gcl,Gpb5,Gyc32E,hiw,IMPPP,Ir21a,Ir25a,lb | amnesiac,Arrestin 1,Autophagy-related 8a,bunched,Ca[2+]-channel protein alpha[[1]] subunit T,cacophony,Cholecystokinin-like receptor at 17D1,Cholecystokinin-like receptor at 17D3,Corepressor of Pangolin,Cyclic nucleotide-gated ion channel-like,Derailed 2,dimmed,Diuretic hormone 44 receptor 1,DJ-1alpha,Dopamine 2-like receptor,earmuff,erect wing,ether a go-go,Follistatin,germ cell-less,Glycoprotein hormone beta 5,Growth arrest and DNA damage-inducible 45,Guanyl cyclase at 32E,highwire,Immune |
| Over-expressed 48 AEL | 1301 | Biological process | G protein-coupled receptor signaling pathway | GO:0007186 | 2.80E-06 | 2.55 | 39 | 165 | 18.26 | 23.6% | 3.0% | 5-HT1A,5-HT2B,Ac78C,amn,Arr1,Arr2,CCHa1,CCKLR-17D1,CCKLR-17D3,CG13579,CG13995,CG32547,CG34384,CG42450,CG43373,CG43795,CG7497,CNMaR,Dh44-R1,Dop1R1,Dop2R,Ets96B,foxo,GABA-B-R3,Galphaf,Gbeta76C,inaC,Lgr3,mGluR,MsR1,mthl8,ninaC,NPFR,Nplp4,Oamb,Octbeta1R,Octbeta3R,PK1-R,PK2-R2,Proc,rdgB,rdgC,Rh5,Rh6, | 5-hydroxytryptamine (serotonin) receptor 1A,5-hydroxytryptamine (serotonin) receptor 2B,Adenylyl cyclase 78C,amnesiac,Arrestin 1,Arrestin 2,CCHamide-1,Cholecystokinin-like receptor at 17D1,Cholecystokinin-like receptor at 17D3,CNMamide Receptor,Diuretic hormone 44 receptor 1,Dopamine 1-like receptor 1,Dopamine 2-like receptor,Ets96B,forkhead box, sub-group O,G protein alpha f subunit,G protein beta-subunit 76C,inactivation no afterpotential C,Leucine-rich repeat- |
| Over-expressed 48 AEL | 1301 | Biological process | antibacterial humoral response | GO:0019731 | 3.40E-06 | 5.18 | 16 | 41 | 4.57 | 39.0% | 1.2% | AttA,AttB,AttC,AttD,CecA1,CecA2,dnr1,DptB,Dro,Drs,IM3,IMPPP,Mtk,PGRP-LF,Rel,Thor | Attacin-A,Attacin-B,Attacin-C,Attacin-D,Cecropin A1,Cecropin A2,defense repressor 1,Diptericin B,Drosocin,Drosomycin,Immune induced molecule 3,Immune induced molecule prepropeptide,Metchnikowin,Peptidoglycan recognition protein LF,Relish,thor |

| | | | | | | | | | | | | | |
|---|---|---|---|---|---|---|---|---|---|---|---|---|---|
| Over-expressed 48 AEL | 1301 | Biological process | detection of light stimulus | GO:0009583 | 4.60E-06 | 4.42 | 18 | 51 | 5.68 | 35.3% | 1.4% | Arr1,Arr2,cac,Cerk,dlp,Gbeta76C,inaC,laza,ninaC,pinta,PIP5K59B,rdgB,rdgC,Rh5,Rh6,rtp,Sh,trpl | Arrestin 1,Arrestin 2,cacophony,Ceramide kinase,dally-like,G protein beta-subunit 76C,inactivation no afterpotential C,lazaro,neither inactivation nor afterpotential C,Phosphatidylinositol 4 phosphate 5-kinase at 59B,prolonged depolarization afterpotential (PDA) is not apparent,retinal degeneration B,retinal degeneration C,retinophilin,Rhodopsin 5,Rhodopsin 6,Shaker,transient receptor potential-like |
| Over-expressed 48 AEL | 1301 | Biological process | intestinal epithelial structure maintenance | GO:0060729 | 1.20E-05 | 48.16 | 6 | 7 | 0.78 | 85.7% | 0.5% | Atg1,Atg13,Atg17,Atg9,upd2,upd3 | Autophagy-related 1,Autophagy-related 13,Autophagy-related 17,Autophagy-related 9,unpaired 2,unpaired 3 |
| Over-expressed 48 AEL | 1301 | Biological process | detection of stimulus | GO:0051606 | 3.20E-05 | 4.08 | 16 | 48 | 5.3 | 33.3% | 1.2% | Ac78C,Arr1,Arr2,cac,Cerk,CheB42c,dlp,Gbeta76C,GNBP-like3,inaC,Ir21a,Ir25a,Ir62a,Ir76b,laza,ninaC,Or82a,pinta,PIP5K59B,prim,rdgB,rdgC,Rh5,Rh6,rtp,Sh,stj,trpl | Adenylyl cyclase 78C,Arrestin 1,Arrestin 2,cacophony,Ceramide kinase,Chemosensory protein B 42c,dally-like,G protein beta-subunit 76C,GNBP-like 3,inactivation no afterpotential C,Ionotropic receptor 21a,Ionotropic receptor 25a,Ionotropic receptor 62a,Ionotropic receptor 76b,lazaro,neither inactivation nor afterpotential C,Odorant receptor 82a,Phosphatidylinositol 4-phosphate 5-kinase at 59B,primacord,prolonged depolarization afterpotential (PDA) is not apparent,retinal degeneration B,retinal degeneration |
| Over-expressed 48 AEL | 1301 | Biological process | defense response to Gram-positive bacterium | GO:0050830 | 3.50E-05 | 4.04 | 16 | 48 | 5.35 | 33.3% | 1.2% | AttA,AttB,AttC,AttD,CecA1,CecA2,DptB,Dro,edin,Listericin,modSP,Mtk,PGRP-LF,PGRP-SB1,PGRP-SC1a,PGRP-SC1b | Attacin-A,Attacin-B,Attacin-C,Attacin-D,Cecropin A1,Cecropin A2,Diptericin B,Drosocin,elevated during infection,listericin,Metchnikowin,modular serine protease,Peptidoglycan recognition protein LF,Peptidoglycan recognition protein SB1,Peptidoglycan recognition protein SC1a,Peptidoglycan recognition protein SC1b |

| | | | | | | | | | | | | | |
|---|---|---|---|---|---|---|---|---|---|---|---|---|---|
| Over-expressed 48 AEL | 1301 | Biological process | mating behavior | GO:0007617 | 6.10E-05 | 2.67 | 26 | 105 | 11.7 | 24.8% | 2.0% | 5-HT1A,amn,cac,CG10433,dati,eag,Esp,FoxP,Gr33a,Gr39a,Gr94a,Ilp3,InR,lectin-46Ca,lectin-46Cb,Lnk,mGluR,Nlg2,Oamb,Obp56h,para,Pde1c,Sh,SIFaR,ss,to | 5-hydroxytryptamine (serotonin) receptor 1A,amnesiac,cacophony,datilografo,Epidermal stripes and patches,ether a go-go,Forkhead box P,Gustatory receptor 33a,Gustatory receptor 39a,Gustatory receptor 94a,Insulin-like peptide 3,Insulin-like receptor,lectin-46Ca,lectin-46Cb,Lnk,metabotropic Glutamate Receptor,Neuroligin 2,Octopamine receptor in mushroom bodies,Odorant-binding protein 56h,paralytic,Phosphodiesterase 1c,Shaker,SIFamide receptor,spineless,takeout,uncharacterized |
| Over-expressed 48 AEL | 1301 | Biological process | response to starvation | GO:0042594 | 8.60E-05 | 3.48 | 17 | 57 | 6.27 | 29.8% | 1.3% | Atg1,Atg17,Atg18a,Atg6,Atg8a,Atg9,Bruce,CG10924,chico,Crtc,Dop1R1,Ets96B,foxo,Gnmt,Iml1,ImpL2,InR,Lst,mio,mthl8,Myc,Oamb,p53,Pepck,Pi3K59F,Rel,REPTOR,REPTOR-BP,Rheb,Ric,rudhira,Slob,SNF4Agamma,Syx17,Thor,to,Wdr59 | Autophagy-related 1,Autophagy-related 17,Autophagy-related 18a,Autophagy-related 6,Autophagy-related 8a,Autophagy-related 9,BIR repeat containing ubiquitin-conjugating enzyme,chico,CREB-regulated transcription coactivator,Dopamine 1-like receptor 1,Ecdysone-inducible gene L2,Ets96B,forkhead box, sub-group O,Glycine N-methyltransferase,Increased minichromosome loss 1,Insulin-like receptor,Limostatin,methuselah-like 8,missing oocyte,Myc,Octopamine receptor in mushroom |
| Over-expressed 48 AEL | 1301 | Biological process | entry into reproductive diapause | GO:0055116 | 9.30E-05 | 40.09 | 5 | 6 | 0.67 | 83.3% | 0.4% | chico,foxo,Ilp3,ImpL2,InR | chico,Ecdysone-inducible gene L2,forkhead box, sub-group O,Insulin-like peptide 3,Insulin-like receptor |
| Over-expressed 48 AEL | 1301 | Biological process | anatomical structure homeostasis | GO:0060249 | 0.00013 | 2.24 | 33 | 153 | 17.04 | 21.6% | 2.5% | Arr1,Arr2,Atg1,Atg13,Atg17,Atg9,cac,cav,Cbp53E,chico,Csk,DJ-1alpha,Dop1R1,dysc,InR,Irbp,Mkp3,mre11,nAChRalpha6,Nap1,ninaC,p53,Pi3K59F,rad50,RanBPM,Smg5,Smg6,sqa,srp,Syx17,upd2,upd3,ver | Arrestin 1,Arrestin 2,Autophagy-related 1,Autophagy-related 13,Autophagy-related 17,Autophagy-related 9,C-terminal Src kinase,cacophony,Calbindin 53E,caravaggio,chico,DJ-1alpha,Dopamine 1-like receptor 1,dyschronic,Insulin-like receptor,Inverted repeat-binding protein,meiotic recombination 11,Mitogen-activated protein kinase phosphatase 3,neither inactivation nor afterpotential C,nicotinic Acetylcholine Receptor alpha6,Nucleosome assembly protein 1,p53,Phosphotidylinosi |

| | | | | | | | | | | | | | |
|---|---|---|---|---|---|---|---|---|---|---|---|---|---|
| Over-expressed 48 AEL | 1301 | Biological process | response to extracellular stimulus | GO:0009991 | 0.00017 | 3 | 19 | 71 | 7.81 | 26.8% | 1.5% | Atg1,Atg17,Atg18a,Atg6,Atg8a,Atg9,Bruce,CG10924,CG32206,CG34402,chico,Crtc,Dop1R1,Ets96B,foxo,Gnmt,Iml1,ImpL2,InR,Lst,mio,mthl8,Myc,Oamb,p53,Pepck,Pi3K59F,Rel,REPTOR,REPTOR-BP,Rheb,Ric,rudhira,Sh,Slob,SNF4Agamma,Syx17,Thor,to,Wdr59 | Autophagy-related 1,Autophagy-related 17,Autophagy-related 18a,Autophagy-related 6,Autophagy-related 8a,Autophagy-related 9,BIR repeat containing ubiquitin-conjugating enzyme,chico,CREB-regulated transcription coactivator,Dopamine 1-like receptor 1,Ecdysone-inducible gene L2,Ets96B,forkhead box, sub-group O,Glycine N-methyltransferase,Increased minichromosome loss 1,Insulin-like receptor,Limostatin,methuselah-like 8,missing oocyte,Myc,Octopamine receptor in mushroom |
| Over-expressed 48 AEL | 1301 | Biological process | cellular response to starvation | GO:0009267 | 0.00021 | 2.84 | 20 | 77 | 8.58 | 26.0% | 1.5% | Atg1,Atg17,Atg18a,Atg6,Atg8a,Atg9,Bruce,chico,foxo,Gnmt,Iml1,InR,Lst,mio,p53,Pi3K59F,Rel,SNF4Agamma,Syx17,Wdr59 | Autophagy-related 1,Autophagy-related 17,Autophagy-related 18a,Autophagy-related 6,Autophagy-related 8a,Autophagy-related 9,BIR repeat containing ubiquitin-conjugating enzyme,chico,forkhead box, sub-group O,Glycine N-methyltransferase,Increased minichromosome loss 1,Insulin-like receptor,Limostatin,missing oocyte,p53,Phosphotidylinositol 3 kinase 59F,Relish,SNF4/AMP-activated protein kinase gamma subunit,Syntaxin 17,WD repeat domain 59 |
| Over-expressed 48 AEL | 1301 | Biological process | cellular response to extracellular stimulus | GO:0031668 | 0.00022 | 2.74 | 21 | 83 | 9.25 | 25.3% | 1.6% | Atg1,Atg17,Atg18a,Atg6,Atg8a,Atg9,Bruce,chico,foxo,Gnmt,Iml1,InR,Lst,mio,Myc,p53,Pi3K59F,Rel,SNF4Agamma,Syx17,Wdr59 | Autophagy-related 1,Autophagy-related 17,Autophagy-related 18a,Autophagy-related 6,Autophagy-related 8a,Autophagy-related 9,BIR repeat containing ubiquitin-conjugating enzyme,chico,forkhead box, sub-group O,Glycine N-methyltransferase,Increased minichromosome loss 1,Insulin-like receptor,Limostatin,missing oocyte,Myc,p53,Phosphotidylinositol 3 kinase 59F,Relish,SNF4/AMP-activated protein kinase gamma subunit,Syntaxin 17,WD repeat domain 59 |

| | | | | | | | | | | | | | |
|---|---|---|---|---|---|---|---|---|---|---|---|---|---|
| Over-expressed 48 AEL | 1301 | Biological process | cellular response to stimulus | GO:0051716 | 0.00024 | 1.34 | 223 | 1653 | 181.85 | 13.5% | 17.1% | amn,Arr1,Atg18b,Atg8a,bun,cac,CCKLR-17D1,CCKLR-17D3,CG10362,CG10702,CG12896,CG13995,CG1657,CG16721,CG32547,CG32683,CG34393,CG42366,CG42450,CG42541,CG5160,CG5937,CG9003,CG9098,CG9272,Cngl,Coop,Cyp18a1,Cyp6g2,Dop2R,Drat,eag,erm,ewg,Fs,GABA-B-R3,Gadd45,gcl,Gpb5,Gyc32E,hiw,Irbp,lbm,Listericin,magu,mio,Mn | amnesiac,Arrestin 1,Autophagy-related 18b,Autophagy-related 8a,bunched,cacophony,Cholecystokinin-like receptor at 17D1,Cholecystokinin-like receptor at 17D3,Corepressor of Pangolin,Cyclic nucleotide-gated ion channel-like,Cyp6g2,Cytochrome P450-18a1,Death resistor Adh domain containing target,Dopamine 2-like receptor,earmuff,erect wing,ether a go-go,Follistatin,germ cell-less,Glycoprotein hormone beta 5,Growth arrest and DNA damage-inducible 45,Guanyl cyclase at 32E,highwire,Inverted |
| Over-expressed 48 AEL | 1301 | Biological process | regulation of membrane potential | GO:0042391 | 0.00026 | 3.24 | 16 | 56 | 6.21 | 28.6% | 1.2% | Ca-alpha1T,cac,eag,Ekar,Fife,GluClalpha,Grd,jus,nAChRalpha2,nAChRalpha3,nAChRalpha5,nAChRalpha6,nAChRalpha7,NaCP60E,para,Rdl,Sh,Shab,SK,SLO2,Slob | Ca[2+]-channel protein alpha[[1]] subunit T,cacophony,ether a go-go,Eye-enriched kainate receptor,fife,GluClalpha,Glycine receptor,julius seizure,Na channel protein 60E,nicotinic Acetylcholine Receptor alpha2,nicotinic Acetylcholine Receptor alpha3,nicotinic Acetylcholine Receptor alpha5,nicotinic Acetylcholine Receptor alpha6,nicotinic Acetylcholine Receptor alpha7,paralytic,Resistant to dieldrin,Shaker,Shaker cognate b,slowpoke 2,Slowpoke binding protein,small conductance calcium- |
| Over-expressed 48 AEL | 1301 | Biological process | protein complex oligomerization | GO:0051259 | 0.00032 | 3.53 | 14 | 46 | 5.12 | 30.4% | 1.1% | CG32206,CG34189,CG34384,CG34402,Crtc,eag,Gnmt,Hk,InR,PCNA2,Sh,Shab,Shaw,Trpm | CREB-regulated transcription coactivator,ether a go-go,Glycine N-methyltransferase,Hyperkinetic,Insulin-like receptor,Proliferating cell nuclear antigen 2,Shaker,Shaker cognate b,Shaker cognate w,Transient receptor potential cation channel, subfamily M,uncharacterized protein |
| Over-expressed 48 AEL | 1301 | Biological process | selective autophagy | GO:0061912 | 0.00035 | 8.03 | 7 | 14 | 1.56 | 50.0% | 0.5% | Atg1,Atg13,Atg17,Atg18a,Atg6,Atg9,ref(2)P | Autophagy-related 1,Autophagy-related 13,Autophagy-related 17,Autophagy-related 18a,Autophagy-related 6,Autophagy-related 9,refractory to sigma P |

| | | | | | | | | | | | | | |
|---|---|---|---|---|---|---|---|---|---|---|---|---|---|
| Over-expressed 48 AEL | 1301 | Biological process | second-messenger-mediated signaling | GO:0019932 | 0.00044 | 2.51 | 22 | 93 | 10.36 | 23.7% | 1.7% | Ac78C,amn,cac,CG16721,CG42260,CG43373,CG7497,Cngl,Dop1R1,Dop2R,Gyc32E,Gycbeta100B,inaC,Lgr3,Lst,Nos,Nox,Octbeta1R,Octbeta3R,rdgC,TpnC41C,trpl | Adenylyl cyclase 78C,amnesiac,cacophony,Cyclic nucleotide-gated ion channel-like,Dopamine 1-like receptor 1,Dopamine 2-like receptor,Guanyl cyclase at 32E,Guanylyl cyclase beta-subunit at 100B,inactivation no afterpotential C,Leucine-rich repeat-containing G protein-coupled receptor 3,Limostatin,NADPH oxidase,Nitric oxide synthase,Octopamine beta1 receptor,Octopamine beta3 receptor,retinal degeneration C,transient receptor potential-like,Troponin C at 41C,uncharacterized protein |
| Over-expressed 48 AEL | 1301 | Biological process | sensory perception of light stimulus | GO:0050953 | 0.00056 | 5.84 | 8 | 19 | 2.12 | 42.1% | 0.6% | cac,CG4025,dlp,dysc,inaC,Rh5,Rh6,trpl | cacophony,dally-like,dyschronic,inactivation no afterpotential C,Rhodopsin 5,Rhodopsin 6,transient receptor potential-like,uncharacterized protein |
| Over-expressed 48 AEL | 1301 | Biological process | dendrite self-avoidance | GO:0070593 | 0.00056 | 5.84 | 8 | 19 | 2.12 | 42.1% | 0.6% | bdl,CG44153,Dscam2,Dscam3,Dscam4,fry,ImpL2,Sema2b | borderless,Down syndrome cell adhesion molecule 2,Down syndrome cell adhesion molecule 3,Down syndrome cell adhesion molecule 4,Ecdysone-inducible gene L2,furry,Semaphorin 2b,uncharacterized protein |
| Over-expressed 48 AEL | 1301 | Biological process | detection of stimulus involved in sensory perception | GO:0050906 | 0.00059 | 3.72 | 12 | 38 | 4.23 | 31.6% | 0.9% | Ac78C,cac,dlp,inaC,Ir21a,Ir25a,Ir76b,Or82a,prim,rdgC,stj,trpl | Adenylyl cyclase 78C,cacophony,dally-like,inactivation no afterpotential C,Ionotropic receptor 21a,Ionotropic receptor 25a,Ionotropic receptor 76b,Odorant receptor 82a,primacord,retinal degeneration C,straightjacket,transient receptor potential-like |
| Over-expressed 48 AEL | 1301 | Biological process | autophagosome organization | GO:1905037 | 0.00059 | 4.02 | 11 | 33 | 3.68 | 33.3% | 0.8% | Atg1,Atg101,Atg13,Atg17,Atg18a,Atg18b,Atg6,Atg8a,Atg9,mv,Pi3K59F | Autophagy-related 1,Autophagy-related 101,Autophagy-related 13,Autophagy-related 17,Autophagy-related 18a,Autophagy-related 18b,Autophagy-related 6,Autophagy-related 8a,Autophagy-related 9,mauve,Phosphotidylinositol 3 kinase 59F |
| Over-expressed 48 AEL | 1301 | Biological process | tissue regeneration | GO:0042246 | 6.00E-04 | 7.02 | 7 | 15 | 1.67 | 46.7% | 0.5% | Alr,bun,Idgf1,p53,Spn88Eb,Sxl,Top3alpha | Augmenter of liver regeneration,bunched,Imaginal disc growth factor 1,p53,Serpin 88Eb,Sex lethal,Topoisomerase 3alpha |

| | | | | | | | | | | | | | |
|---|---|---|---|---|---|---|---|---|---|---|---|---|---|
| Over-expressed 48 AEL | 1301 | Biological process | adaptation of signaling pathway | GO:0023058 | 7.00E-04 | 32.04 | 4 | 5 | 0.56 | 80.0% | 0.3% | Arr1,Arr2,inaC,ninaC | Arrestin 1,Arrestin 2,inactivation no afterpotential C,neither inactivation nor afterpotential C |
| Over-expressed 48 AEL | 1301 | Biological process | IMP metabolic process | GO:0046040 | 7.00E-04 | 32.04 | 4 | 5 | 0.56 | 80.0% | 0.3% | ade2,ade5,AdSS,AMPdeam | ade5,adenosine 2,Adenylosuccinate Synthetase,AMP deaminase |
| Over-expressed 48 AEL | 1301 | Biological process | phototransduction, visible light | GO:0007603 | 7.00E-04 | 4.82 | 9 | 24 | 2.67 | 37.5% | 0.7% | Arr1,Arr2,Gbeta76C,inaC,ninaC,PIP5K59B,rdgB,rdgC,trpl | Arrestin 1,Arrestin 2,G protein beta-subunit 76C,inactivation no afterpotential C,neither inactivation nor afterpotential C,Phosphatidylinositol 4 phosphate 5-kinase at 59B,retinal degeneration B,retinal degeneration C,transient receptor potential-like |
| Over-expressed 48 AEL | 1301 | Biological process | epithelial structure maintenance | GO:0010669 | 7.00E-04 | 4.82 | 9 | 24 | 2.67 | 37.5% | 0.7% | Atg1,Atg13,Atg17,Atg9,Csk,Mkp3,Pi3K59F,upd2,upd3 | Autophagy-related 1,Autophagy-related 13,Autophagy-related 17,Autophagy-related 9,C-terminal Src kinase,Mitogen-activated protein kinase phosphatase 3,Phosphotidylinositol 3 kinase 59F,unpaired 2,unpaired 3 |
| Over-expressed 48 AEL | 1301 | Biological process | action potential | GO:0001508 | 0.00071 | 13.36 | 5 | 8 | 0.89 | 62.5% | 0.4% | Ca-alpha1T,Grd,NaCP60E,Sh,Shab | Ca[2+]-channel protein alpha[[1]] subunit T,Glycine receptor,Na channel protein 60E,Shaker,Shaker cognate b |
| Over-expressed 48 AEL | 1301 | Biological process | response to light intensity | GO:0009642 | 0.00071 | 13.36 | 5 | 8 | 0.89 | 62.5% | 0.4% | Arr2,Ekar,inaC,laza,ninaC | Arrestin 2,Eye-enriched kainate receptor,inactivation no afterpotential C,lazaro,neither inactivation nor afterpotential C |
| Over-expressed 48 AEL | 1301 | Biological process | dormancy process | GO:0022611 | 0.00071 | 13.36 | 5 | 8 | 0.89 | 62.5% | 0.4% | chico,foxo,Ilp3,ImpL2,InR | chico,Ecdysone-inducible gene L2,forkhead box, sub-group O,Insulin-like peptide 3,Insulin-like receptor |
| Over-expressed 48 AEL | 1301 | Biological process | cellular response to light stimulus | GO:0071482 | 0.00079 | 3.85 | 11 | 34 | 3.79 | 32.4% | 0.8% | Arr1,Arr2,Ekar,Gbeta76C,inaC,ninaC,PIP5K59B,rdgB,rdgC,Rh5,Rh6 | Arrestin 1,Arrestin 2,Eye-enriched kainate receptor,G protein beta-subunit 76C,inactivation no afterpotential C,neither inactivation nor afterpotential C,Phosphatidylinositol 4 phosphate 5-kinase at 59B,retinal degeneration B,retinal degeneration C,Rhodopsin 5,Rhodopsin 6 |

| | | | | | | | | | | | | | |
|---|---|---|---|---|---|---|---|---|---|---|---|---|---|
| Over-expressed 48 AEL | 1301 | Biological process | chemosensory behavior | GO:0007635 | 0.00085 | 2.56 | 19 | 79 | 8.8 | 24.1% | 1.5% | Alk,amn,CG13377,chico,Dop1R1,drl,futsch,Gr33a,Gr39a,Gr94a,NaCP60E,NPFR,Oamb,Obp56d,Obp56h,Pino,rad,Rdl,Sh | amnesiac,Anaplastic lymphoma kinase,chico,derailed,Dopamine 1-like receptor 1,futsch,Gustatory receptor 33a,Gustatory receptor 39a,Gustatory receptor 94a,Na channel protein 60E,Neuropeptide F receptor,Octopamine receptor in mushroom bodies,Odorant-binding protein 56d,Odorant-binding protein 56h,Pinocchio,radish,Resistant to dieldrin,Shaker,uncharacterized protein |
| Over-expressed 48 AEL | 1301 | Biological process | deactivation of rhodopsin mediated signaling | GO:0016059 | 0.00096 | 6.24 | 7 | 16 | 1.78 | 43.8% | 0.5% | Arr1,Arr2,Gbeta76C,inaC,ninaC,rdgB,rdgC | Arrestin 1,Arrestin 2,G protein beta-subunit 76C,inactivation no afterpotential C,neither inactivation nor afterpotential C,retinal degeneration B,retinal degeneration C |
| Over-expressed 48 AEL | 1301 | Biological process | regulation of G protein-coupled receptor signaling pathway | GO:0008277 | 0.00099 | 4.52 | 9 | 25 | 2.78 | 36.0% | 0.7% | Arr1,Arr2,Ets96B,foxo,Gbeta76C,inaC,ninaC,rdgB,rdgC | Arrestin 1,Arrestin 2,Ets96B,forkhead box, sub-group O,G protein beta-subunit 76C,inactivation no afterpotential C,neither inactivation nor afterpotential C,retinal degeneration B,retinal degeneration C |
| Over-expressed 48 AEL | 1301 | Biological process | larval midgut cell programmed cell death | GO:0035096 | 0.00099 | 4.52 | 9 | 25 | 2.78 | 36.0% | 0.7% | Atg1,Atg101,Atg13,Atg17,Atg18a,Atg6,Atg8a,Atg9,Pi3K59F | Autophagy-related 1,Autophagy-related 101,Autophagy-related 13,Autophagy-related 17,Autophagy-related 18a,Autophagy-related 6,Autophagy-related 8a,Autophagy-related 9,Phosphotidylinositol 3 kinase 59F |
| Over-expressed 48 AEL | 1301 | Molecular function | molecular transducer activity | GO:0060089 | 3.20E-11 | 2.75 | 71 | 291 | 31.96 | 24.4% | 5.5% | 5-HT1A,5-HT2B,Alk,CCKLR-17D1,CCKLR-17D3,CG10702,CG13579,CG13995,CG31760,CG32206,CG32547,CG34402,CG43795,CG44153,CG7497,CNMaR,Dh44-R1,Dop1R1,Dop2R,drl,Drl-2,Ekar,ftz-f1,GABA-B-R3,GluClalpha,GluRIA,GNBP2,Gr33a,Gr39a,Gr94a,Grd,Gyc32E,InR,Ir21a,Ir25a,Ir76b,kek6,Lgr3,Met,mGluR,MsR1,mthl8,nAC | 5-hydroxytryptamine (serotonin) receptor 1A,5-hydroxytryptamine (serotonin) receptor 2B,Anaplastic lymphoma kinase,Cholecystokinin-like receptor at 17D1,Cholecystokinin-like receptor at 17D3,CNMamide Receptor,derailed,Derailed 2,Diuretic hormone 44 receptor 1,Dopamine 1-like receptor 1,Dopamine 2-like receptor,Eye-enriched kainate receptor,ftz transcription factor 1,GluClalpha,Glutamate receptor IA,Glycine receptor,Gram-negative bacteria binding protein |

| | | | | | | | | | | | | | |
|---|---|---|---|---|---|---|---|---|---|---|---|---|---|
| Over-expressed 48 AEL | 1301 | Molecular function | G protein-coupled receptor activity | GO:0004930 | 1.10E-08 | 3.92 | 32 | 100 | 10.98 | 32.0% | 2.5% | 5-HT1A,5-HT2B,CCKLR-17D1,CCKLR-17D3,CG13579,CG13995,CG31760,CG32547,CG43795,CG44153,CG7497,CNMaR,Dh44-R1,Dop1R1,Dop2R,GABA-B-R3,Lgr3,mGluR,MsR1,mthl8,NPFR,Oamb,Octbeta1R,Octbeta3R,PK1-R,PK2-R2,Rh5,Rh6,RYa-R,SIFaR,smog,TkR86C | 5-hydroxytryptamine (serotonin) receptor 1A,5-hydroxytryptamine (serotonin) receptor 2B,Cholecystokinin-like receptor at 17D1,Cholecystokinin-like receptor at 17D3,CNMamide Receptor,Diuretic hormone 44 receptor 1,Dopamine 1-like receptor 1,Dopamine 2-like receptor,Leucine-rich repeat-containing G protein-coupled receptor 3,metabotropic GABA-B receptor subtype 3,metabotropic Glutamate Receptor,methuselah-like 8,Myosuppressin receptor 1,Neuropeptide F |
| Over-expressed 48 AEL | 1301 | Molecular function | ion gated channel activity | GO:0022839 | 1.90E-08 | 3.91 | 31 | 97 | 10.65 | 32.0% | 2.4% | Ca-alpha1T,cac,CG15270,CG42260,CG4587,Cngl,eag,Ekar,GluClalpha,GluRIA,Grd,Hk,Ir21a,Ir25a,Ir62a,Ir76b,nAChRalpha2,nAChRalpha3,nAChRalpha5,nAChRalpha6,nAChRalpha7,NaCP60E,Nmdar2,Rdl,Sh,Shab,Shaw,SK,SLO2,stj,Trpm | Ca[2+]-channel protein alpha[[1]] subunit T,cacophony,Cyclic nucleotide-gated ion channel-like,ether a go-go,Eye-enriched kainate receptor,GluClalpha,Glutamate receptor IA,Glycine receptor,Hyperkinetic,Ionotropic receptor 21a,Ionotropic receptor 25a,Ionotropic receptor 62a,Ionotropic receptor 76b,Na channel protein 60E,nicotinic Acetylcholine Receptor alpha2,nicotinic Acetylcholine Receptor alpha3,nicotinic Acetylcholine Receptor alpha5,nicotinic Acetylcholine Receptor alpha6,nicotinic Acetylcholine Receptor |
| Over-expressed 48 AEL | 1301 | Molecular function | neurotransmitter receptor activity | GO:0030594 | 2.30E-08 | 5.51 | 22 | 55 | 6.04 | 40.0% | 1.7% | 5-HT1A,5-HT2B,CG43795,Dop1R1,Dop2R,Ekar,GABA-B-R3,GluRIA,Grd,Ir21a,Ir25a,Ir76b,nAChRalpha2,nAChRalpha3,nAChRalpha5,nAChRalpha6,nAChRalpha7,Nmdar2,Oamb,Octbeta1R,Octbeta3R,Rdl | 5-hydroxytryptamine (serotonin) receptor 1A,5-hydroxytryptamine (serotonin) receptor 2B,Dopamine 1-like receptor 1,Dopamine 2-like receptor,Eye-enriched kainate receptor,Glutamate receptor IA,Glycine receptor,Ionotropic receptor 21a,Ionotropic receptor 25a,Ionotropic receptor 76b,metabotropic GABA-B receptor subtype 3,nicotinic Acetylcholine Receptor alpha2,nicotinic Acetylcholine Receptor alpha3,nicotinic Acetylcholine Receptor alpha5,nicotinic Acetylcholine Receptor alpha6,nicotinic |

| | | | | | | | | | | | | | |
|---|---|---|---|---|---|---|---|---|---|---|---|---|---|
| Over-expressed 48 AEL | 1301 | Molecular function | ion channel activity | GO:0005216 | 7.10E-08 | 3.11 | 38 | 140 | 15.38 | 27.1% | 2.9% | Ca-alpha1T,cac,CG15270,CG1688,CG42260,CG4587,Cngl,eag,Ekar,GluClalpha,GluRIA,Grd,Hk,iav,Ir21a,Ir25a,Ir62a,Ir76b,nAChRalpha2,nAChRalpha3,nAChRalpha5,nAChRalpha6,nAChRalpha7,NaCP60E,Nmdar2,para,Rdl,Sh,Shab,Shaw,SK,SLO2,stj,Task6,trpl,Trpm,unc80,wtrw | Ca[2+]-channel protein alpha[[1]] subunit T,cacophony,Cyclic nucleotide-gated ion channel-like,ether a go-go,Eye-enriched kainate receptor,GluClalpha,Glutamate receptor IA,Glycine receptor,Hyperkinetic,inactive,Ionotropic receptor 21a,Ionotropic receptor 25a,Ionotropic receptor 62a,Ionotropic receptor 76b,Na channel protein 60E,nicotinic Acetylcholine Receptor alpha2,nicotinic Acetylcholine Receptor alpha3,nicotinic Acetylcholine Receptor alpha5,nicotinic Acetylcholine Receptor alpha6,nicotinic |
| Over-expressed 48 AEL | 1301 | Molecular function | channel activity | GO:0015267 | 5.90E-07 | 2.8 | 38 | 151 | 16.59 | 25.2% | 2.9% | Ca-alpha1T,cac,CG15270,CG1688,CG42260,CG4587,Cngl,eag,Ekar,GluClalpha,GluRIA,Grd,Hk,iav,Ir21a,Ir25a,Ir62a,Ir76b,nAChRalpha2,nAChRalpha3,nAChRalpha5,nAChRalpha6,nAChRalpha7,NaCP60E,Nmdar2,para,Rdl,Sh,Shab,Shaw,SK,SLO2,stj,Task6,trpl,Trpm,unc80,wtrw | Ca[2+]-channel protein alpha[[1]] subunit T,cacophony,Cyclic nucleotide-gated ion channel-like,ether a go-go,Eye-enriched kainate receptor,GluClalpha,Glutamate receptor IA,Glycine receptor,Hyperkinetic,inactive,Ionotropic receptor 21a,Ionotropic receptor 25a,Ionotropic receptor 62a,Ionotropic receptor 76b,Na channel protein 60E,nicotinic Acetylcholine Receptor alpha2,nicotinic Acetylcholine Receptor alpha3,nicotinic Acetylcholine Receptor alpha5,nicotinic Acetylcholine Receptor alpha6,nicotinic |
| Over-expressed 48 AEL | 1301 | Molecular function | transmitter-gated ion channel activity | GO:0022824 | 1.50E-05 | 5.61 | 13 | 32 | 3.51 | 40.6% | 1.0% | Ekar,GluRIA,Grd,Ir21a,Ir25a,Ir76b,nAChRalpha2,nAChRalpha3,nAChRalpha5,nAChRalpha6,nAChRalpha7,Nmdar2,Rdl | Eye-enriched kainate receptor,Glutamate receptor IA,Glycine receptor,Ionotropic receptor 21a,Ionotropic receptor 25a,Ionotropic receptor 76b,nicotinic Acetylcholine Receptor alpha2,nicotinic Acetylcholine Receptor alpha3,nicotinic Acetylcholine Receptor alpha5,nicotinic Acetylcholine Receptor alpha6,nicotinic Acetylcholine Receptor alpha7,NMDA receptor 2,Resistant to dieldrin |

| | | | | | | | | | | | | | |
|---|---|---|---|---|---|---|---|---|---|---|---|---|---|
| Over-expressed 48 AEL | 1301 | Molecular function | ligand-gated channel activity | GO:0022834 | 1.70E-05 | 3.9 | 18 | 56 | 6.15 | 32.1% | 1.4% | CG42260,Cngl,Ekar,GluClalpha,GluRIA,Grd,Ir21a,Ir25a,Ir62a,Ir76b,nAChRalpha2,nAChRalpha3,nAChRalpha5,nAChRalpha6,nAChRalpha7,Nmdar2,Rdl,Trpm | Cyclic nucleotide-gated ion channel-like,Eye-enriched kainate receptor,GluClalpha,Glutamate receptor IA,Glycine receptor,Ionotropic receptor 21a,Ionotropic receptor 25a,Ionotropic receptor 62a,Ionotropic receptor 76b,nicotinic Acetylcholine Receptor alpha2,nicotinic Acetylcholine Receptor alpha3,nicotinic Acetylcholine Receptor alpha5,nicotinic Acetylcholine Receptor alpha6,nicotinic Acetylcholine Receptor alpha7,NMDA receptor 2,Resistant to dieldrin,Transient receptor potential cation channel, subfamily |
| Over-expressed 48 AEL | 1301 | Molecular function | cation transmembrane transporter activity | GO:0008324 | 2.30E-05 | 2.02 | 52 | 267 | 29.33 | 19.5% | 4.0% | Ca-alpha1T,cac,CG10804,CG14606,CG15270,CG1628,CG1688,CG3285,CG33282,CG33310,CG42260,CG43066,CG4476,CG4587,CG5646,Cngl,Eaat2,eag,Ekar,GluRIA,Grd,Hk,iav,Ir25a,Ir76b,List,MME1,nAChRalpha2,nAChRalpha3,nAChRalpha5,nAChRalpha6,nAChRalpha7,NaCP60E,Nhe3,NKCC,Nmdar2,para,rdgB,Sh,Shab,Shaw,SK,SLO2,stj,Task6,Tret1- | Ca[2+]-channel protein alpha[[1]] subunit T,cacophony,Cyclic nucleotide-gated ion channel-like,ether a go-go,Excitatory amino acid transporter 2,Eye-enriched kainate receptor,Glutamate receptor IA,Glycine receptor,Hyperkinetic,inactive,Ionotropic receptor 25a,Ionotropic receptor 76b,Lithium-inducible SLC6 transporter,Mitochondrial Magnesium Exporter 1,Na channel protein 60E,Na[+]/H[+] hydrogen exchanger 3,nicotinic Acetylcholine Receptor alpha2,nicotinic Acetylcholine Receptor alpha3,nicotinic Acetylcholine Receptor |
| Over-expressed 48 AEL | 1301 | Molecular function | voltage-gated cation channel activity | GO:0022843 | 2.50E-05 | 5.78 | 12 | 29 | 3.19 | 41.4% | 0.9% | Ca-alpha1T,cac,CG4587,eag,Hk,NaCP60E,Nmdar2,Sh,Shab,Shaw,SLO2,stj | Ca[2+]-channel protein alpha[[1]] subunit T,cacophony,ether a go-go,Hyperkinetic,Na channel protein 60E,NMDA receptor 2,Shaker,Shaker cognate b,Shaker cognate w,slowpoke 2,straightjacket,uncharacterized protein |

| | | | | | | | | | | | | | |
|---|---|---|---|---|---|---|---|---|---|---|---|---|---|
| Over-expressed 48 AEL | 1301 | Molecular function | metal ion transmembrane transporter activity | GO:0046873 | 8.00E-05 | 2.27 | 34 | 158 | 17.35 | 21.5% | 2.6% | Ca-alpha1T,cac,CG10804,CG1688,CG33310,CG43066,CG4476,CG4587,Eaat2,eag,Ekar,GluRIA,Hk,iav,Ir76b,List,MME1,NaCP60E,Nhe3,NKCC,para,rdgB,Sh,Shab,Shaw,SK,SLO2,stj,Task6,trpl,Trpm,wtrw,Zip99C,ZnT35C | Ca[2+]-channel protein alpha[[1]] subunit T,cacophony,ether a go-go,Excitatory amino acid transporter 2,Eye-enriched kainate receptor,Glutamate receptor IA,Hyperkinetic,inactive,Ionotropic receptor 76b,Lithium-inducible SLC6 transporter,Mitochondrial Magnesium Exporter 1,Na channel protein 60E,Na[+]/H[+] hydrogen exchanger 3,paralytic,retinal degeneration B,Shaker,Shaker cognate b,Shaker cognate w,slowpoke 2,small conductance calcium-activated potassium channel,sodium |
| Over-expressed 48 AEL | 1301 | Molecular function | excitatory extracellular ligand-gated ion channel activity | GO:0005231 | 9.10E-05 | 11.43 | 7 | 12 | 1.32 | 58.3% | 0.5% | GluClalpha,Grd,nAChRalpha2,nAChRalpha3,nAChRalpha5,nAChRalpha6,nAChRalpha7 | GluClalpha,Glycine receptor,nicotinic Acetylcholine Receptor alpha2,nicotinic Acetylcholine Receptor alpha3,nicotinic Acetylcholine Receptor alpha5,nicotinic Acetylcholine Receptor alpha6,nicotinic Acetylcholine Receptor alpha7 |
| Over-expressed 48 AEL | 1301 | Molecular function | neuropeptide receptor activity | GO:0008188 | 0.00012 | 3.96 | 14 | 43 | 4.72 | 32.6% | 1.1% | CCKLR-17D1,CCKLR-17D3,CG13995,CG32547,CNMaR,Dh44-R1,Lgr3,MsR1,NPFR,PK1-R,PK2-R2,RYa-R,SIFaR,TkR86C | Cholecystokinin-like receptor at 17D1,Cholecystokinin-like receptor at 17D3,CNMamide Receptor,Diuretic hormone 44 receptor 1,Leucine-rich repeat-containing G protein-coupled receptor 3,Myosuppressin receptor 1,Neuropeptide F receptor,Pyrokinin 1 receptor,Pyrokinin 2 receptor 2,RYamide receptor,SIFamide receptor,Tachykinin-like receptor at 86C,uncharacterized protein |
| Over-expressed 48 AEL | 1301 | Molecular function | ligand-gated cation channel activity | GO:0099094 | 0.00022 | 3.94 | 13 | 40 | 4.39 | 32.5% | 1.0% | CG42260,Cngl,Ekar,GluRIA,Ir25a,Ir76b,nAChRalpha2,nAChRalpha3,nAChRalpha5,nAChRalpha6,nAChRalpha7,Nmdar2,Trpm | Cyclic nucleotide-gated ion channel-like,Eye-enriched kainate receptor,Glutamate receptor IA,Ionotropic receptor 25a,Ionotropic receptor 76b,nicotinic Acetylcholine Receptor alpha2,nicotinic Acetylcholine Receptor alpha3,nicotinic Acetylcholine Receptor alpha5,nicotinic Acetylcholine Receptor alpha6,nicotinic Acetylcholine Receptor alpha7,NMDA receptor 2,Transient receptor potential cation channel, subfamily M,uncharacterized protein |

| | | | | | | | | | | | | | |
|---|---|---|---|---|---|---|---|---|---|---|---|---|---|
| Over-expressed 48 AEL | 1301 | Molecular function | transmembrane signaling receptor activity | GO:0004888 | 0.00024 | 2.51 | 24 | 105 | 11.26 | 22.9% | 1.8% | 5-HT1A,5-HT2B,Alk,CCKLR-17D1,CCKLR-17D3,CG10702,CG13579,CG13995,CG31760,CG32547,CG43795,CG44153,CG7497,CNMaR,Dh44-R1,Dop1R1,Dop2R,drl,Drl-2,Ekar,GABA-B-R3,GluClalpha,GluRIA,Gr33a,Gr39a,Gr94a,Grd,InR,Ir21a,Ir25a,Ir76b,Lgr3,mGluR,MsR1,mthl8,nAChRalpha2,nAChRalpha3,nAChRalpha5,nAChRalpha6,n | 5-hydroxytryptamine (serotonin) receptor 1A,5-hydroxytryptamine (serotonin) receptor 2B,Anaplastic lymphoma kinase,Cholecystokinin-like receptor at 17D1,Cholecystokinin-like receptor at 17D3,CNMamide Receptor,derailed,Derailed 2,Diuretic hormone 44 receptor 1,Dopamine 1-like receptor 1,Dopamine 2-like receptor,Eye-enriched kainate receptor,GluClalpha,Glutamate receptor IA,Glycine receptor,Gustatory receptor 33a,Gustatory receptor 39a,Gustatory receptor 94a,Insulin- |
| Over-expressed 48 AEL | 1301 | Molecular function | cell-cell adhesion mediator activity | GO:0098632 | 0.00025 | 12.23 | 6 | 10 | 1.1 | 60.0% | 0.5% | bdl,CG44153,Dscam2,Dscam3,Dscam4,ImpL2 | borderless,Down syndrome cell adhesion molecule 2,Down syndrome cell adhesion molecule 3,Down syndrome cell adhesion molecule 4,Ecdysone-inducible gene L2,uncharacterized protein |
| Over-expressed 48 AEL | 1301 | Molecular function | voltage-gated channel activity | GO:0022832 | 0.00029 | 4.09 | 12 | 36 | 3.95 | 33.3% | 0.9% | Ca-alpha1T,cac,CG4587,eag,Hk,NaCP60E,Nmdar2,Sh,Shab,Shaw,SLO2,stj | Ca[2+]-channel protein alpha[[1]] subunit T,cacophony,ether a go-go,Hyperkinetic,Na channel protein 60E,NMDA receptor 2,Shaker,Shaker cognate b,Shaker cognate w,slowpoke 2,straightjacket,uncharacterized protein |
| Over-expressed 48 AEL | 1301 | Molecular function | ATP-dependent microtubule motor activity, minus-end-directed | GO:0008569 | 0.00049 | 9.78 | 6 | 11 | 1.21 | 54.5% | 0.5% | CG3339,CG9492,Dhc62B,Dhc93AB,Dhc98D,Kif3C | Dynein heavy chain at 62B,Dynein heavy chain at 89D,Dynein heavy chain at 93AB,Kinesin family member 3C,uncharacterized protein |
| Over-expressed 48 AEL | 1301 | Molecular function | peptide receptor activity | GO:0001653 | 0.00071 | 3.19 | 14 | 50 | 5.49 | 28.0% | 1.1% | CCKLR-17D1,CCKLR-17D3,CG13995,CG32547,CNMaR,Dh44-R1,Lgr3,MsR1,NPFR,PK1-R,PK2-R2,RYa-R,SIFaR,TkR86C | Cholecystokinin-like receptor at 17D1,Cholecystokinin-like receptor at 17D3,CNMamide Receptor,Diuretic hormone 44 receptor 1,Leucine-rich repeat-containing G protein-coupled receptor 3,Myosuppressin receptor 1,Neuropeptide F receptor,Pyrokinin 1 receptor,Pyrokinin 2 receptor 2,RYamide receptor,SIFamide receptor,Tachykinin-like receptor at 86C,uncharacterized protein |
| Over-expressed 48 AEL | 1301 | Molecular function | telomeric DNA binding | GO:0042162 | 0.00089 | 8.15 | 6 | 12 | 1.32 | 50.0% | 0.5% | cav,Irbp,rad50,Smg5,Smg6,ver | caravaggio,Inverted repeat-binding protein,rad50,Smg5,Smg6,verrocchio |

| | | | | | | | | | | | | | |
|---|---|---|---|---|---|---|---|---|---|---|---|---|---|
| Over-expressed 48 AEL | 1301 | Cellular component | cell projection | GO:0042995 | 1.70E-15 | 2.82 | 100 | 423 | 44.87 | 23.6% | 7.7% | 5-HT1A,Alk,ana1,Arr1,Arr2,Atg8a,BBS4,bdl,Burs,Cbp53E,Cc2d2a,CCKLR-17D1,Cep135,cep290,CG10702,CG1126,CG15666,CG15923,CG16719,CG17387,CG32206,CG3339,CG34402,CG42366,CG44153,CG5142,CG9492,ChAT,CLIP-190,cmpy,Dab,Dhc62B,Dhc93AB,Dhc98D,dia,Dop1R1,dpr11,dpr12,dpr18,dpr2,dpr20,dpr4,dpr5,dpr6,drl,Dscam2, | 5-hydroxytryptamine (serotonin) receptor 1A,Anaplastic lymphoma kinase,anastral spindle 1,Arrestin 1,Arrestin 2,Autophagy-related 8a,Bardet-Biedl syndrome 4,borderless,Bursicon,Calbindin 53E,Centrosomal protein 135kDa,cep290,Cholecystokinin-like receptor at 17D1,Choline acetyltransferase,Coiled-coil and C2 domain containing 2A,crimpy,Cytoplasmic linker protein 190,defective proboscis extension response 11,defective proboscis extension response 12,defective proboscis |
| Over-expressed 48 AEL | 1301 | Cellular component | integral component of plasma membrane | GO:0005887 | 2.80E-15 | 2.86 | 96 | 401 | 42.53 | 23.9% | 7.4% | 5-HT1A,Ac78C,Alk,Atet,bdl,beat-Ic,Ca-alpha1T,CadN2,CCKLR-17D1,CG10702,CG11425,CG13404,CG14606,CG1688,CG31814,CG32206,CG3285,CG33282,CG33310,CG34402,CG43373,CG44325,Dh44-R1,DIP-beta,DIP-delta,DIP-eta,Dop1R1,Dop2R,dpr11,dpr18,dpr2,dpr20,dpr5,drl,Drl-2,Dscam2,Dscam3,Dscam4, | 5-hydroxytryptamine (serotonin) receptor 1A,ABC transporter expressed in trachea,Adenylyl cyclase 78C,Anaplastic lymphoma kinase,beaten path Ic,borderless,Ca[2+]-channel protein alpha[[1]] subunit T,Cadherin-N2,Cholecystokinin-like receptor at 17D1,defective proboscis extension response 11,defective proboscis extension response 18,defective proboscis extension response 2,defective proboscis extension response 20,defective proboscis extension response 5,derailed,Derailed |
| Over-expressed 48 AEL | 1301 | Cellular component | plasma membrane part | GO:0044459 | 1.60E-10 | 2.05 | 123 | 668 | 70.86 | 18.4% | 9.5% | 5-HT1A,Ac78C,Atet,bdl,beat-Ic,Ca-alpha1T,cac,CadN2,CCKLR-17D1,CG10702,CG11425,CG13404,CG14606,CG1688,CG31814,CG32206,CG3285,CG33282,CG33310,CG34402,CG43373,CG44325,CG4587,Dh44-R1,dia,DIP-beta,DIP-eta,dlp,Dop1R1,Dop2R,dpr11,dpr18,dpr2,dpr20,dpr4,dpr5,Drl-2,Dscam2,Dscam4,Dyb,eag, | 5-hydroxytryptamine (serotonin) receptor 1A,ABC transporter expressed in trachea,Adenylyl cyclase 78C,beaten path Ic,borderless,Ca[2+]-channel protein alpha[[1]] subunit T,cacophony,Cadherin-N2,Cholecystokinin-like receptor at 17D1,dally-like,defective proboscis extension response 11,defective proboscis extension response 18,defective proboscis extension response 2,defective proboscis extension response 20,defective proboscis extension response 4,defective proboscis extension response 5,Derailed |

| | | | | | | | | | | | | | |
|---|---|---|---|---|---|---|---|---|---|---|---|---|---|
| Over-expressed 48 AEL | 1301 | Cellular component | intrinsic component of membrane | GO:0031224 | 3.80E-10 | 1.81 | 175 | 1071 | 113.6 | 16.3% | 13.5% | 5-HT1A,Atet,Atg9,bdl,beat-Ic,bwa,Ca-alpha1T,cac,CadN2,CCKLR-17D1,CCKLR-17D3,CG10702,CG10804,CG13404,CG13579,CG13995,CG14439,CG15098,CG1572,CG15760,CG1688,CG18609,CG18635,CG2975,CG31272,CG31321,CG31760,CG31814,CG32547,CG3285,CG33282,CG33310,CG33474,CG34120,CG42260,CG43066,CG44325,CG4587, | 5-hydroxytryptamine (serotonin) receptor 1A,ABC transporter expressed in trachea,Autophagy-related 9,beaten path Ic,borderless,brain washing,Ca[2+]-channel protein alpha[[1]] subunit T,cacophony,Cadherin-N2,Cholecystokinin-like receptor at 17D1,Cholecystokinin-like receptor at 17D3,Cyclic nucleotide-gated ion channel-like,Cytochrome b5-related,defective proboscis extension response 18,defective proboscis extension response 2,Derailed 2,Diuretic hormone 44 receptor 1,Dopamine 2-like receptor,Down |
| Over-expressed 48 AEL | 1301 | Cellular component | transmembrane transporter complex | GO:1902495 | 3.40E-09 | 4.57 | 29 | 84 | 8.91 | 34.5% | 2.2% | Ca-alpha1T,cac,CG32206,CG33310,CG34402,CG42260,CG4587,Cngl,eag,GluRIA,Hk,iav,Ir25a,Ir76b,nAChRalpha2,nAChRalpha3,nAChRalpha5,nAChRalpha6,nAChRalpha7,NaCP60E,Nmdar2,para,Sh,Shab,Shaw,trpl,Trpm,unc80,wtrw | Ca[2+]-channel protein alpha[[1]] subunit T,cacophony,Cyclic nucleotide-gated ion channel-like,ether a go-go,Glutamate receptor IA,Hyperkinetic,inactive,Ionotropic receptor 25a,Ionotropic receptor 76b,Na channel protein 60E,nicotinic Acetylcholine Receptor alpha2,nicotinic Acetylcholine Receptor alpha3,nicotinic Acetylcholine Receptor alpha5,nicotinic Acetylcholine Receptor alpha6,nicotinic Acetylcholine Receptor alpha7,NMDA receptor 2,paralytic,Shaker,Shaker cognate b,Shaker cognate w,Transient receptor potential cation channel, |
| Over-expressed 48 AEL | 1301 | Cellular component | cation channel complex | GO:0034703 | 1.60E-08 | 6.37 | 20 | 47 | 4.99 | 42.6% | 1.5% | Ca-alpha1T,cac,CG42260,CG4587,Cngl,eag,GluRIA,Hk,iav,Ir25a,Ir76b,NaCP60E,para,Sh,Shab,Shaw,trpl,Trpm,unc80,wtrw | Ca[2+]-channel protein alpha[[1]] subunit T,cacophony,Cyclic nucleotide-gated ion channel-like,ether a go-go,Glutamate receptor IA,Hyperkinetic,inactive,Ionotropic receptor 25a,Ionotropic receptor 76b,Na channel protein 60E,paralytic,Shaker,Shaker cognate b,Shaker cognate w,Transient receptor potential cation channel, subfamily M,transient receptor potential-like,uncharacterized protein,uncoordinated 80,water witch |

| | | | | | | | | | | | | | |
|---|---|---|---|---|---|---|---|---|---|---|---|---|---|
| Over-expressed 48 AEL | 1301 | Cellular component | cell periphery | GO:0071944 | 3.90E-07 | 1.61 | 172 | 1148 | 121.77 | 15.0% | 13.2% | 5-HT1A,Ance-4,Atet,bdl,beat-Ic,Ca-alpha1T,cac,CadN2,CCKLR-17D1,CG10483,CG10702,CG10804,CG13404,CG15270,CG1688,CG3078,CG31814,CG32683,CG32812,CG3285,CG33282,CG33310,CG42541,CG44153,CG44325,CG4587,Dh44-R1,dia,DIP-beta,DIP-eta,Dop2R,dos,dpr18,dpr2,dpr20,dpr4,Drl-2,drongo,Dscam4,Dyb,Eaat2 | 5-hydroxytryptamine (serotonin) receptor 1A,ABC transporter expressed in trachea,Ance-4,beaten path Ic,borderless,Ca[2+]-channel protein alpha[[1]] subunit T,cacophony,Cadherin-N2,Cholecystokinin-like receptor at 17D1,daughter of sevenless,defective proboscis extension response 18,defective proboscis extension response 2,defective proboscis extension response 20,defective proboscis extension response 4,Derailed 2,diaphanous,Diuretic hormone 44 receptor 1,Dopamine 2-like receptor,Down |
| Over-expressed 48 AEL | 1301 | Cellular component | neuron projection membrane | GO:0032589 | 1.00E-05 | 5.84 | 13 | 32 | 3.39 | 40.6% | 1.0% | CG32206,CG34402,dpr11,dpr12,dpr18,dpr2,dpr20,dpr4,dpr5,dpr6,Or82a,Rdl,Shaw | defective proboscis extension response 11,defective proboscis extension response 12,defective proboscis extension response 18,defective proboscis extension response 2,defective proboscis extension response 20,defective proboscis extension response 4,defective proboscis extension response 5,defective proboscis extension response 6,Odorant receptor 82a,Resistant to dieldrin,Shaker cognate w,uncharacterized protein |
| Over-expressed 48 AEL | 1301 | Cellular component | ciliary plasm | GO:0097014 | 1.20E-05 | 8.52 | 10 | 20 | 2.12 | 50.0% | 0.8% | CG17387,CG3339,CG5142,CG9492,Dhc62B,Dhc93AB,Dhc98D,mlt,Oseg2,rempA | Dynein heavy chain at 62B,Dynein heavy chain at 89D,Dynein heavy chain at 93AB,mulet,Outer segment 2,reduced mechanoreceptor potential A,uncharacterized protein |
| Over-expressed 48 AEL | 1301 | Cellular component | plasma membrane receptor complex | GO:0098802 | 1.30E-05 | 4.75 | 15 | 42 | 4.46 | 35.7% | 1.2% | CG32206,CG34402,GABA-B-R3,GluRIA,InR,Ir25a,Ir76b,ItgaPS4,mGluR,nAChRalpha2,nAChRalpha3,nAChRalpha5,nAChRalpha6,nAChRalpha7,Nmdar2 | Glutamate receptor IA,Insulin-like receptor,Integrin alphaPS4 subunit,Ionotropic receptor 25a,Ionotropic receptor 76b,metabotropic GABA-B receptor subtype 3,metabotropic Glutamate Receptor,nicotinic Acetylcholine Receptor alpha2,nicotinic Acetylcholine Receptor alpha3,nicotinic Acetylcholine Receptor alpha5,nicotinic Acetylcholine Receptor alpha6,nicotinic Acetylcholine Receptor alpha7,NMDA receptor 2,uncharacterized protein |

| | | | | | | | | | | | | | |
|---|---|---|---|---|---|---|---|---|---|---|---|---|---|
| Over-expressed 48 AEL | 1301 | Cellular component | phagophore assembly site | GO:0000407 | 1.80E-05 | 9.57 | 9 | 17 | 1.8 | 52.9% | 0.7% | Atg1,Atg101,Atg13,Atg17,Atg18a,Atg18b,Atg6,CG5445,Pi3K59F | Autophagy-related 1,Autophagy-related 101,Autophagy-related 13,Autophagy-related 17,Autophagy-related 18a,Autophagy-related 18b,Autophagy-related 6,Phosphotidylinositol 3 kinase 59F,uncharacterized protein |
| Over-expressed 48 AEL | 1301 | Cellular component | rhabdomere | GO:0016028 | 1.00E-04 | 5.2 | 11 | 29 | 3.08 | 37.9% | 0.8% | Arr1,Arr2,Gbeta76C,inaC,ninaC,PIP5K59B,rdgB,Rh5,Rh6,rtp,trpl | Arrestin 1,Arrestin 2,G protein beta-subunit 76C,inactivation no afterpotential C,neither inactivation nor afterpotential C,Phosphatidylinositol 4 phosphate 5-kinase at 59B,retinal degeneration B,retinophilin,Rhodopsin 5,Rhodopsin 6,transient receptor potential-like |
| Over-expressed 48 AEL | 1301 | Cellular component | Atg1/ULK1 kinase complex | GO:1990316 | 0.00013 | Inf | 4 | 4 | 0.42 | 100.0% | 0.3% | Atg1,Atg101,Atg13,Atg17 | Autophagy-related 1,Autophagy-related 101,Autophagy-related 13,Autophagy-related 17 |
| Over-expressed 48 AEL | 1301 | Cellular component | dendrite | GO:0030425 | 0.00016 | 2.73 | 22 | 91 | 9.65 | 24.2% | 1.7% | 5-HT1A,Cbp53E,Dop1R1,drl,fry,futsch,Gr33a,Gr39a,Gr94a,hdc,Ir25a,Ir76b,nAChRalpha7,nudE,Oamb,Or82a,plum,Rdl,Shaw,Timp,trpl,unc-104 | 5-hydroxytryptamine (serotonin) receptor 1A,Calbindin 53E,derailed,Dopamine 1-like receptor 1,furry,futsch,Gustatory receptor 33a,Gustatory receptor 39a,Gustatory receptor 94a,headcase,Ionotropic receptor 25a,Ionotropic receptor 76b,nicotinic Acetylcholine Receptor alpha7,nudE,Octopamine receptor in mushroom bodies,Odorant receptor 82a,plum,Resistant to dieldrin,Shaker cognate w,Tissue inhibitor of metalloproteases,transient receptor potential-like,uncoordinated-104 |
| Over-expressed 48 AEL | 1301 | Cellular component | extracellular region | GO:0005576 | 0.00017 | 1.5 | 114 | 783 | 83.05 | 14.6% | 8.8% | Actbeta,Adgf-D,ALiX,amn,Ance-4,AttA,AttB,AttC,AttD,beat-Ia,Burs,CCHa1,CecA1,CecA2,CG10031,CG10433,CG10730,CG10918,CG11131,CG11852,CG12896,CG13837,CG14457,CG1552,CG16995,CG17843,CG18067,CG18135,CG2650,CG32206,CG33985,CG34402,CG4982,CG5267,CG5867,CG7142,CG8093,CG9449,CheB42c,cmpy,Cpr100A,Cpr49Af,Cpr | Activin-beta,Adenosine deaminase-related growth factor D,ALG-2 interacting protein X,amnesiac,Ance-4,Attacin-A,Attacin-B,Attacin-C,Attacin-D,beaten path Ia,Bursicon,CCHamide-1,Cecropin A1,Cecropin A2,Chemosensory protein B 42c,crimpy,Cuticular protein 100A,Cuticular protein 49Af,Cuticular protein 67B,Cuticular protein 76Bb,Cuticular protein 97Eb,dally-like,defective proboscis extension response 16,Diptericin B,Drosocin,Drosomycin,Ecdysone-inducible gene L2,elevated during infection,Follistatin,Gly |

| | | | | | | | | | | | | | |
|---|---|---|---|---|---|---|---|---|---|---|---|---|---|
| Over-expressed 48 AEL | 1301 | Cellular component | plasma membrane bounded cell projection part | GO:0120038 | 0.00018 | 2.36 | 28 | 134 | 13.76 | 20.9% | 2.2% | 5-HT1A,Alk,ana1,Arr2,Atg8a,BBS4,Burs,Cbp53E,Cc2d2a,CCKLR-17D1,Cep135,cep290,CG1126,CG15666,CG15923,CG17387,CG32206,CG3339,CG34402,CG5142,CG9492,CLIP-190,cmpy,Dhc62B,Dhc93AB,Dhc98D,Dop1R1,dpr11,dpr12,dpr18,dpr2,dpr20,dpr4,dpr5,dpr6,drl,dysc,erm,fry,futsch,gogo,Gr33a,Gr39a,Gr94a,hdc,inaC,Ir25a,Ir76b,kek | 5-hydroxytryptamine (serotonin) receptor 1A,Anaplastic lymphoma kinase,anastral spindle 1,Arrestin 2,Autophagy-related 8a,Bardet-Biedl syndrome 4,Bursicon,Calbindin 53E,Centrosomal protein 135kDa,cep290,Cholecystokinin-like receptor at 17D1,Coiled-coil and C2 domain containing 2A,crimpy,Cytoplasmic linker protein 190,defective proboscis extension response 11,defective proboscis extension response 12,defective proboscis extension response 18,defective proboscis extension response 2,defective proboscis |
| Over-expressed 48 AEL | 1301 | Cellular component | axon | GO:0030424 | 0.00019 | 2.1 | 36 | 184 | 19.52 | 19.6% | 2.8% | Alk,Atg8a,bdl,Burs,Cbp53E,CCKLR-17D1,CG44153,CLIP-190,cmpy,Dab,drl,Dscam2,Dscam3,Dscam4,erm,fry,futsch,gogo,Gr33a,Gr39a,Gr94a,hdc,ImpL2,InR,kek6,nudE,plum,Ptp99A,Rab3-GEF,rad,Rdl,Shaw,Syt7,Trim9,unc-104,unc-13 | Anaplastic lymphoma kinase,Autophagy-related 8a,borderless,Bursicon,Calbindin 53E,Cholecystokinin-like receptor at 17D1,crimpy,Cytoplasmic linker protein 190,derailed,Disabled,Down syndrome cell adhesion molecule 2,Down syndrome cell adhesion molecule 3,Down syndrome cell adhesion molecule 4,earmuff,Ecdysone-inducible gene L2,furry,futsch,golden goal,Gustatory receptor 33a,Gustatory receptor 39a,Gustatory receptor 94a,headcase,Insulin-like receptor,kek6,nudE,plum,Protein tyrosine |
| Over-expressed 48 AEL | 1301 | Cellular component | somatodendritic compartment | GO:0036477 | 0.00019 | 2.2 | 32 | 157 | 16.65 | 20.4% | 2.5% | 5-HT1A,Cbp53E,CCKLR-17D1,Dop1R1,drl,erm,fry,futsch,Gr33a,Gr39a,Gr94a,hdc,hig,Ir25a,Ir76b,nAChRalpha7,nudE,Oamb,Or82a,PICK1,plum,Rab18,Rab3-GEF,Rab39,RabX4,rad,Rdl,Shaw,SK,Timp,trpl,unc-104 | 5-hydroxytryptamine (serotonin) receptor 1A,Calbindin 53E,Cholecystokinin-like receptor at 17D1,derailed,Dopamine 1-like receptor 1,earmuff,furry,futsch,Gustatory receptor 33a,Gustatory receptor 39a,Gustatory receptor 94a,headcase,hikaru genki,Ionotropic receptor 25a,Ionotropic receptor 76b,nicotinic Acetylcholine Receptor alpha7,nudE,Octopamine receptor in mushroom bodies,Odorant receptor 82a,plum,Protein interacting with C kinase 1,Rab18,Rab3 GDP-GTP exchange factor,Rab39,RabX4,rad |

| | | | | | | | | | | | | | |
|---|---|---|---|---|---|---|---|---|---|---|---|---|---|
| Over-expressed 48 AEL | 1301 | Cellular component | extracellular space | GO:0005615 | 0.00025 | 1.68 | 67 | 415 | 44.02 | 16.1% | 5.1% | Actbeta,Adgf-D,amn,Ance-4,AttA,AttB,AttC,AttD,Burs,CCHa1,CecA1,CecA2,CG10031,CG10433,CG10730,CG10918,CG11131,CG11852,CG12896,CG14457,CG1552,CG16995,CG17843,CG18067,CG18135,CG2650,CG32206,CG34402,CG4982,CG5267,CG5867,CG6421,CG7142,CG8093,CG9449,CheB42c,DptB,Drs,edin,Gpb5,Ilp3,ImpL2,IMPPP,lectin-46Ca,lectin- | Activin-beta,Adenosine deaminase-related growth factor D,amnesiac,Ance-4,Attacin-A,Attacin-B,Attacin-C,Attacin-D,Bursicon,CCHamide-1,Cecropin A1,Cecropin A2,Chemosensory protein B 42c,Diptericin B,Drosomycin,Ecdysone-inducible gene L2,elevated during infection,Glycoprotein hormone beta 5,Immune induced molecule prepropeptide,Insulin-like peptide 3,lectin-46Ca,lectin-46Cb,listericin,Neural Lazarillo,obstructor-G,obstructor-H,Odorant receptor 82a,Peptidoglycan recognition protein |
| Over-expressed 48 AEL | 1301 | Cellular component | neuron projection | GO:0043005 | 0.00051 | 3.49 | 13 | 46 | 4.71 | 28.3% | 1.0% | 5-HT1A,Alk,Atg8a,bdl,Burs,Cbp53E,CCKLR-17D1,CG10702,CG32206,CG34402,CG44153,ChAT,CLIP-190,cmpy,Dab,Dop1R1,dpr11,dpr12,dpr18,dpr2,dpr20,dpr4,dpr5,dpr6,drl,Dscam2,Dscam3,Dscam4,dysc,erm,fry,futsch,GluClalpha,gogo,Gr33a,Gr39a,Gr94a,Grd,hdc,ImpL2,InR,Ir25a,Ir76b,kek6,Kif3C,nAChRalpha2,nAChRalpha3,nAChRalpha5,nAChRalph | 5-hydroxytryptamine (serotonin) receptor 1A,Anaplastic lymphoma kinase,Autophagy-related 8a,borderless,Bursicon,Calbindin 53E,Cholecystokinin-like receptor at 17D1,Choline acetyltransferase,crimpy,Cytoplasmic linker protein 190,defective proboscis extension response 11,defective proboscis extension response 12,defective proboscis extension response 18,defective proboscis extension response 2,defective proboscis extension response 20,defective proboscis extension response 4,defective |
| Over-expressed 48 AEL | 1301 | Cellular component | axoneme part | GO:0044447 | 0.00074 | 8.48 | 6 | 12 | 1.27 | 50.0% | 0.5% | CG3339,CG5142,CG9492,Dhc62B,Dhc93AB,Dhc98D | Dynein heavy chain at 62B,Dynein heavy chain at 89D,Dynein heavy chain at 93AB,uncharacterized protein |
| Over-expressed 48 AEL | 1301 | Cellular component | ciliary part | GO:0044441 | 9.00E-04 | 4.55 | 9 | 26 | 2.73 | 34.6% | 0.7% | ana1,BBS4,Cc2d2a,Cep135,cep290,CG1126,CG15666,CG15923,CG17387,CG3339,CG5142,CG9492,Dhc62B,Dhc93AB,Dhc98D,dysc,mlt,Oseg2,rempA | anastral spindle 1,Bardet-Biedl syndrome 4,Centrosomal protein 135kDa,cep290,Coiled-coil and C2 domain containing 2A,Dynein heavy chain at 62B,Dynein heavy chain at 89D,Dynein heavy chain at 93AB,dyschronic,mulet,Outer segment 2,reduced mechanoreceptor potential A,uncharacterized protein |

| | | | | | | | | | | | | | |
|---|---|---|---|---|---|---|---|---|---|---|---|---|---|
| Under-expressed 48 AEL | 1726 | Biological process | proteolysis | GO:0006508 | 1.60E-20 | 2.39 | 201 | 703 | 108.96 | 28.6% | 11.6% | allen,Ance,Ance-2,CalpC,CG10602,CG10764,CG11911,CG11912,CG12256,CG1304,CG13744,CG15253,CG15254,CG15255,CG15485,CG1632,CG17134,CG17337,CG17571,CG17633,CG18477,CG2915,CG30025,CG30031,CG30371,CG30382,CG31267,CG31269,CG31780,CG31954,CG32376,CG32379,CG33459,CG3355,CG4017,CG42264,CG43124,CG43 | allen,Ance-2,Angiotensin converting enzyme,Calpain C,Derlin-1,gammaTrypsin,Heat shock 70-kDa protein cognate 3,Jonah 25Bi,Jonah 25Bii,Jonah 25Biii,Jonah 44E,kelch like family member 10,lambdaTry,lethal (2) k05911,Nardilysin,Nedd8,Notopleural,pomp,prenyl protease type I,Proteasome alpha4 subunit,Proteasome alpha5 subunit,Proteasome alpha6 subunit,Proteasome alpha7 subunit,Proteasome beta4 subunit,Regulatory particle non-ATPase |
| Under-expressed 48 AEL | 1726 | Biological process | chitin-based cuticle development | GO:0040003 | 1.90E-20 | 4.96 | 73 | 157 | 24.33 | 46.5% | 4.2% | amd,Ccp84Aa,Ccp84Ab,Ccp84Ac,Ccp84Ad,Ccp84Af,CG34461,CG34462,Cht5,Cht9,ck,Cpr12A,Cpr23B,Cpr30F,Cpr35B,Cpr47Eb,Cpr47Ec,Cpr47Ed,Cpr49Ac,Cpr49Ad,Cpr49Ag,Cpr49Ah,Cpr51A,Cpr5C,Cpr60D,Cpr64Ab,Cpr64Ad,Cpr65Ea,Cpr65Eb,Cpr65Ec,Cpr66Cb,Cpr66D,Cpr67Fa2,Cpr67Fb,Cpr78Cb,CrebA,dpy,dsx-c73A,exp,gammaCOP,Gld,knk,Lcp2,Lcp4,Lcp65Ab1,Lcp6 | alpha methyl dopa-resistant,Ccp84Aa,Ccp84Ab,Ccp84Ac,Ccp84Ad,Ccp84Af,Chitinase 5,Chitinase 9,Coat Protein (coatomer) gamma,crinkled,Cuticular protein 12A,Cuticular protein 23B,Cuticular protein 30F,Cuticular protein 35B,Cuticular protein 47Eb,Cuticular protein 47Ec,Cuticular protein 47Ed,Cuticular protein 49Ac,Cuticular protein 49Ad,Cuticular protein 49Ag,Cuticular protein 49Ah,Cuticular protein 51A,Cuticular protein 5C,Cuticular protein 60D,Cuticular protein 64Ab,Cuticular protein 64Ad,Cuticular protein 65Ea,Cuticular protein 65Eb,Cuticular protein 65Ec,Cuticular |
| Under-expressed 48 AEL | 1726 | Biological process | DNA replication | GO:0006260 | 1.50E-11 | 4.2 | 45 | 105 | 16.27 | 42.9% | 2.6% | Caf1-180,CDC45L,Cdc6,Cdk1,CG10336,CG11788,CG13690,CG2990,CG3430,Ctf4,CycE,DNA-ligI,DNApol-alpha180,DNApol-alpha50,DNApol-alpha60,DNApol-alpha73,DNApol-epsilon255,DNApol-epsilon58,dpa,dup,Fen1,geminin,hd,Mcm10,Mcm2,Mcm3,Mcm5,Mcm6,Mcm7,mus308,Orc1,O | CDC45L,Cdc6,Chromatin assembly factor 1, p180 subunit,Chromosome transmission fidelity 4,Cyclin E,Cyclin-dependent kinase 1,disc proliferation abnormal,DNA ligase I,DNA polymerase alpha 180kD,DNA polymerase alpha 50kD,DNA polymerase alpha 60kD,DNA polymerase alpha 73kD,DNA polymerase epsilon 255kD subunit,DNA polymerase epsilon 58kD subunit,double parked,Flap endonuclease 1,geminin,humpty dumpty,Minichromosome maintenance 10,Minichromosome |

| | | | | | | | | | | | | | |
|---|---|---|---|---|---|---|---|---|---|---|---|---|---|
| Under-expressed 48 AEL | 1726 | Biological process | DNA replication initiation | GO:0006270 | 3.70E-11 | 19.88 | 18 | 23 | 3.56 | 78.3% | 1.0% | CDC45L,Cdc6,Cdk1,DNApol-alpha180,DNApol-alpha73,dpa,Mcm10,Mcm2,Mcm3,Mcm5,Mcm6,Mcm7,Orc1,Orc2,Orc3,Orc5,Orc6,Rif1 | CDC45L,Cdc6,Cyclin-dependent kinase 1,disc proliferation abnormal,DNA polymerase alpha 180kD,DNA polymerase alpha 73kD,Minichromosome maintenance 10,Minichromosome maintenance 2,Minichromosome maintenance 3,Minichromosome maintenance 5,Minichromosome maintenance 6,Minichromosome maintenance 7,Origin recognition complex subunit 1,Origin recognition complex subunit 2,Origin recognition complex subunit 3,Origin recognition complex |
| Under-expressed 48 AEL | 1726 | Biological process | establishment of protein localization to endoplasmic reticulum | GO:0072599 | 1.90E-10 | 18.76 | 17 | 22 | 3.41 | 77.3% | 1.0% | CG5885,Gtp-bp,Sec61alpha,Sec61beta,Sec63,Sgt,Spase12,Spase22-23,Spase25,Srp14,Srp19,Srp54k,Srp68,Srp72,Srp9,TRAM,Trp1 | GTP-binding protein,Sec61 alpha subunit,Sec61 beta subunit,Secretory 63,Signal recognition particle protein 14,Signal recognition particle protein 19,Signal recognition particle protein 54k,Signal recognition particle protein 68,Signal recognition particle protein 72,Signal recognition particle protein 9,small glutamine-rich tetratricopeptide containing protein,Spase 12-subunit,Spase 22/23-subunit,Spase 25-subunit,TRAM,Translocation protein 1,uncharacterized protein |
| Under-expressed 48 AEL | 1726 | Biological process | proteasomal ubiquitin-independent protein catabolic process | GO:0010499 | 2.20E-09 | 35.78 | 13 | 15 | 2.32 | 86.7% | 0.8% | Prosalpha2,Prosalpha3,Prosalpha4,Prosalpha5,Prosalpha6,Prosalpha7,Prosbeta1,Prosbeta2,Prosbeta3,Prosbeta4,Prosbeta5,Prosbeta6,Prosbeta7 | Proteasome alpha2 subunit,Proteasome alpha3 subunit,Proteasome alpha4 subunit,Proteasome alpha5 subunit,Proteasome alpha6 subunit,Proteasome alpha7 subunit,Proteasome beta1 subunit,Proteasome beta2 subunit,Proteasome beta3 subunit,Proteasome beta4 subunit,Proteasome beta5 subunit,Proteasome beta6 subunit,Proteasome beta7 subunit |

| | | | | | | | | | | | | | |
|---|---|---|---|---|---|---|---|---|---|---|---|---|---|
| Under-expressed 48 AEL | 1726 | Biological process | protein folding | GO:0006457 | 3.40E-07 | 3.97 | 27 | 65 | 10.01 | 41.5% | 1.6% | boca,Calr,CCT1,CCT2,CCT3,CCT5,CCT7,CCT8,CG11267,CG16817,CG17266,CG2852,CG3678,CG7872,CNPYb,Cnx99A,Cyp1,Cypl,EMC1,EMC3,ERp60,FK506-bp2,FKBP59,Gp93,HIP,HIP-R,Hsc70-3,Hsp67Ba,Hsp68,Hsp70Ba,l(2)efl,Nrd1,nudC,P58IPK,Pdi,Pfdn5,Pfdn6,sel,Sgt,shv,Torsin,wbl | boca,Calnexin 99A,Calreticulin,Canopy b,Chaperonin containing TCP1 subunit 1,Chaperonin containing TCP1 subunit 2,Chaperonin containing TCP1 subunit 3,Chaperonin containing TCP1 subunit 5,Chaperonin containing TCP1 subunit 7,Chaperonin containing TCP1 subunit 8,Cyclophilin 1,Cyclophilin-like,ER Membrane protein Complex 1,ER Membrane protein Complex 3,ERp60,FK506-binding protein 2,FK506-binding protein FKBP59,Glycoprotein 93,Heat shock 70-kDa protein cognate 3,Heat |
| Under-expressed 48 AEL | 1726 | Biological process | ER to Golgi vesicle-mediated transport | GO:0006888 | 4.20E-07 | 3.78 | 28 | 69 | 10.69 | 40.6% | 1.6% | alphaCOP,Arl1,bai,beta'COP,betaCOP,boca,CG32069,CG4293,CG5484,CG5510,CG7011,CG9175,CHOp24,deltaCOP,eca,epsilonCOP,ergic53,gammaCOP,KdelR,loj,Membrin,p115,p24-1,p24-2,Sec13,Sec22,Sec23,Sec24CD | ADP ribosylation factor-like 1,baiser,boca,CHOp24,Coat Protein (coatomer) alpha,Coat Protein (coatomer) beta,Coat Protein (coatomer) beta',Coat Protein (coatomer) delta,Coat Protein (coatomer) epsilon,Coat Protein (coatomer) gamma,eclair,ergic53,KDEL receptor,logjam,membrin,p115,p24-related-1,p24-related-2,Secretory 13,Secretory 22,Secretory 23,Secretory 24CD,uncharacterized protein |
| Under-expressed 48 AEL | 1726 | Biological process | eggshell chorion gene amplification | GO:0007307 | 5.20E-05 | 6.71 | 11 | 20 | 3.1 | 55.0% | 0.6% | Caf1-55,CycE,dup,geminin,hd,Mcm6,mip40,Orc2,PCNA,pic,RecQ4 | Chromatin assembly factor 1, p55 subunit,Cyclin E,double parked,geminin,humpty dumpty,Minichromosome maintenance 6,Myb-interacting protein 40,Origin recognition complex subunit 2,piccolo,Proliferating cell nuclear antigen,RecQ4 helicase |
| Under-expressed 48 AEL | 1726 | Biological process | body morphogenesis | GO:0010171 | 8.00E-05 | 5.49 | 12 | 24 | 3.72 | 50.0% | 0.7% | Tb,TwdlB,Twdlbeta,TwdlD,TwdlG,TwdlM,TwdlR,TwdlS,TwdlT,TwdlW,TwdlX,TwdlY | Tubby,TweedleB,Tweedlebeta,TweedleD,TweedleG,TweedleM,TweedleR,TweedleS,TweedleT,TweedleW,TweedleX,TweedleY |

| | | | | | | | | | | | | | |
|---|---|---|---|---|---|---|---|---|---|---|---|---|---|
| Under-expressed 48 AEL | 1726 | Biological process | SRP-dependent cotranslational protein targeting to membrane | GO:0006614 | 8.20E-05 | 33 | 6 | 7 | 1.08 | 85.7% | 0.3% | CG5885,Gtp-bp,Sec61alpha,Sec61beta,Sec63,Srp14,Srp19,Srp54k,Srp68,Srp72,Srp9,TRAM | GTP-binding protein,Sec61 alpha subunit,Sec61 beta subunit,Secretory 63,Signal recognition particle protein 14,Signal recognition particle protein 19,Signal recognition particle protein 54k,Signal recognition particle protein 68,Signal recognition particle protein 72,Signal recognition particle protein 9,TRAM,uncharacterized protein |
| Under-expressed 48 AEL | 1726 | Biological process | organonitrogen compound metabolic process | GO:1901564 | 8.20E-05 | 1.27 | 477 | 2693 | 417.41 | 17.7% | 27.6% | AIF,Alg10,alpha-PheRS,ArgRS,Atox1,Btnd,CalpC,CG11911,CG11912,CG12256,CG1304,CG15343,CG1632,CG1885,CG30008,CG30025,CG30031,CG30291,CG30382,CG30496,CG31267,CG31269,CG31780,CG31917,CG31954,CG32276,CG32284,CG32376,CG32379,CG33303,CG33459,CG3355,CG3964,CG4020,CG42264,CG43124,CG43125,CG4678,CG48 | Alpha 3 glucosyltransferase,Antioxidant 1 copper chaperone,Apoptosis inducing factor,Arginyl-tRNA synthetase,Biotinidase,Calpain C,Catalytic subunit 3A of the oligosaccharyltransferase complex,Chitinase 10,Cyclophilin 1,dachsous,Derlin-1,dodo,Heat shock 70-kDa protein cognate 3,Jonah 25Bi,Jonah 25Bii,Jonah 25Biii,kelch like family member 10,lesswright,mitochondrial ribosomal protein L10,mitochondrial ribosomal protein L14,mitochondrial ribosomal protein L16,mitochondrial ribosomal protein |
| Under-expressed 48 AEL | 1726 | Biological process | SRP-dependent cotranslational protein targeting to membrane, translocation | GO:0006616 | 8.30E-05 | 32.85 | 6 | 7 | 1.08 | 85.7% | 0.3% | Sec61alpha,Sec61beta,Srp19,Srp72,Srp9,TRAM | Sec61 alpha subunit,Sec61 beta subunit,Signal recognition particle protein 19,Signal recognition particle protein 72,Signal recognition particle protein 9,TRAM |
| Under-expressed 48 AEL | 1726 | Biological process | positive regulation of RNA polymerase II transcriptional preinitiation complex assembly | GO:0045899 | 8.30E-05 | 32.85 | 6 | 7 | 1.08 | 85.7% | 0.3% | Rpt1,Rpt3,Rpt3R,Rpt4,Rpt5,Rpt6 | Regulatory particle triple-A ATPase 1,Regulatory particle triple-A ATPase 3,Regulatory particle triple-A ATPase 3-related,Regulatory particle triple-A ATPase 4,Regulatory particle triple-A ATPase 5,Regulatory particle triple-A ATPase 6 |

| | | | | | | | | | | | | | |
|---|---|---|---|---|---|---|---|---|---|---|---|---|---|
| Under-expressed 48 AEL | 1726 | Biological process | macromolecule glycosylation | GO:0043413 | 1.00E-04 | 2.62 | 27 | 84 | 13.02 | 32.1% | 1.6% | Alg10,CG10166,CG11149,CG32276,CG33303,CG33774,CG33977,CG34452,CG3792,CG7840,CG8708,CG9669,Dad1,meigo,O-fut1,Ost48,OstDelta,Ostgamma,Pabp2,pgant3,rt,rumi,Stt3A,Stt3B,tw,Ugt,wol | Alpha 3 glucosyltransferase,Catalytic subunit 3A of the oligosaccharyltransferase complex,Catalytic subunit 3B of the oligosaccharyltransferase complex,Defender against apoptotic cell death 1,medial glomeruli,O-fucosyltransferase 1,Oligosaccharide transferase Delta subunit,Oligosaccharide transferase gamma subunit,Oligosaccharyltransferase 48kD subunit,Pabp2,polypeptide GalNAc transferase 3,rotated abdomen,rumi,twisted,UDP-glucose-glycoprotein glucosyltransferase,uncharacterized |
| Under-expressed 48 AEL | 1726 | Biological process | Golgi organization | GO:0007030 | 0.00017 | 2.56 | 26 | 82 | 12.71 | 31.7% | 1.5% | Arl1,bai,CG10738,CG13049,CG13284,CG16817,CG5510,CG7456,CHOp24,eca,ergic53,GPHR,Grasp65,loj,mEFTu1,Ost48,p115,p24-1,p24-2,rumi,Sec23,Sec24CD,stg,Tango2,TER94,wol | ADP ribosylation factor-like 1,baiser,CHOp24,eclair,ergic53,Golgi pH Regulator,Grasp65,logjam,mitochondrial translation elongation factor Tu 1,Oligosaccharyltransferase 48kD subunit,p115,p24-related-1,p24-related-2,rumi,Secretory 23,Secretory 24CD,string,TER94,Transport and Golgi organization 2,uncharacterized protein,wollknaeuel |
| Under-expressed 48 AEL | 1726 | Biological process | glycoprotein biosynthetic process | GO:0009101 | 0.00017 | 2.37 | 30 | 100 | 15.5 | 30.0% | 1.7% | Alg10,beta4GalT7,CG10166,CG11149,CG32276,CG33303,CG33774,CG33977,CG34452,CG3792,CG7840,CG8708,CG9669,Dad1,meigo,O-fut1,Ost48,OstDelta,Ostgamma,Pabp2,Papst2,pgant3,rt,rumi,sll,Stt3A,Stt3B,tw,Ugt,wol | Alpha 3 glucosyltransferase,beta-4-galactosyltransferase 7,Catalytic subunit 3A of the oligosaccharyltransferase complex,Catalytic subunit 3B of the oligosaccharyltransferase complex,Defender against apoptotic cell death 1,medial glomeruli,O-fucosyltransferase 1,Oligosaccharide transferase Delta subunit,Oligosaccharide transferase gamma subunit,Oligosaccharyltransferase 48kD subunit,Pabp2,PAPS transporter 2,polypeptide GalNAc transferase 3,rotated abdomen,rumi,slalom,t |
| Under-expressed 48 AEL | 1726 | Biological process | DNA strand elongation involved in DNA replication | GO:0006271 | 2.00E-04 | 7.05 | 9 | 16 | 2.48 | 56.2% | 0.5% | CG13690,CG2990,DNA-ligI,DNApol-alpha180,DNApol-epsilon255,Fen1,PCNA,Psf1,RfC38 | DNA ligase I,DNA polymerase alpha 180kD,DNA polymerase epsilon 255kD subunit,Flap endonuclease 1,Proliferating cell nuclear antigen,Psf1,Replication factor C 38kD subunit,uncharacterized protein |

| | | | | | | | | | | | | | |
|---|---|---|---|---|---|---|---|---|---|---|---|---|---|
| Under-expressed 48 AEL | 1726 | Biological process | protein targeting | GO:0006605 | 0.00022 | 2.42 | 28 | 92 | 14.26 | 30.4% | 1.6% | boca,CG14270,CG34132,CG5885,CG8860,Gtp-bp,Membrin,Sec61alpha,Sec61beta,Sec61gamma,Sec63,Sgt,Spase12,Spase22-23,Spase25,Srp14,Srp19,Srp54k,Srp68,Srp72,Srp9,Tim8,Tim9a,Tim9b,Tom70,TRAM,Trp1,ttm50 | boca,GTP-binding protein,membrin,Sec61 alpha subunit,Sec61 beta subunit,Sec61 gamma subunit,Secretory 63,Signal recognition particle protein 14,Signal recognition particle protein 19,Signal recognition particle protein 54k,Signal recognition particle protein 68,Signal recognition particle protein 72,Signal recognition particle protein 9,small glutamine-rich tetratricopeptide containing protein,Spase 12-subunit,Spase 22/23-subunit,Spase 25-subunit,Tim8,Tim9a,tiny tim |
| Under-expressed 48 AEL | 1726 | Biological process | proteasome-mediated ubiquitin-dependent protein catabolic process | GO:0043161 | 0.00022 | 1.93 | 46 | 178 | 27.59 | 25.8% | 2.7% | CG30382,Der-1,Fbxl7,Hsc70-3,ida,pic,Plap,Pomp,Prosalpha2,Prosalpha3,Prosalpha4,Prosalpha5,Prosalpha6,Prosalpha7,Prosbeta1,Prosbeta2,Prosbeta3,Prosbeta4,Prosbeta5,Prosbeta6,Prosbeta7,Rpn1,Rpn10,Rpn11,Rpn12,Rpn2,Rpn3,Rpn5,Rpn7,Rpn8,Rpn9,Rpt1,Rpt2,Rpt3,Rpt3R,Rpt4,Rpt5,Rpt6,Sec13,Sem1,Sgt,sip3,Skp2,TER94,Uch-L5,Usp5 | Derlin-1,F-box and leucine-rich repeat protein 7,Heat shock 70-kDa protein cognate 3,imaginal discs arrested,Phospholipase A2 activator protein,piccolo,pomp,Proteasome alpha2 subunit,Proteasome alpha3 subunit,Proteasome alpha4 subunit,Proteasome alpha5 subunit,Proteasome alpha6 subunit,Proteasome alpha7 subunit,Proteasome beta1 subunit,Proteasome beta2 subunit,Proteasome beta3 subunit,Proteasome |
| Under-expressed 48 AEL | 1726 | Biological process | DNA biosynthetic process | GO:0071897 | 0.00031 | 3.38 | 16 | 42 | 6.51 | 38.1% | 0.9% | Caf1-55,CDC45L,CycE,DNA-ligI,DNApol-epsilon58,dup,geminin,hd,Mcm6,mip40,Orc1,Orc2,PCNA,pic,RecQ4,RPA2 | CDC45L,Chromatin assembly factor 1, p55 subunit,Cyclin E,DNA ligase I,DNA polymerase epsilon 58kD subunit,double parked,geminin,humpty dumpty,Minichromosome maintenance 6,Myb-interacting protein 40,Origin recognition complex subunit 1,Origin recognition complex subunit 2,piccolo,Proliferating cell nuclear antigen,RecQ4 helicase,Replication protein A2 |

| | | | | | | | | | | | | | |
|---|---|---|---|---|---|---|---|---|---|---|---|---|---|
| Under-expressed 48 AEL | 1726 | Biological process | protein N-linked glycosylation | GO:0006487 | 0.00037 | 3.97 | 13 | 31 | 4.8 | 41.9% | 0.8% | Alg10,CG33303,CG33774,CG7840,CG9669,Dad1,meigo,Ost48,OstDelta,Ostgamma,Stt3B,Ugt,wol | Alpha 3 glucosyltransferase,Catalytic subunit 3B of the oligosaccharyltransferase complex,Defender against apoptotic cell death 1,medial glomeruli,Oligosaccharide transferase Delta subunit,Oligosaccharide transferase gamma subunit,Oligosaccharyltransferase 48kD subunit,UDP-glucose-glycoprotein glucosyltransferase,uncharacterized protein,wollknaeuel |
| Under-expressed 48 AEL | 1726 | Biological process | wax biosynthetic process | GO:0010025 | 0.00039 | 9.59 | 7 | 11 | 1.7 | 63.6% | 0.4% | CG30427,CG4020,CG5065,CG8303,CG8306,FarO,wat | Fatty acyl-CoA reductase in oenocytes,uncharacterized protein,waterproof |
| Under-expressed 48 AEL | 1726 | Biological process | posttranslational protein targeting to endoplasmic reticulum membrane | GO:0006620 | 0.00046 | 27.36 | 5 | 6 | 0.93 | 83.3% | 0.3% | Sec61alpha,Sec61beta,Sec63,Sgt,Trp1 | Sec61 alpha subunit,Sec61 beta subunit,Secretory 63,small glutamine-rich tetratricopeptide containing protein,Translocation protein 1 |
| Under-expressed 48 AEL | 1726 | Biological process | chaperone-mediated protein folding | GO:0061077 | 0.00057 | 3.3 | 15 | 40 | 6.2 | 37.5% | 0.9% | CG11267,CNPYb,FK506-bp2,FKBP59,HIP,HIP-R,Hsc70-3,Hsp67Ba,Hsp68,Hsp70Ba,l(2)efl,Nrd1,sel,Sgt,Torsin | Canopy b,FK506-binding protein 2,FK506-binding protein FKBP59,Heat shock 70-kDa protein cognate 3,Heat shock gene 67Ba,Heat shock protein 68,Heat-shock-protein-70Ba,Hsc/Hsp70-interacting protein,Hsc/Hsp70-interacting protein related,lethal (2) essential for life,Nardilysin,seele,small glutamine-rich tetratricopeptide containing protein,torsin,uncharacterized protein |
| Under-expressed 48 AEL | 1726 | Biological process | regulation of post-mating oviposition | GO:0048042 | 0.00075 | 10.95 | 6 | 9 | 1.39 | 66.7% | 0.3% | bai,CHOp24,eca,loj,p24-1,p24-2 | baiser,CHOp24,eclair,logjam,p24-related-1,p24-related-2 |
| Under-expressed 48 AEL | 1726 | Biological process | acyl-CoA metabolic process | GO:0006637 | 0.00077 | 3.57 | 13 | 33 | 5.11 | 39.4% | 0.8% | ATPCL,CG10268,CG30427,CG33671,CG4020,CG5065,CG8239,CG8303,CG8306,FarO,scu,Sucb,wat | ATP citrate lyase,Fatty acyl-CoA reductase in oenocytes,scully,sucb,uncharacterized protein,waterproof |
| Under-expressed 48 AEL | 1726 | Biological process | long-chain fatty-acyl-CoA metabolic process | GO:0035336 | 0.00082 | 7.67 | 7 | 12 | 1.86 | 58.3% | 0.4% | CG30427,CG4020,CG5065,CG8303,CG8306,FarO,wat | Fatty acyl-CoA reductase in oenocytes,uncharacterized protein,waterproof |

| | | | | | | | | | | | | | |
|---|---|---|---|---|---|---|---|---|---|---|---|---|---|
| Under-expressed 48 AEL | 1726 | Biological process | carbohydrate derivative metabolic process | GO:1901135 | 0.00097 | 1.43 | 112 | 550 | 85.25 | 20.4% | 6.5% | Adgf-C,Adk3,AIF,Alg10,ATPCL,awd,beta4GalT7,CG10166,CG10268,CG10474,CG10738,CG11149,CG11164,CG12009,CG13675,CG13676,CG13806,CG14291,CG14304,CG14880,CG17145,CG30427,CG32276,CG32284,CG33303,CG33671,CG33774,CG33977,CG34282,CG3788,CG3792,CG4020,CG4302,CG5065,CG5167,CG5613,CG7840,CG8239,CG8303,C | abnormal wing discs,Adenosine deaminase-related growth factor C,Adenylate kinase 3,Alpha 3 glucosyltransferase,Apoptosis inducing factor,ATP citrate lyase,beta-4-galactosyltransferase 7,Catalytic subunit 3A of the oligosaccharyltransferase complex,Catalytic subunit 3B of the oligosaccharyltransferase complex,Chitinase 10,Chitinase 5,Chitinase 9,Defender against apoptotic cell death 1,deoxyribonucleoside kinase,Deoxyuridine triphosphatase,expansion,GDP-4-keto-6-deoxy- |
| Under-expressed 48 AEL | 1726 | Molecular function | peptidase activity, acting on L-amino acid peptides | GO:0070011 | 5.70E-21 | 3.71 | 102 | 261 | 40.57 | 39.1% | 5.9% | Ance,Ance-2,Ance-5,CalpC,CG10472,CG10602,CG10764,CG11911,CG11912,CG12256,CG1304,CG13744,CG14820,CG15253,CG15254,CG15255,CG15485,CG16998,CG17134,CG17337,CG17571,CG17633,CG2915,CG30025,CG30031,CG30371,CG30382,CG31267,CG31269,CG31780,CG31954,CG32374,CG32376,CG32379,CG3344,CG33459,CG33465,CG3 | Ance-2,Ance-5,Angiotensin converting enzyme,Calpain C,gammaTrypsin,Jonah 25Bi,Jonah 25Bii,Jonah 25Biii,Jonah 44E,Jonah 65Ai,Jonah 65Aii,Jonah 65Aiii,Jonah 65Aiv,Jonah 66Ci,Jonah 66Cii,lambdaTry,lethal (2) k05911,Nardilysin,Notopleural,prenyl protease type I,Proteasome alpha3 subunit,Proteasome alpha4 subunit,Proteasome alpha5 subunit,Proteasome alpha6 subunit,Proteasome alpha7 subunit,Proteasome beta4 |
| Under-expressed 48 AEL | 1726 | Molecular function | serine-type endopeptidase activity | GO:0004252 | 4.10E-18 | 4.52 | 70 | 156 | 24.67 | 44.9% | 4.1% | CG10472,CG10764,CG11836,CG11842,CG11843,CG11911,CG11912,CG12256,CG1304,CG13744,CG16749,CG16998,CG17475,CG17477,CG17571,CG30025,CG30031,CG30371,CG31266,CG31267,CG31269,CG31780,CG31954,CG32374,CG32376,CG33459,CG33465,CG3355,CG34437,CG34458,CG4053,CG43124,CG43125,CG4386,CG4613,CG4914,CG5255,CG | deltaTrypsin,easter,epsilonTrypsin,gammaTrypsin,Jonah 25Bi,Jonah 25Bii,Jonah 25Biii,Jonah 44E,Jonah 65Ai,Jonah 65Aii,Jonah 65Aiii,Jonah 65Aiv,Jonah 66Ci,Jonah 66Cii,Jonah 74E,Jonah 99Cii,Jonah 99Ciii,Jonah 99Fii,lambdaTry,lethal (2) k05911,Notopleural,Serine protease 6,Stubble,thetaTrypsin,uncharacterized protein,yippee interacting protein 7 |

| | | | | | | | | | | | | | |
|---|---|---|---|---|---|---|---|---|---|---|---|---|---|
| Under-expressed 48 AEL | 1726 | Molecular function | serine hydrolase activity | GO:0017171 | 5.80E-17 | 3.99 | 74 | 177 | 28 | 41.8% | 4.3% | CG10472,CG10764,CG11836,CG11842,CG11843,CG11911,CG11912,CG12256,CG1304,CG13744,CG16749,CG16998,CG17475,CG17477,CG17571,CG30025,CG30031,CG30371,CG31266,CG31267,CG31269,CG31780,CG31954,CG32374,CG32376,CG3344,CG33459,CG33465,CG3355,CG34437,CG34458,CG3734,CG4053,CG43124,CG43125,CG4386,CG4613,CG491 | deltaTrypsin,easter,epsilonTrypsin,gammaTrypsin,Jonah 25Bi,Jonah 25Bii,Jonah 25Biii,Jonah 44E,Jonah 65Ai,Jonah 65Aii,Jonah 65Aiii,Jonah 65Aiv,Jonah 66Ci,Jonah 66Cii,Jonah 74E,Jonah 99Cii,Jonah 99Ciii,Jonah 99Fii,lambdaTry,lethal (2) k05911,Notopleural,Serine protease 6,Stubble,thetaTrypsin,twisted bristles roughened eye,uncharacterized protein,yippee interacting protein 7 |
| Under-expressed 48 AEL | 1726 | Molecular function | structural constituent of chitin-based larval cuticle | GO:0008010 | 4.50E-15 | 7.05 | 40 | 71 | 11.23 | 56.3% | 2.3% | Ccp84Aa,Ccp84Ab,Ccp84Ac,Ccp84Ad,Ccp84Af,CG34461,CG34462,Cpr12A,Cpr23B,Cpr30F,Cpr35B,Cpr47Eb,Cpr47Ec,Cpr47Ed,Cpr49Ac,Cpr49Ad,Cpr49Ag,Cpr49Ah,Cpr51A,Cpr5C,Cpr60D,Cpr64Ab,Cpr64Ad,Cpr65Ea,Cpr65Eb,Cpr65Ec,Cpr66Cb,Cpr66D,Cpr67Fa2,Cpr67Fb,Cpr78Cb,Lcp2,Lcp4,Lcp65Ab1,Lcp65Ad,Lcp65Af,Lcp65Ag1,Pcp,TwdlD,TwdlH | Ccp84Aa,Ccp84Ab,Ccp84Ac,Ccp84Ad,Ccp84Af,Cuticular protein 12A,Cuticular protein 23B,Cuticular protein 30F,Cuticular protein 35B,Cuticular protein 47Eb,Cuticular protein 47Ec,Cuticular protein 47Ed,Cuticular protein 49Ac,Cuticular protein 49Ad,Cuticular protein 49Ag,Cuticular protein 49Ah,Cuticular protein 51A,Cuticular protein 5C,Cuticular protein 60D,Cuticular protein 64Ab,Cuticular protein 64Ad,Cuticular protein 65Ea,Cuticular protein 65Eb,Cuticular protein 65Ec,Cuticular protein 66Cb,Cuticular protein 66D,Cuticular protein 67Fa2,Cuticular protein 67Fb,Cuticular protein |
| Under-expressed 48 AEL | 1726 | Molecular function | threonine-type endopeptidase activity | GO:0004298 | 5.10E-10 | 37.65 | 14 | 16 | 2.53 | 87.5% | 0.8% | CG30382,Prosalpha2,Prosalpha3,Prosalpha4,Prosalpha5,Prosalpha6,Prosalpha7,Prosbeta1,Prosbeta2,Prosbeta3,Prosbeta4,Prosbeta5,Prosbeta6,Prosbeta7 | Proteasome alpha2 subunit,Proteasome alpha3 subunit,Proteasome alpha4 subunit,Proteasome alpha5 subunit,Proteasome alpha6 subunit,Proteasome alpha7 subunit,Proteasome beta1 subunit,Proteasome beta2 subunit,Proteasome beta3 subunit,Proteasome beta4 subunit,Proteasome beta5 subunit,Proteasome beta6 subunit,Proteasome beta7 subunit,uncharacterize |

| | | | | | | | | | | | | | |
|---|---|---|---|---|---|---|---|---|---|---|---|---|---|
| Under-expressed 48 AEL | 1726 | Molecular function | structural constituent of chitin-based cuticle | GO:0005214 | 5.00E-08 | 9.83 | 16 | 25 | 3.87 | 64.0% | 0.9% | Ccp84Aa,Ccp84Ab,Ccp84Ac, Ccp84Ad,Ccp84Af,CG34461, CG34462,Cpr12A,Cpr23B,Cpr30F,Cpr35B,Cpr47Eb,Cpr47Ec,Cpr47Ed,Cpr49Ac,Cpr49Ad,Cpr49Ag,Cpr49Ah,Cpr51A, Cpr5C,Cpr60D, Cpr64Ab,Cpr64Ad,Cpr65Ea, Cpr65Eb,Cpr65Ec,Cpr66Cb,Cpr66D,Cpr67Fa2,Cpr67Fb,Cpr78Cb,dsx-c73A,Lcp2,Lcp4,Lcp65Ab1,Lcp65Ad,Lcp65Af,Lcp65Ag1,m,Pcp,Tb,TwdlB,Twdlbeta,Tw | Ccp84Aa,Ccp84Ab,Ccp84Ac,Ccp84Ad,Ccp84Af,Cuticular protein 12A,Cuticular protein 23B,Cuticular protein 30F,Cuticular protein 35B,Cuticular protein 47Eb,Cuticular protein 47Ec,Cuticular protein 47Ed,Cuticular protein 49Ac,Cuticular protein 49Ad,Cuticular protein 49Ag,Cuticular protein 49Ah,Cuticular protein 51A,Cuticular protein 5C,Cuticular protein 60D,Cuticular protein 64Ab,Cuticular protein 64Ad,Cuticular protein 65Ea,Cuticular protein 65Eb,Cuticular protein 65Ec,Cuticular protein 66Cb,Cuticular protein 66D,Cuticular protein 67Fa2,Cuticular protein 67Fb,Cuticular protein |
| Under-expressed 48 AEL | 1726 | Molecular function | DNA replication origin binding | GO:0003688 | 9.00E-08 | 53.63 | 10 | 11 | 1.74 | 90.9% | 0.6% | CDC45L,Cdc6, DNApol-alpha180,Mcm10,Mcm5,Orc1,Orc2,Orc3,Orc5,Orc6 | CDC45L,Cdc6,DNA polymerase alpha 180kD,Minichromosome maintenance 10,Minichromosome maintenance 5,Origin recognition complex subunit 1,Origin recognition complex subunit 2,Origin recognition complex subunit 3,Origin recognition complex subunit 5,Origin recognition complex subunit 6 |
| Under-expressed 48 AEL | 1726 | Molecular function | structural molecule activity | GO:0005198 | 1.70E-07 | 1.88 | 106 | 418 | 66.11 | 25.4% | 6.1% | Act42A,alphaCOP,alphaTub84B,alphaTub84D,betaCOP, betaTub56D,bonsai,Ccp84Aa,Ccp84Ab,Ccp84Ac,Ccp84Ad,Ccp84Af,CG10602,CG31268,CG8771,Clc,Cpr12A,Cpr23B,Cpr30F,Cpr35B,Cpr47Eb, Cpr47Ec,Cpr47Ed,Cpr49Ac, Cpr49Ad,Cpr49Ag,Cpr49Ah, Cpr51A,Cpr5C, Cpr60D,Cpr64Ab,Cpr64Ad,Cpr65Ea,Cpr65Eb,Cpr65Ec,Cpr66Cb,Cpr66D, Cpr67Fa2,Cpr67Fb,Cpr78Cb | Actin 42A,alpha-Tubulin at 84B,alpha-Tubulin at 84D,beta-Tubulin at 56D,bonsai,Ccp84Aa,Ccp84Ab,Ccp84Ac,Ccp84Ad,Ccp84Af,Clathrin light chain,Coat Protein (coatomer) alpha,Coat Protein (coatomer) beta,Coat Protein (coatomer) epsilon,Coat Protein (coatomer) gamma,Cuticular protein 12A,Cuticular protein 23B,Cuticular protein 30F,Cuticular protein 35B,Cuticular protein 47Eb,Cuticular protein 47Ec,Cuticular protein 47Ed,Cuticular protein 49Ac,Cuticular protein 49Ad,Cuticular protein 49Ag,Cuticular protein 49Ah,Cuticular |
| Under-expressed 48 AEL | 1726 | Molecular function | dolichyl-diphosphooligosaccharide-protein glycotransferase activity | GO:0004579 | 2.40E-06 | Inf | 7 | 7 | 1.11 | 100.0% | 0.4% | CG33303,CG33774,Dad1,Ost48,OstDelta, Ostgamma,Stt3B | Catalytic subunit 3B of the oligosaccharyltransferase complex,Defender against apoptotic cell death 1,Oligosaccharide transferase Delta subunit,Oligosaccharide transferase gamma subunit,Oligosaccharyltransferase 48kD subunit,uncharacterized protein |

| | | | | | | | | | | | | | |
|---|---|---|---|---|---|---|---|---|---|---|---|---|---|
| Under-expressed 48 AEL | 1726 | Molecular function | exopeptidase activity | GO:0008238 | 8.30E-06 | 3.09 | 28 | 77 | 12.18 | 36.4% | 1.6% | Ance,Ance-2,Ance-5,CG14820,CG17337,CG17633,CG2915,CG32379,CG32473,CG3344,CG3502,CG3734,CG4017,CG42264,CG4408,CG4678,CG4829,CG5188,CG5355,CG7025,CG7860,CG8560,CG8562,CG8945,CG9953,DppIII,grsm,SP1029 | Ance-2,Ance-5,Angiotensin converting enzyme,Dipeptidyl aminopeptidase III,granny smith,SP1029,uncharacterized protein |
| Under-expressed 48 AEL | 1726 | Molecular function | alkaline phosphatase activity | GO:0004035 | 8.40E-06 | 16.07 | 9 | 12 | 1.9 | 75.0% | 0.5% | Alp1,Alp4,CG10592,CG16771,CG3264,CG3292,CG5150,CG8105,phu | Alkaline phosphatase 1,Alkaline phosphatase 4,phurba tashi,uncharacterized protein |
| Under-expressed 48 AEL | 1726 | Molecular function | 7S RNA binding | GO:0008312 | 1.60E-05 | Inf | 6 | 6 | 0.95 | 100.0% | 0.3% | Srp14,Srp19,Srp54k,Srp68,Srp72,Srp9 | Signal recognition particle protein 14,Signal recognition particle protein 19,Signal recognition particle protein 54k,Signal recognition particle protein 68,Signal recognition particle protein 72,Signal recognition particle protein 9 |
| Under-expressed 48 AEL | 1726 | Molecular function | proteasome-activating ATPase activity | GO:0036402 | 1.70E-05 | 37.45 | 7 | 8 | 1.27 | 87.5% | 0.4% | Rpt1,Rpt2,Rpt3,Rpt3R,Rpt4,Rpt5,Rpt6 | Regulatory particle triple-A ATPase 1,Regulatory particle triple-A ATPase 2,Regulatory particle triple-A ATPase 3,Regulatory particle triple-A ATPase 3-related,Regulatory particle triple-A ATPase 4,Regulatory particle triple-A ATPase 5,Regulatory particle triple-A ATPase 6 |
| Under-expressed 48 AEL | 1726 | Molecular function | metallocarboxypeptidase activity | GO:0004181 | 1.70E-05 | 7.15 | 12 | 21 | 3.32 | 57.1% | 0.7% | CG14820,CG17633,CG2915,CG32379,CG4017,CG42264,CG4408,CG4678,CG7025,CG8560,CG8562,CG8945 | uncharacterized protein |

| | | | | | | | | | | | | | |
|---|---|---|---|---|---|---|---|---|---|---|---|---|---|
| Under-expressed 48 AEL | 1726 | Molecular function | metallopeptidase activity | GO:0008237 | 2.10E-05 | 2.44 | 37 | 119 | 18.82 | 31.1% | 2.1% | Ance,Ance-2,CG10602,CG14527,CG14820,CG15253,CG15254,CG15255,CG15485,CG17337,CG17633,CG2915,CG32379,CG32473,CG3502,CG4017,CG42264,CG4408,CG4678,CG5131,CG5188,CG5715,CG6696,CG6763,CG7025,CG7631,CG8550,CG8560,CG8562,CG8945,grsm,Nep2,Nrd1,Rpn11,SP1029,ste24a,stl | Ance-2,Angiotensin converting enzyme,granny smith,Nardilysin,Neprilysin 2,prenyl protease type I,Regulatory particle non-ATPase 11,SP1029,stall,uncharacterized protein |
| Under-expressed 48 AEL | 1726 | Molecular function | hydrolase activity | GO:0016787 | 3.00E-05 | 1.39 | 246 | 1292 | 196.93 | 19.0% | 14.3% | alpha-PheRS,Btnd,CalpC,CDC45L,CG11164,CG11911,CG11912,CG12256,CG1304,CG13690,CG15743,CG17134,CG17633,CG18258,CG18641,CG1986,CG2990,CG30016,CG30025,CG30031,CG30382,CG31267,CG31269,CG31780,CG31954,CG32376,CG32379,CG32750,CG32751,CG33459,CG3355,CG4017,CG42237,CG42264,CG43124,CG43125,CG4678,C | Biotinidase,Calpain C,CDC45L,Chitinase 10,dachs,Deoxyuridine triphosphatase,Derlin-1,eukaryotic translation release factor 3,Fatty acid synthase 3,Heat shock 70-kDa protein cognate 3,Helicase at 25E,Jonah 25Bi,Jonah 25Bii,Jonah 25Biii,Juvenile hormone epoxide hydrolase 1,Lysosomal alpha-mannosidase V,Maltase B1,Minichromosome maintenance 3,Minichromosome maintenance 6,mitochondrial translation elongation factor G 1,mulder,mus81,Myosin 10A,Nardilysin,Peptido |
| Under-expressed 48 AEL | 1726 | Molecular function | transferase activity, transferring hexosyl groups | GO:0016758 | 4.60E-05 | 2.27 | 39 | 132 | 20.88 | 29.5% | 2.3% | Alg10,beta4GalT7,CG10166,CG10178,CG11149,CG11851,CG30438,CG33303,CG33774,CG33977,CG34452,CG4302,CG5724,CG6475,CG8708,Dad1,GalNAc-T1,GalNAc-T2,O-fut1,Ost48,OstDelta,Ostgamma,pgant3,pgant5,pgant6,PIG-B,PIG-C,rt,rumi,Stt3A,Stt3B,tw,Ugt,Ugt35b,Ugt37b1,Ugt37c1,Ugt86Dd,Ugt86De,wol | Alpha 3 glucosyltransferase,beta-4-galactosyltransferase 7,Catalytic subunit 3A of the oligosaccharyltransferase complex,Catalytic subunit 3B of the oligosaccharyltransferase complex,Defender against apoptotic cell death 1,GalNAc-T1,O-fucosyltransferase 1,Oligosaccharide transferase Delta subunit,Oligosaccharide transferase gamma subunit,Oligosaccharyltransferase 48kD subunit,Phosphatidylinositol glycan anchor biosynthesis class B,Phosphatidylinositol glycan anchor biosynthesis class |

| | | | | | | | | | | | | | |
|---|---|---|---|---|---|---|---|---|---|---|---|---|---|
| Under-expressed 48 AEL | 1726 | Molecular function | oxidoreductase activity, acting on the CH-OH group of donors, NAD or NADP as acceptor | GO:0016616 | 6.40E-05 | 3.36 | 20 | 52 | 8.22 | 38.5% | 1.2% | Adh,Adhr,CG10131,CG10638,CG1236,CG12766,CG18814,CG2767,CG6287,CG7322,Fdh,Gmer,Gpdh,Idh,Mfe2,Mtpalpha,Pgd,scu,Sptr,Zw | Adh-related,Alcohol dehydrogenase,Formaldehyde dehydrogenase,GDP-4-keto-6-deoxy-D-mannose 3,5-epimerase/4-reductase,Glycerol 3 phosphate dehydrogenase,Isocitrate dehydrogenase,Mitochondrial trifunctional protein alpha subunit,peroxisomal Multifunctional enzyme type 2,Phosphogluconate dehydrogenase,scully,Sepiapterin reductase,uncharacterized protein,Zwischenferment |
| Under-expressed 48 AEL | 1726 | Molecular function | oxidoreductase activity | GO:0016491 | 8.00E-05 | 1.58 | 106 | 475 | 75.13 | 22.3% | 6.1% | Acox57D-d,Adh,Adhr,AIF,AOX2,Arc42,CG10131,CG10211,CG10638,CG10672,CG11162,CG12262,CG1236,CG12766,CG14997,CG15343,CG15531,CG18547,CG1998,CG2767,CG30427,CG31075,CG31371,CG31559,CG3270,CG4020,CG42331,CG45065,CG4860,CG5065,CG5167,CG5321,CG5948,CG6287,CG7322,CG7460,CG7484,CG7560,CG7840,CG8303,CG830 | acyl-Coenzyme A oxidase at 57D distal,Adh-related,Alcohol dehydrogenase,Aldehyde oxidase 2,Apoptosis inducing factor,Arc42,Coproporphyrinogen oxidase,Curly Su,Cyp28d2,Cyp301a1,Cyp304a1,Cyp305a1,Cyp310a1,Cyp318a1,Cyp49a1,Cyp4ad1,Cyp4d14,Cyp6a18,Cyp6a21,Cyp6g1,Cyp6t1,Cyp9f2,Cytochrome P450-4d1,Cytochrome P450-4e1,Cytochrome P450-4e2,Cytochrome P450-6a2,Cytochrome P450-6a8,Dihydropteridine reductase,Ecdysone oxidase,Fatty acid synthase 3,Formaldehyde dehydrogenase,GDP-4- |
| Under-expressed 48 AEL | 1726 | Molecular function | unfolded protein binding | GO:0051082 | 0.00012 | 2.64 | 26 | 79 | 12.49 | 32.9% | 1.5% | Calr,CCT1,CCT2,CCT3,CCT5,CCT7,CCT8,CG11267,CG17266,CG2852,Cnx99A,Cyp1,Gp93,Hsc70-3,Hsp67Ba,Hsp68,Hsp70Ba,l(2)efl,nudC,P58IPK,Pfdn1,Pfdn5,Pfdn6,shv,sip3,Ugt | Calnexin 99A,Calreticulin,Chaperonin containing TCP1 subunit 1,Chaperonin containing TCP1 subunit 2,Chaperonin containing TCP1 subunit 3,Chaperonin containing TCP1 subunit 5,Chaperonin containing TCP1 subunit 7,Chaperonin containing TCP1 subunit 8,Cyclophilin 1,Glycoprotein 93,Heat shock 70-kDa protein cognate 3,Heat shock gene 67Ba,Heat shock protein 68,Heat-shock-protein-70Ba,lethal (2) essential for life,nudC,P58IPK,Prefoldin 1,Prefoldin 5,Prefoldin 6,septin interacting protein 3,shriveled,UDP- |
| Under-expressed 48 AEL | 1726 | Molecular function | fatty-acyl-CoA reductase (alcohol-forming) activity | GO:0080019 | 0.00045 | 9.36 | 7 | 11 | 1.74 | 63.6% | 0.4% | CG30427,CG4020,CG5065,CG8303,CG8306,FarO,wat | Fatty acyl-CoA reductase in oenocytes,uncharacterized protein,waterproof |

| | | | | | | | | | | | | | |
|---|---|---|---|---|---|---|---|---|---|---|---|---|---|
| Under-expressed 48 AEL | 1726 | Molecular function | catalytic activity, acting on a protein | GO:0140096 | 0.00083 | 1.31 | 213 | 1114 | 176.2 | 19.1% | 12.3% | Ance,Ance-2,Art8,aurB,CalpC,Cdk1,Cdk4,CG10602,CG11911,CG11912,CG12256,CG1304,CG13744,CG15253,CG15254,CG15255,CG15485,CG17134,CG17266,CG17337,CG17571,CG17633,CG2915,CG30025,CG30031,CG30371,CG30382,CG31267,CG31269,CG31780,CG31954,CG32376,CG32379,CG33459,CG3355,CG3964,CG4017,CG42264,CG43124,CG4312 | Ance-2,Angiotensin converting enzyme,Arginine methyltransferase 8,aurora B,Calpain C,Cyclin E,Cyclin-dependent kinase 1,Cyclin-dependent kinase 4,Cyclophilin 1,Derlin-1,dodo,ERp60,GalNAc-T1,gammaTrypsin,Jonah 25Bi,Jonah 25Bii,Jonah 25Biii,Jonah 44E,lambdaTry,lesswright,lethal (2) k05911,Nardilysin,Notopleural,O-fucosyltransferase 1,Phosphatidylinositol glycan anchor biosynthesis class U,polypeptide GalNAc transferase 3,polypeptide GalNAc |
| Under-expressed 48 AEL | 1726 | Cellular component | endoplasmic reticulum | GO:0005783 | 2.20E-25 | 3.44 | 139 | 382 | 58.56 | 36.4% | 8.1% | Alg10,CaBP1,CG10166,CG10470,CG11162,CG11999,CG13284,CG1371,CG15168,CG17593,CG1998,CG2023,CG2046,CG2254,CG2781,CG2852,CG2918,CG30008,CG32069,CG32276,CG32576,CG33129,CG33303,CG33774,CG33977,CG4293,CG4302,CG44252,CG4880,CG4972,CG6230,CG6746,CG7011,CG7149,CG7484,CG7840,CG7872,CG8331,CG9175,CG966 | Alpha 3 glucosyltransferase,calcium-binding protein 1,Canopy b,CHOp24,Defender against apoptotic cell death 1,Derlin-1,ER Membrane protein Complex 1,ER Membrane protein Complex 3,ERp60,FK506-binding protein 14,Golgi pH Regulator,Grasp65,Heat shock 70-kDa protein cognate 3,heixuedian,jagunal,KDEL receptor,Lamin B receptor,lethal (1) G0320,lethal (2) neighbor of tid,logjam,membrin,Microsomal triacylglycerol transfer protein,O-fucosyltransferase 1,Oligosaccharide |
| Under-expressed 48 AEL | 1726 | Cellular component | extracellular matrix | GO:0031012 | 3.60E-20 | 4.86 | 73 | 160 | 24.53 | 45.6% | 4.2% | Ccp84Aa,Ccp84Ab,Ccp84Ac,Ccp84Ad,Ccp84Af,CG17211,CG30203,CG31268,CG34461,CG34462,CG43333,Cpr12A,Cpr23B,Cpr30F,Cpr35B,Cpr47Eb,Cpr47Ec,Cpr47Ed,Cpr49Ac,Cpr49Ad,Cpr49Ag,Cpr49Ah,Cpr51A,Cpr5C,Cpr60D,Cpr64Ab,Cpr64Ad,Cpr65Ea,Cpr65Eb,Cpr65Ec,Cpr66Cb,Cpr66D,Cpr67Fa2,Cpr67Fb,Cpr78Cb,dpy,LanA,LanB1,Lcp2,Lcp4,Lcp65Ab1,Lcp65Ad,Lc | Ccp84Aa,Ccp84Ab,Ccp84Ac,Ccp84Ad,Ccp84Af,Cuticular protein 12A,Cuticular protein 23B,Cuticular protein 30F,Cuticular protein 35B,Cuticular protein 47Eb,Cuticular protein 47Ec,Cuticular protein 47Ed,Cuticular protein 49Ac,Cuticular protein 49Ad,Cuticular protein 49Ag,Cuticular protein 49Ah,Cuticular protein 51A,Cuticular protein 5C,Cuticular protein 60D,Cuticular protein 64Ab,Cuticular protein 64Ad,Cuticular protein 65Ea,Cuticular protein 65Eb,Cuticular protein 65Ec,Cuticular protein 66Cb,Cuticular protein 66D,Cuticular protein 67Fa2,Cuticular protein 67Fb,Cuticular protein |

| | | | | | | | | | | | | | |
|---|---|---|---|---|---|---|---|---|---|---|---|---|---|
| Under-expressed 48 AEL | 1726 | Cellular component | extracellular region | GO:0005576 | 7.20E-18 | 2.2 | 208 | 783 | 120.03 | 26.6% | 12.1% | Ance,Ance-2,Ance-5,CG12256,CG13748,CG13806,CG15225,CG17211,CG17337,CG17633,CG17974,CG2852,CG2915,CG2918,CG30197,CG30203,CG31268,CG31997,CG32284,CG32379,CG33459,CG3355,CG4017,CG42264,CG42326,CG43333,CG43366,CG4386,CG4678,CG4847,CG5177,CG5397,CG6337,CG6347,CG6426,CG7025,CG8160,CG8213,CG846 | Ance-2,Ance-5,Angiotensin converting enzyme,Chitinase 10,Chitinase 9,crinkled,Cuticular protein 12A,Cuticular protein 23B,Cuticular protein 30F,Cuticular protein 35B,Cuticular protein 47Eb,Cuticular protein 47Ec,Cuticular protein 47Ed,Cuticular protein 49Ac,Cuticular protein 49Ad,Cuticular protein 49Ag,Cuticular protein 49Ah,Cuticular protein 51A,Cuticular protein 5C,Cuticular protein 60D,dumpy,FK506-binding protein 14,four-jointed,GIP-like,Glial Lazarillo,Heat shock 70-kDa protein cognate 3,LanB1,Larval cuticle |
| Under-expressed 48 AEL | 1726 | Cellular component | peptidase complex | GO:1905368 | 8.40E-13 | 6.2 | 36 | 69 | 10.55 | 52.2% | 2.1% | CG30382,not,Pomp,Prosalpha2,Prosalpha3,Prosalpha4,Prosalpha5,Prosalpha6,Prosalpha7,Prosbeta1,Prosbeta2,Prosbeta3,Prosbeta4,Prosbeta5,Prosbeta6,Prosbeta7,Rpn1,Rpn10,Rpn11,Rpn12,Rpn2,Rpn3,Rpn5,Rpn7,Rpn8,Rpn9,Rpt1,Rpt2,Rpt3,Rpt3R,Rpt4,Rpt5,Rpt6,Sem1,Spase12,Spase22-23,Spase25,TER94,twr,Uch-L5 | non-stop,pomp,Proteasome alpha2 subunit,Proteasome alpha3 subunit,Proteasome alpha4 subunit,Proteasome alpha5 subunit,Proteasome alpha6 subunit,Proteasome alpha7 subunit,Proteasome beta1 subunit,Proteasome beta2 subunit,Proteasome beta3 subunit,Proteasome beta4 subunit,Proteasome beta5 subunit,Proteasome beta6 subunit,Proteasome |
| Under-expressed 48 AEL | 1726 | Cellular component | proteasome accessory complex | GO:0022624 | 2.10E-11 | 17.75 | 19 | 25 | 3.83 | 76.0% | 1.1% | Rpn1,Rpn10,Rpn11,Rpn12,Rpn2,Rpn3,Rpn5,Rpn7,Rpn8,Rpn9,Rpt1,Rpt2,Rpt3,Rpt3R,Rpt4,Rpt5,Rpt6,Sem1,Uch-L5 | Regulatory particle non-ATPase 1,Regulatory particle non-ATPase 10,Regulatory particle non-ATPase 11,Regulatory particle non-ATPase 12,Regulatory particle non-ATPase 2,Regulatory particle non-ATPase 3,Regulatory particle non-ATPase 5,Regulatory particle non-ATPase 7,Regulatory particle non-ATPase 8,Regulatory particle non-ATPase 9,Regulatory particle triple-A ATPase 1,Regulatory particle triple-A ATPase 2,Regulatory particle triple-A ATPase 3,Regulatory particle |

| | | | | | | | | | | | | | |
|---|---|---|---|---|---|---|---|---|---|---|---|---|---|
| Under-expressed 48 AEL | 1726 | Cellular component | organelle subcompartment | GO:0031984 | 1.10E-09 | 2.3 | 88 | 308 | 47.22 | 28.6% | 5.1% | Alg10,alt,Arf79F,ArfGAP1,Arl1,beta'COP,CG10166,CG10470,CG11162,CG11999,CG1371,CG14511,CG15168,CG15531,CG16904,CG1998,CG2781,CG30008,CG32069,CG33052,CG33303,CG33774,CG33977,CG3678,CG4293,CG4972,CG5484,CG5510,CG6230,CG6660,CG7011,CG7149,CG7872,CG8708,CG9175,CG9662,CG9669,CG9747,Csat,Dad1,Der-1,EMC1,EMC3 | ADP ribosylation factor at 79F,ADP ribosylation factor-like 1,ADP-ribosylation factor GTPase activating protein 1,Alpha 3 glucosyltransferase,aluminum tubes,Catalytic subunit 3B of the oligosaccharyltransferase complex,Coat Protein (coatomer) beta',Coat Protein (coatomer) gamma,csat,Defender against apoptotic cell death 1,Derlin-1,ER Membrane protein Complex 1,ER Membrane protein Complex 3,ergic53,GalNAc-T1,Golgi complex-localized glycoprotein 1,Golgi pH Regulator,Grasp65,GTP- |
| Under-expressed 48 AEL | 1726 | Cellular component | COPII-coated ER to Golgi transport vesicle | GO:0030134 | 2.30E-09 | 11.2 | 18 | 27 | 4.14 | 66.7% | 1.0% | bai,CG32069,CG4293,CG5484,CG5510,CG7011,CHOp24,eca,ergic53,loj,Membrin,p115,p24-1,p24-2,Sec13,Sec22,Sec23,Sec24CD | baiser,CHOp24,eclair,ergic53,logjam,membrin,p115,p24-related-1,p24-related-2,Secretory 13,Secretory 22,Secretory 23,Secretory 24CD,uncharacterized protein |
| Under-expressed 48 AEL | 1726 | Cellular component | proteasome complex | GO:0000502 | 5.90E-09 | 12.97 | 16 | 23 | 3.48 | 69.6% | 0.9% | CG30382,Pomp,Prosalpha2,Prosalpha3,Prosalpha4,Prosalpha5,Prosalpha6,Prosalpha7,Prosbeta1,Prosbeta2,Prosbeta3,Prosbeta4,Prosbeta5,Prosbeta6,Prosbeta7,Rpn1,Rpn10,Rpn11,Rpn12,Rpn2,Rpn3,Rpn5,Rpn7,Rpn8,Rpn9,Rpt1,Rpt2,Rpt3,Rpt3R,Rpt4,Rpt5,Rpt6,Sem1,TER94,Uch-L5 | pomp,Proteasome alpha2 subunit,Proteasome alpha3 subunit,Proteasome alpha4 subunit,Proteasome alpha5 subunit,Proteasome alpha6 subunit,Proteasome alpha7 subunit,Proteasome beta1 subunit,Proteasome beta2 subunit,Proteasome beta3 subunit,Proteasome beta4 subunit,Proteasome beta5 subunit,Proteasome beta6 subunit,Proteasome beta7 |

| | | | | | | | | | | | | | |
|---|---|---|---|---|---|---|---|---|---|---|---|---|---|
| Under-expressed 48 AEL | 1726 | Cellular component | extracellular space | GO:0005615 | 1.00E-08 | 2.01 | 107 | 415 | 63.62 | 25.8% | 6.2% | 26-29-p,Adgf-C,Alp4,Ance,Ance-2,Ance-5,apolpp,betaGlu,Calr,Ccp84Ad,CG10264,CG11836,CG11854,CG12256,CG13748,CG14820,CG15225,CG15497,CG17271,CG17633,CG17919,CG17974,CG2016,CG2852,CG2915,CG2918,CG31266,CG31268,CG31997,CG32379,CG33459,CG3355,CG34034,CG3868,CG4017,CG42264,CG42326,CG43333,CG43366,CG4386,C | 26-29kD-proteinase,Adenosine deaminase-related growth factor C,Alkaline phosphatase 4,Ance-2,Ance-5,Angiotensin converting enzyme,apolipophorin,beta glucuronidase,Calreticulin,Ccp84Ad,Chitinase 5,Chitinase 9,Cuticular protein 51A,Cuticular protein 67Fb,easter,Esterase 6,FK506-binding protein 14,four-jointed,Glial Lazarillo,Glycoprotein 93,Heat shock 70-kDa protein cognate 3,Insulin-like peptide 2,lethal (2) k05911,Lysozyme D,Lysozyme P,Lysozyme |
| Under-expressed 48 AEL | 1726 | Cellular component | oligosaccharyltransferase complex | GO:0008250 | 4.60E-08 | Inf | 9 | 9 | 1.38 | 100.0% | 0.5% | CG33303,CG33774,CG9662,CG9669,Dad1,Ost48,OstDelta,Ostgamma,Stt3B | Catalytic subunit 3B of the oligosaccharyltransferase complex,Defender against apoptotic cell death 1,Oligosaccharide transferase Delta subunit,Oligosaccharide transferase gamma subunit,Oligosaccharyltransferase 48kD subunit,uncharacterized protein |
| Under-expressed 48 AEL | 1726 | Cellular component | endoplasmic reticulum membrane | GO:0005789 | 8.50E-08 | 2.69 | 50 | 157 | 23.77 | 31.8% | 2.9% | Alg10,alt,CG10166,CG10470,CG11162,CG11999,CG1371,CG14511,CG15168,CG15531,CG16904,CG1998,CG2781,CG30008,CG32069,CG33303,CG33774,CG33977,CG3678,CG4293,CG4972,CG5484,CG5510,CG6230,CG6660,CG7011,CG7149,CG7872,CG9175,CG9662,CG9669,CG9747,Dad1,Der-1,EMC1,EMC3,ergic53,Gtp-bp,jagn,l(1)G0320,l(2)not,L | Alpha 3 glucosyltransferase,aluminum tubes,Catalytic subunit 3B of the oligosaccharyltransferase complex,Defender against apoptotic cell death 1,Derlin-1,ER Membrane protein Complex 1,ER Membrane protein Complex 3,ergic53,GTP-binding protein,jagunal,Lamin B receptor,lethal (1) G0320,lethal (2) neighbor of tid,medial glomeruli,membrin,Oligosaccharide transferase Delta subunit,Oligosaccharide transferase gamma subunit,Oligosaccharyltransferase 48kD subunit,PAPS transporter |
| Under-expressed 48 AEL | 1726 | Cellular component | endoplasmic reticulum-Golgi intermediate compartment | GO:0005793 | 4.70E-07 | 13.38 | 12 | 17 | 2.61 | 70.6% | 0.7% | alphaCOP,bai,CG5484,CG5510,CHOp24,eca,ergic53,gammaCOP,loj,p24-1,p24-2,Sec22 | baiser,CHOp24,Coat Protein (coatomer) alpha,Coat Protein (coatomer) gamma,eclair,ergic53,logjam,p24-related-1,p24-related-2,Secretory 22,uncharacterized protein |

| | | | | | | | | | | | | |
|---|---|---|---|---|---|---|---|---|---|---|---|---|
| Under-expressed 48 AEL | 1726 | Cellular component | proteasome regulatory particle, base subcomplex | GO:0008540 | 1.30E-06 | 18.55 | 10 | 13 | 1.99 | 76.9% | 0.6% | Rpn1,Rpn10,Rpn2,Rpt1,Rpt2,Rpt3,Rpt3R,Rpt4,Rpt5,Rpt6 | Regulatory particle non-ATPase 1,Regulatory particle non-ATPase 10,Regulatory particle non-ATPase 2,Regulatory particle triple-A ATPase 1,Regulatory particle triple-A ATPase 2,Regulatory particle triple-A ATPase 3,Regulatory particle triple-A ATPase 3-related,Regulatory particle triple-A ATPase 4,Regulatory particle triple-A ATPase 5,Regulatory particle triple-A ATPase 6 |
| Under-expressed 48 AEL | 1726 | Cellular component | MCM complex | GO:0042555 | 2.00E-06 | Inf | 7 | 7 | 1.07 | 100.0% | 0.4% | CG3430,dpa,Mcm2,Mcm3,Mcm5,Mcm6,Mcm7 | disc proliferation abnormal,Minichromosome maintenance 2,Minichromosome maintenance 3,Minichromosome maintenance 5,Minichromosome maintenance 6,Minichromosome maintenance 7,uncharacterized protein |
| Under-expressed 48 AEL | 1726 | Cellular component | integral component of endoplasmic reticulum membrane | GO:0030176 | 2.40E-06 | 3.56 | 26 | 67 | 10.27 | 38.8% | 1.5% | alt,CG10470,CG14511,CG15168,CG16904,CG2781,CG30008,CG3678,CG4293,CG6660,CG7011,CG9175,Dad1,EMC1,EMC3,l(2)not,LBR,meigo,Papst2,PIG-G,PIG-U,Sec63,Spase12,Spp,ste24a,Trp1 | aluminum tubes,Defender against apoptotic cell death 1,ER Membrane protein Complex 1,ER Membrane protein Complex 3,Lamin B receptor,lethal (2) neighbor of tid,medial glomeruli,PAPS transporter 2,Phosphatidylinositol glycan anchor biosynthesis class G,Phosphatidylinositol glycan anchor biosynthesis class U,prenyl protease type I,Secretory 63,Signal peptide peptidase,Spase 12-subunit,Translocation protein 1,uncharacterized protein |
| Under-expressed 48 AEL | 1726 | Cellular component | proteasome regulatory particle, lid subcomplex | GO:0008541 | 1.00E-05 | 22.23 | 8 | 10 | 1.53 | 80.0% | 0.5% | Rpn11,Rpn12,Rpn3,Rpn5,Rpn7,Rpn8,Rpn9,Sem1 | Regulatory particle non-ATPase 11,Regulatory particle non-ATPase 12,Regulatory particle non-ATPase 3,Regulatory particle non-ATPase 5,Regulatory particle non-ATPase 7,Regulatory particle non-ATPase 8,Regulatory particle non-ATPase 9,Suppressor of exocyst mutations 1 |

| | | | | | | | | | | | | | |
|---|---|---|---|---|---|---|---|---|---|---|---|---|---|
| Under-expressed 48 AEL | 1726 | Cellular component | proteasome core complex, beta-subunit complex | GO:0019774 | 1.30E-05 | Inf | 6 | 6 | 0.92 | 100.0% | 0.3% | Prosbeta1,Prosbeta2,Prosbeta3,Prosbeta5,Prosbeta6,Prosbeta7 | Proteasome beta1 subunit,Proteasome beta2 subunit,Proteasome beta3 subunit,Proteasome beta5 subunit,Proteasome beta6 subunit,Proteasome beta7 subunit |
| Under-expressed 48 AEL | 1726 | Cellular component | signal recognition particle, endoplasmic reticulum targeting | GO:0005786 | 1.40E-05 | 38.88 | 7 | 8 | 1.23 | 87.5% | 0.4% | Srp14,Srp19,Srp54k,Srp68,Srp72,Srp9,SsRbeta | Signal recognition particle protein 14,Signal recognition particle protein 19,Signal recognition particle protein 54k,Signal recognition particle protein 68,Signal recognition particle protein 72,Signal recognition particle protein 9,Signal sequence receptor beta |
| Under-expressed 48 AEL | 1726 | Cellular component | proteasome core complex, alpha-subunit complex | GO:0019773 | 1.40E-05 | 38.88 | 7 | 8 | 1.23 | 87.5% | 0.4% | CG30382,Prosalpha2,Prosalpha3,Prosalpha4,Prosalpha5,Prosalpha6,Prosalpha7 | Proteasome alpha2 subunit,Proteasome alpha3 subunit,Proteasome alpha4 subunit,Proteasome alpha5 subunit,Proteasome alpha6 subunit,Proteasome alpha7 subunit,uncharacterized protein |
| Under-expressed 48 AEL | 1726 | Cellular component | U6 snRNP | GO:0005688 | 7.80E-05 | 33.3 | 6 | 7 | 1.07 | 85.7% | 0.3% | CG10418,CG2021,CG6610,CG9344,LSm3,LSm7 | Like Sm 3,Like Sm 7,uncharacterized protein |
| Under-expressed 48 AEL | 1726 | Cellular component | cytosolic proteasome complex | GO:0031597 | 7.80E-05 | 33.3 | 6 | 7 | 1.07 | 85.7% | 0.3% | Rpn1,Rpn2,Rpt3,Rpt3R,Rpt4,Rpt6 | Regulatory particle non-ATPase 1,Regulatory particle non-ATPase 2,Regulatory particle triple-A ATPase 3,Regulatory particle triple-A ATPase 3-related,Regulatory particle triple-A ATPase 4,Regulatory particle triple-A ATPase 6 |
| Under-expressed 48 AEL | 1726 | Cellular component | nuclear nucleosome | GO:0000788 | 8.40E-05 | Inf | 5 | 5 | 0.77 | 100.0% | 0.3% | cid,His2A:CG33808,His2B:CG33872,His2B:CG33874,His2B:CG33876 | centromere identifier,histone H2A,histone H2B |
| Under-expressed 48 AEL | 1726 | Cellular component | Golgi-associated vesicle membrane | GO:0030660 | 9.50E-05 | 4.83 | 13 | 28 | 4.29 | 46.4% | 0.8% | alphaCOP,beta'COP,betaCOP,Clc,deltaCOP,epsilonCOP,gammaCOP,Membrin,p115,Sec13,Sec22,Sec23,Sec24CD | Clathrin light chain,Coat Protein (coatomer) alpha,Coat Protein (coatomer) beta,Coat Protein (coatomer) beta',Coat Protein (coatomer) delta,Coat Protein (coatomer) epsilon,Coat Protein (coatomer) gamma,membrin,p115,Secretory 13,Secretory 22,Secretory 23,Secretory 24CD |

| | | | | | | | | | | | | | |
|---|---|---|---|---|---|---|---|---|---|---|---|---|---|
| Under-expressed 48 AEL | 1726 | Cellular component | protein-DNA complex | GO:0032993 | 0.00013 | 3.98 | 15 | 36 | 5.52 | 41.7% | 0.9% | CDC45L,cid,D1,DNApol-alpha180,DNApol-alpha50,DNApol-alpha60,DNApol-alpha73,His2A:CG33808,His2Av,His2B:CG33872,His2B:CG33874,His2B:CG33876,Orc3,Psf1,RPA2 | CDC45L,centromere identifier,D1 chromosomal protein,DNA polymerase alpha 180kD,DNA polymerase alpha 50kD,DNA polymerase alpha 60kD,DNA polymerase alpha 73kD,histone H2A,Histone H2A variant,histone H2B,Origin recognition complex subunit 3,Psf1,Replication protein A2 |
| Under-expressed 48 AEL | 1726 | Cellular component | chaperonin-containing T-complex | GO:0005832 | 0.00027 | 16.65 | 6 | 8 | 1.23 | 75.0% | 0.3% | CCT1,CCT2,CCT3,CCT5,CCT7,CCT8 | Chaperonin containing TCP1 subunit 1,Chaperonin containing TCP1 subunit 2,Chaperonin containing TCP1 subunit 3,Chaperonin containing TCP1 subunit 5,Chaperonin containing TCP1 subunit 7,Chaperonin containing TCP1 subunit 8 |
| Under-expressed 48 AEL | 1726 | Cellular component | nuclear origin of replication recognition complex | GO:0005664 | 0.00044 | 27.73 | 5 | 6 | 0.92 | 83.3% | 0.3% | Orc1,Orc2,Orc3,Orc5,Orc6 | Origin recognition complex subunit 1,Origin recognition complex subunit 2,Origin recognition complex subunit 3,Origin recognition complex subunit 5,Origin recognition complex subunit 6 |
| Under-expressed 48 AEL | 1726 | Cellular component | rough endoplasmic reticulum membrane | GO:0030867 | 0.00044 | 27.73 | 5 | 6 | 0.92 | 83.3% | 0.3% | Der-1,Gtp-bp,Sec61alpha,Sec61beta,Sec63 | Derlin-1,GTP-binding protein,Sec61 alpha subunit,Sec61 beta subunit,Secretory 63 |
| Under-expressed 48 AEL | 1726 | Cellular component | nuclear replication fork | GO:0043596 | 0.00048 | 5.05 | 10 | 21 | 3.22 | 47.6% | 0.6% | CDC45L,CG10336,Ctf4,DNApol-alpha180,DNApol-alpha50,DNApol-alpha60,DNApol-alpha73,Mcm10,RPA2,timeout | CDC45L,Chromosome transmission fidelity 4,DNA polymerase alpha 180kD,DNA polymerase alpha 50kD,DNA polymerase alpha 60kD,DNA polymerase alpha 73kD,Minichromosome maintenance 10,Replication protein A2,timeout,uncharacterized protein |
| Under-expressed 48 AEL | 1726 | Cellular component | alpha DNA polymerase:primase complex | GO:0005658 | 0.00055 | Inf | 4 | 4 | 0.61 | 100.0% | 0.2% | DNApol-alpha180,DNApol-alpha50,DNApol-alpha60,DNApol-alpha73 | DNA polymerase alpha 180kD,DNA polymerase alpha 50kD,DNA polymerase alpha 60kD,DNA polymerase alpha 73kD |
| Under-expressed 48 AEL | 1726 | Cellular component | signal peptidase complex | GO:0005787 | 0.00055 | Inf | 4 | 4 | 0.61 | 100.0% | 0.2% | Spase12,Spase22-23,Spase25,twr | Spase 12-subunit,Spase 22/23-subunit,Spase 25-subunit,twisted bristles roughened eye |

| | | | | | | | | | | | | | |
|---|---|---|---|---|---|---|---|---|---|---|---|---|---|
| Under-expressed 48 AEL | 1726 | Cellular component | small nuclear ribonucleoprotein complex | GO:0030532 | 7.00E-04 | 2.72 | 19 | 58 | 8.89 | 32.8% | 1.1% | CG10418,CG2021,CG3542,CG4849,CG6610,CG9344,hoip,LSm3,LSm7,Phf5a,Prp8,Sf3b6,SmB,SmD1,SmD2,SmE,SmF,SNRPG,U2A | hoi-polloi,Like Sm 3,Like Sm 7,PHD finger protein 5a,pre-mRNA processing factor 8,Small nuclear ribonucleoprotein G,Small ribonucleoprotein particle protein SmB,Small ribonucleoprotein particle protein SmD1,Small ribonucleoprotein particle protein SmD2,Small ribonucleoprotein particle protein SmE,Small ribonucleoprotein particle protein SmF,Splicing factor 3b subunit 6,U2A,uncharacterized protein |
| Under-expressed 48 AEL | 1726 | Cellular component | COPI vesicle coat | GO:0030126 | 0.00071 | 11.1 | 6 | 9 | 1.38 | 66.7% | 0.3% | alphaCOP,beta'COP,betaCOP,deltaCOP,epsilonCOP,gammaCOP | Coat Protein (coatomer) alpha,Coat Protein (coatomer) beta,Coat Protein (coatomer) beta',Coat Protein (coatomer) delta,Coat Protein (coatomer) epsilon,Coat Protein (coatomer) gamma |
| Under-expressed 48 AEL | 1726 | Cellular component | COPI-coated vesicle | GO:0030137 | 0.00071 | 11.1 | 6 | 9 | 1.38 | 66.7% | 0.3% | alphaCOP,beta'COP,betaCOP,deltaCOP,epsilonCOP,gammaCOP | Coat Protein (coatomer) alpha,Coat Protein (coatomer) beta,Coat Protein (coatomer) beta',Coat Protein (coatomer) delta,Coat Protein (coatomer) epsilon,Coat Protein (coatomer) gamma |
| Under-expressed 48 AEL | 1726 | Cellular component | nuclear pore outer ring | GO:0031080 | 0.00071 | 11.1 | 6 | 9 | 1.38 | 66.7% | 0.3% | Nup107,Nup133,Nup37,Nup43,Nup75,Sec13 | Nucleoporin 107kD,Nucleoporin 133kD,Nucleoporin 37kD,Nucleoporin 43kD,Nucleoporin 75kD,Secretory 13 |
| Under-expressed 48 AEL | 1726 | Cellular component | U4/U6 x U5 tri-snRNP complex | GO:0046540 | 0.00098 | 5 | 9 | 19 | 2.91 | 47.4% | 0.5% | CG10418,CG2021,CG6610,CG9344,hoip,LSm3,SmB,SmD2,SmE | hoi-polloi,Like Sm 3,Small ribonucleoprotein particle protein SmB,Small ribonucleoprotein particle protein SmD2,Small ribonucleoprotein particle protein SmE,uncharacterized protein |
