## Supplementary Figures S1-S6 for "Innovation of heterochromatin functions drives rapid evolution of essential ZAD-ZNF genes in *Drosophila*"

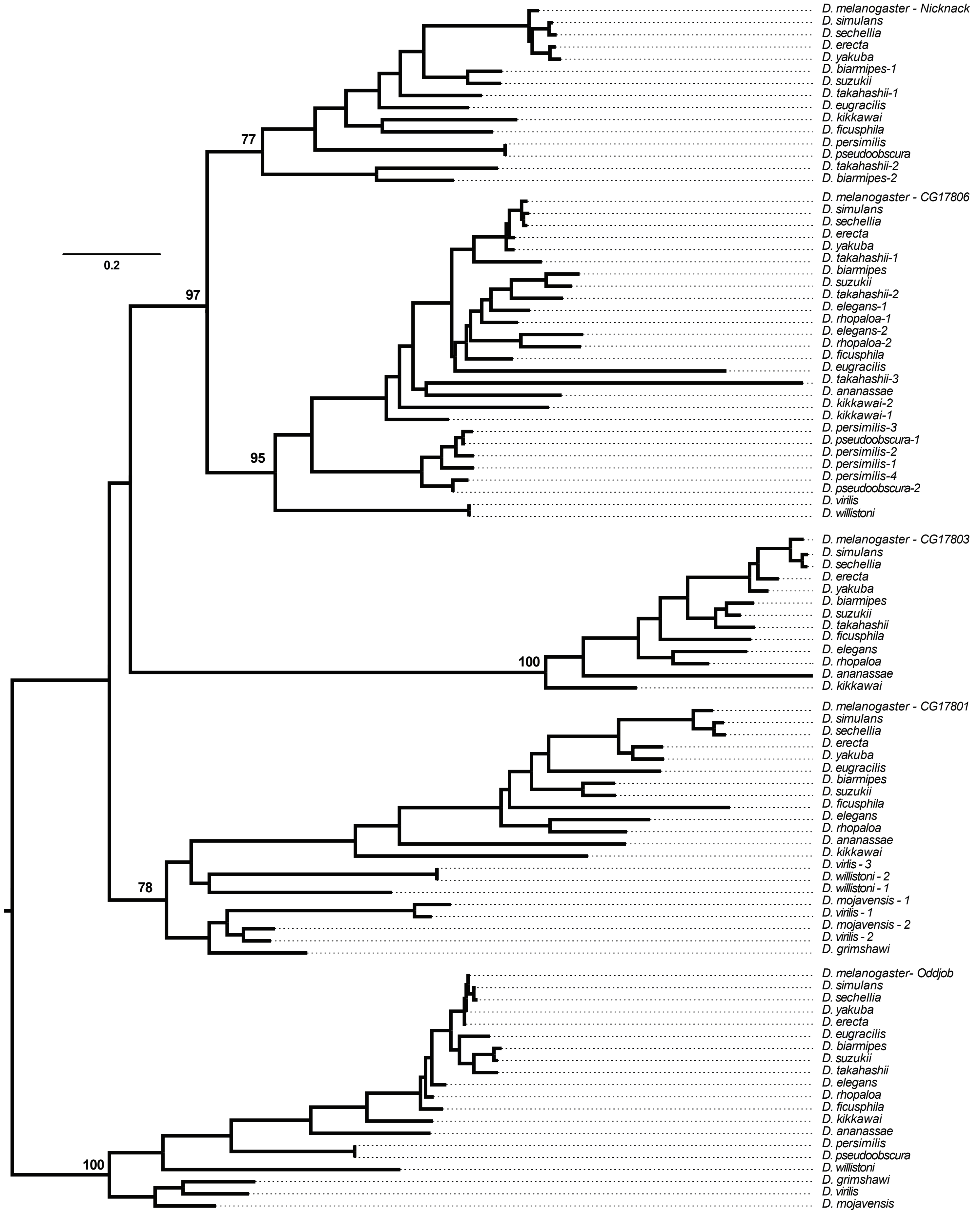


### Supplementary Figure S1

| Genotype | Rescue with transgene | Rescue | Heterozygous | Total | Replicates |
|---|---|---|---|---|---|
| *piggyBac* null / *piggyBac* null | yes | 64 (71) | 221 (214) | 285 | 3 |
| CRISPR null / CRISPR null | yes | 46 (44) | 130 (132) | 176 | 3 |
| $Nnk^{RNAi}$ / Act5C | yes | 169 (161) | 313 (321) | 482 | 9 |

### Supplementary Figure S2

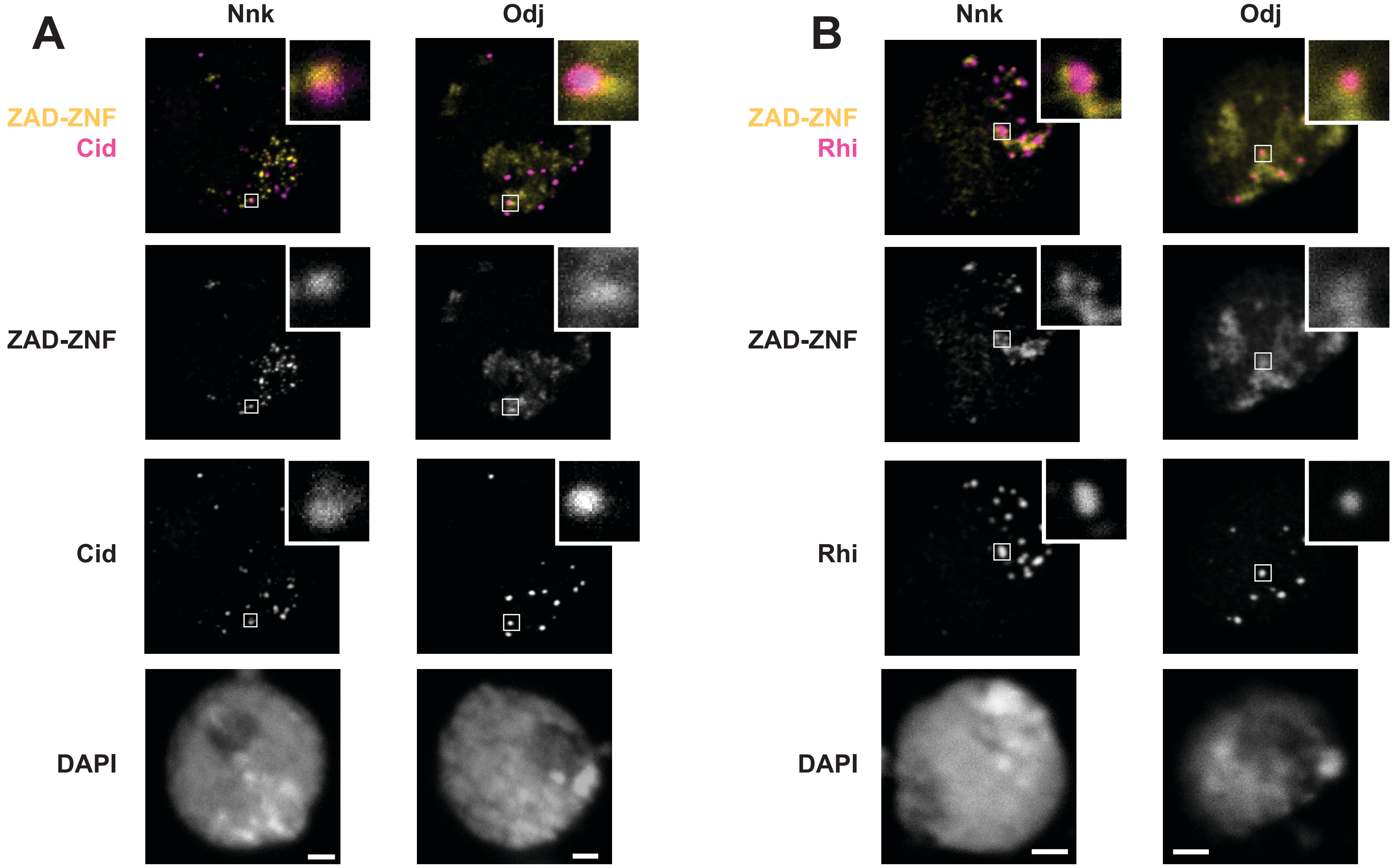


### Supplementary Figure S3

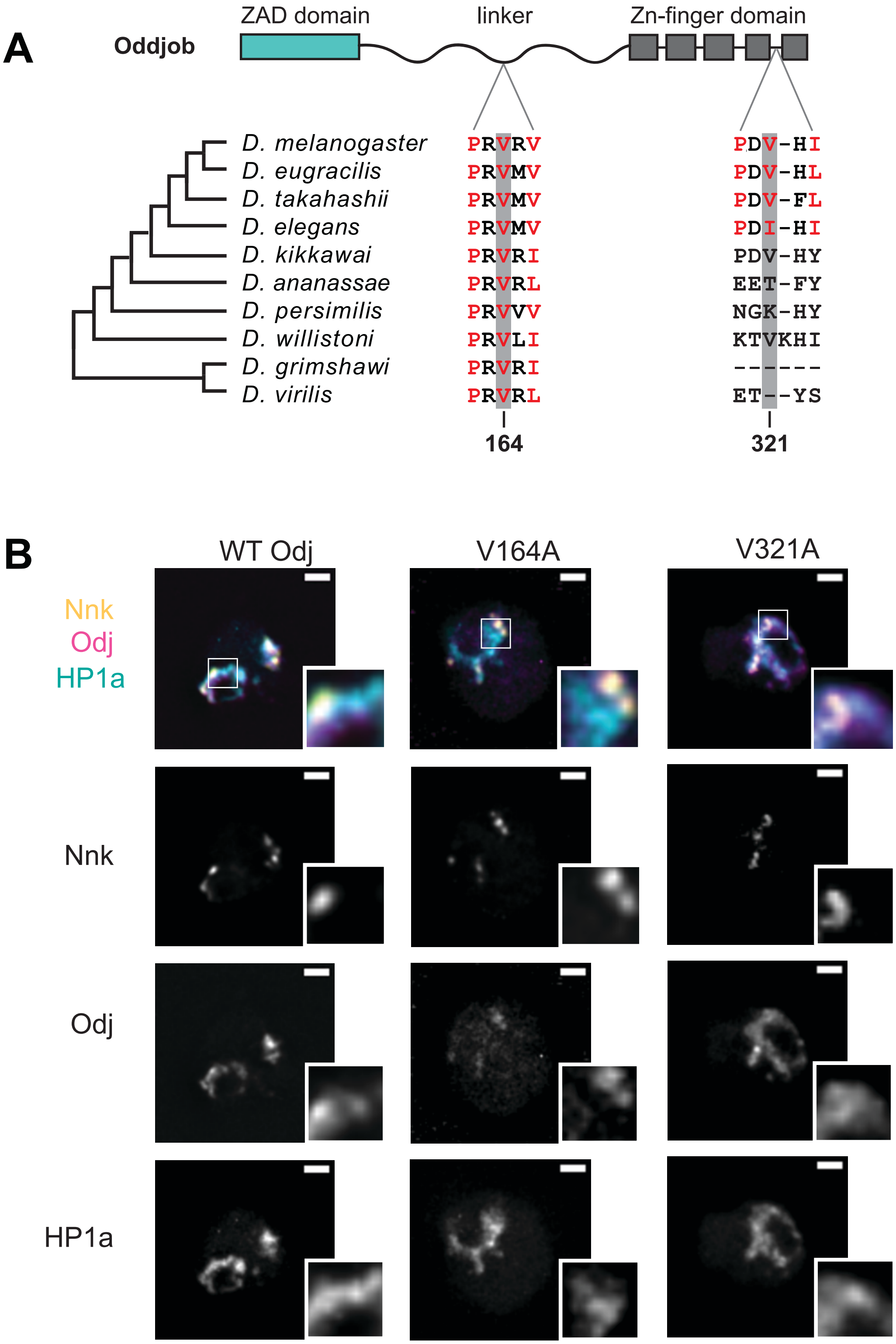


### Supplementary Figure S4

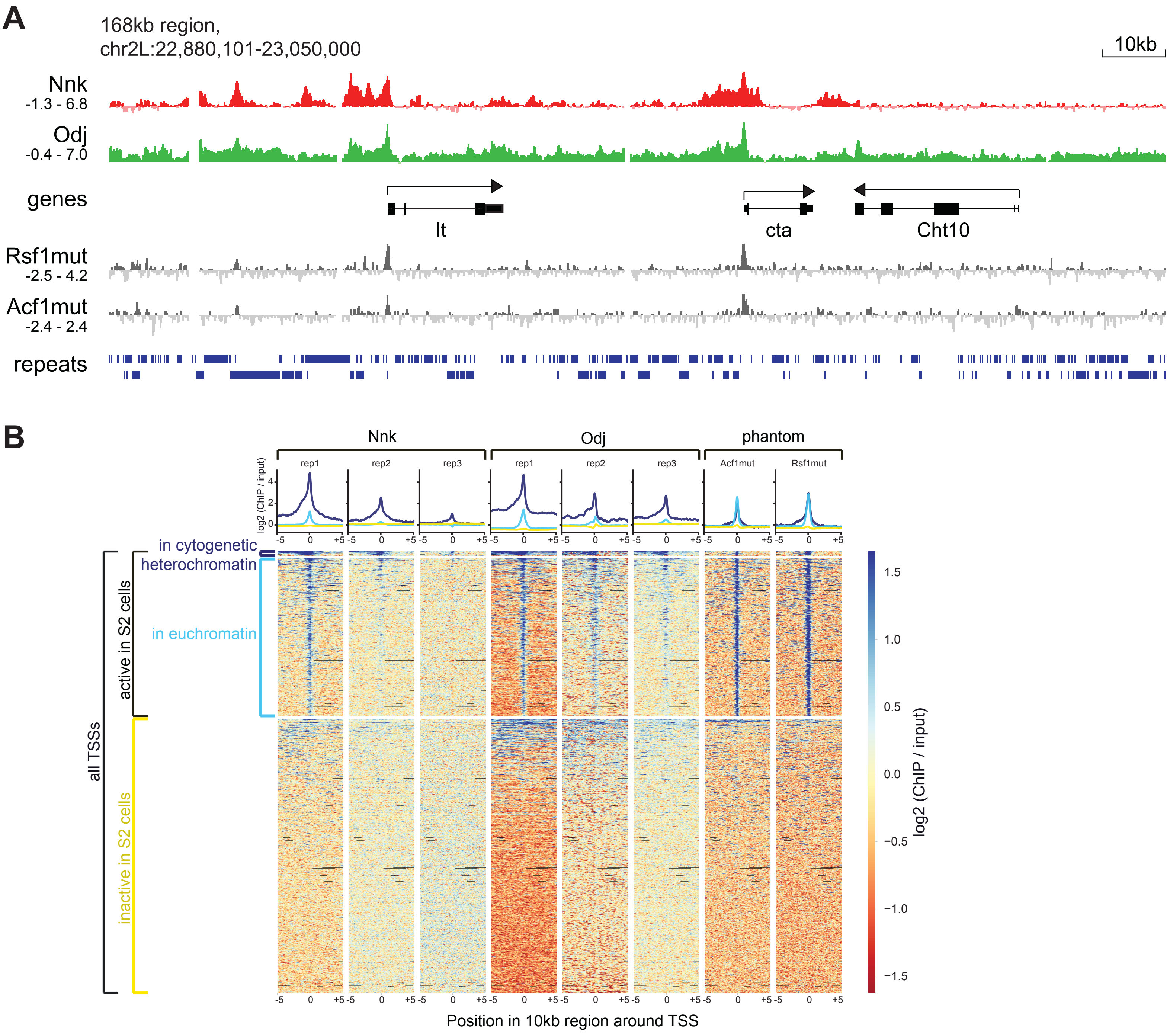


### Supplementary Figure S5

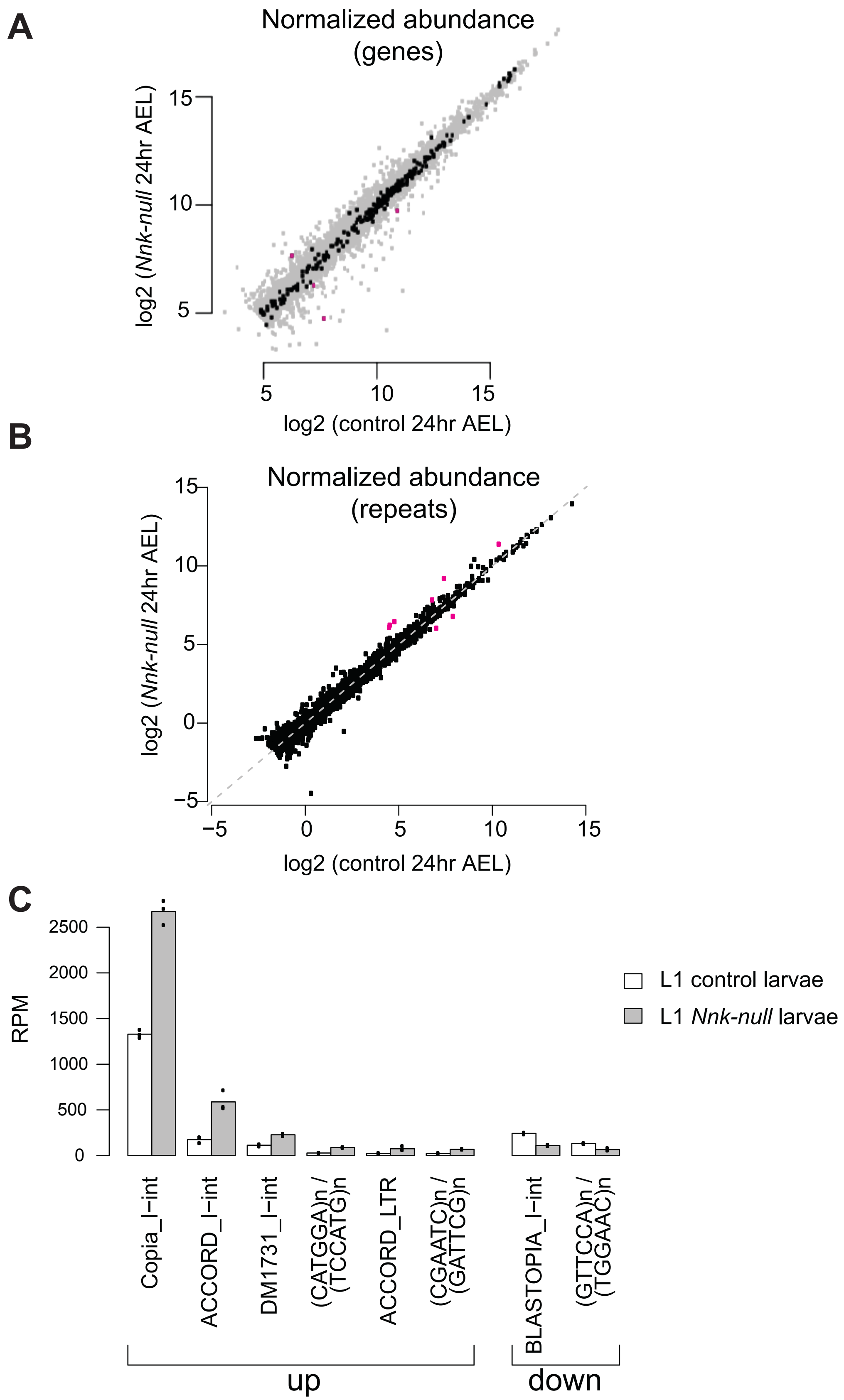


**Supplementary Figure S6**
